# A cancer immunotherapy modality based on dendritic cell reprogramming *in vivo*

**DOI:** 10.1101/2024.07.08.602356

**Authors:** Ervin Ascic, Fritiof Åkerström, Malavika Sreekumar Nair, André Rosa, Ilia Kurochkin, Olga Zimmermannova, Xavier Catena, Nadezhda Rotankova, Charlotte Veser, Michal Rudnik, Tommaso Ballocci, Tiffany Schärer, Xiaoli Huang, Maria de Rosa Torres, Emilie Renaud, Marta Velasco Santiago, Özcan Met, David Askmyr, Malin Lindstedt, Lennart Greiff, Laure-Anne Ligeon, Irina Agarkova, Inge Marie Svane, Cristiana F. Pires, Fábio F. Rosa, Carlos-Filipe Pereira

## Abstract

Immunotherapy leads to long-term survival of cancer patients, yet generalized success has been hampered by insufficient antigen presentation and exclusion of immunogenic cells from the tumor microenvironment. Here, we developed an approach to reprogram tumor cells *in vivo* by adenoviral delivery of the transcription factors PU.1, IRF8, and BATF3, which enabled them to present antigens as type 1 conventional dendritic cells. Reprogrammed tumor cells remodeled their tumor microenvironment, recruited, and expanded polyclonal cytotoxic T cells, induced complete tumor regressions, and established long-term systemic immunity in different mouse melanoma models. In human tumor spheroids and xenografts, reprogramming to immunogenic dendritic-like cells progressed independently of immunosuppression, which usually limits immunotherapy. Our study paves the way for first-in-human trials and other applications of immune cell reprogramming *in vivo*.

**One-Sentence Summary:** Reprogramming of tumor cells to cDC1-like cells *in vivo* elicits systemic and long-term antitumor immunity.

## Introduction

Cancer immunotherapies depend on the establishment of tumor antigen-specific T cell responses (*1*). T cells recognize tumor antigens presented on major histocompatibility complexes (MHC) of tumor cells and execute their effector function by cell killing and production of inflammatory cytokines (*2*, *3*). However, tumor cells often do not activate T cells due to downregulation of antigen presentation pathways, mounting an immunosuppressive tumor microenvironment (TME) and lack or dysfunction of professional antigen presenting cells (*1*). These include dendritic cells (DCs) that capture and present tumor antigens to T cells. Therefore, it has been challenging to achieve generalized success with current cancer immunotherapy modalities. For instance, immune checkpoint blockade (ICB), which has revolutionized the treatment of solid tumors, results in a 60% response rate in melanoma patients treated with anti–programmed cell death protein 1 (PD-1) and anti–cytotoxic T lymphocyte–associated protein 4 (CTLA-4) (*4*). Other low immunogenic cancer types are more refractory to immunotherapy including breast, microsatellite-stable colorectal cancer and glioblastoma where long-term immunity is only induced in <5% of patients (*5–7*). A growing body of evidence indicates that conventional dendritic cells type 1 (cDC1s) are required for T cell-mediated tumor regression and response to ICB across many cancer types (*8*– *11*). cDC1s are a rare subset of DCs that after maturation express high levels of MHC class I and II, the co-stimulatory molecule CD40, and the subset restricted markers XCR1 and CLEC9A (*12*). Within tumors, cDC1s have vital functions on the recruitment and activation of T cells by chemokine secretion and antigen cross-presentation (*13*), which mediate effective immunity against cancer (*14*). These unique functional properties of cDC1 are not yet deployed for immunotherapy.

Cellular reprogramming provides a strategy for generating individual cell types *in vivo* through enforced expression of transcription factor combinations (*15*). *In vivo* cell fate reprogramming enables the conversion of endogenous somatic cells into another cell identity within the organism for therapeutic benefit directly at the disease location. This strategy has the potential to overcome the substantial challenge of *ex vivo* cell manufacturing for personalized cell therapies. For instance, mouse pancreatic exocrine cells were shown to be converted *in situ* to insulin-secreting β-cells by delivering three transcription factors to the pancreas using adenoviral vectors (*16*). In mouse models of myocardial infarction, scar-forming cardiac fibroblasts were converted into cardiomyocytes leading to improved heart function (*17*). Glial cells were converted to functional neurons after brain injury or in models of neurodegenerative diseases (*18*), and rod photoreceptors were generated within the retina resulting in improved vision (*19*). *In vivo* reprogramming may however differ from the conversion process *in vitro*. Insulin-producing β-cells and cardiomyocytes were shown to acquire improved functional properties when generated *in vivo* due to the availability of biochemical and mechanical signals (*16*, *17*). In addition, the transcription factors Ngn2, Dlx2 or NeuroD1 were differentially employed to induce astrocyte-to-neuron conversion *in vitro* and *in vivo* (*20*). Differences in the transcription factor combination requirement and the maturity of the cells reported in these studies demonstrated that the *in vivo* environment has a significant impact on the reprogramming process, highlighting the need to characterize *in vivo* reprogramming mechanisms and induced phenotypes.

We previously identified the combination of transcription factors composed by PU.1, IRF8, and BATF3 (PIB) as sufficient to reprogram fibroblasts or tumor cells into cDC1-like cells *in vitro* endowed with the three signals required to activate T cells, including antigen presentation on MHC class I and II, co-stimulatory molecule expression and chemokine/cytokine secretion (*21–23*). In this study, we hypothesized that PIB mediate the reprogramming of tumor cells into immunogenic cDC1-like cells entirely *in vivo* within the TME. Our findings show that cDC1 reprogramming progresses *in situ* and leads to robust, long-lasting, and systemic antitumor immunity independently of exogenous stimulation, providing a tractable strategy to induce antigen presentation and cDC1 function *in vivo* and set in motion tumor antigen-specific immune responses.

## Results

### Systemic and durable antitumor immunity induced by cDC1 reprogramming *in vivo*

To evaluate the feasibility of cDC1 reprogramming *in situ* as a cancer immunotherapeutic modality, we first assessed whether the PIB transcription factors were sufficient to drive *in vivo* reprogramming of tumor cells to immunogenic cDC1-like cells within the TME without relying on artificial antigens or exogenous stimulation and characterized induced immune mechanisms. We then evaluated the reprogramming of human cancer cells in spheroids and in xenografts and identified a viral vector to deliver the transcription factors to tumors as a gene therapy approach based on *in situ* cDC1 reprogramming (Fig. 1A).

**Figure 1.**
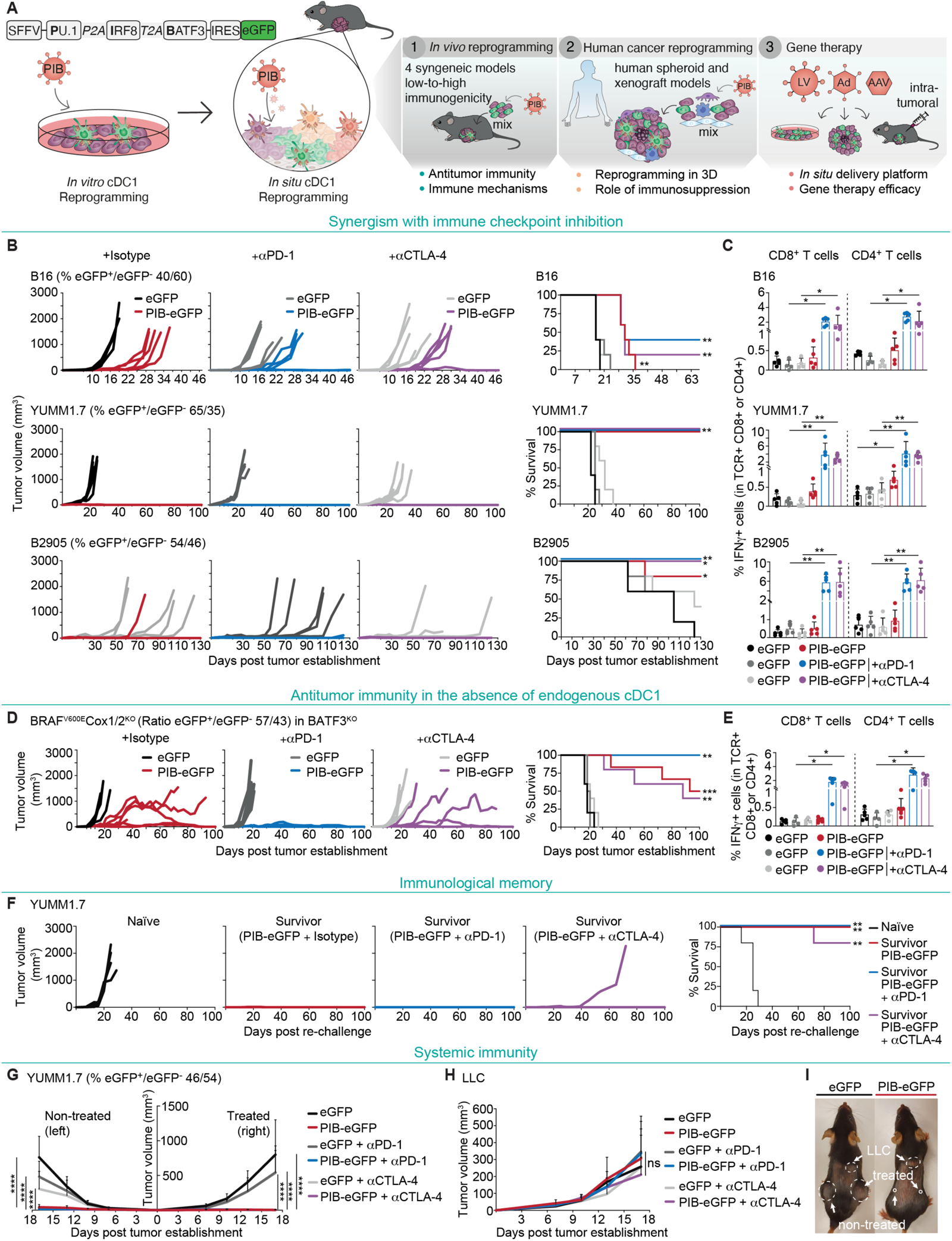
*In vivo* cDC1 reprogramming elicits systemic and durable antitumor immunity. **(A)** Experimental strategy to induce cDC1 reprogramming *in vivo* employing a polycistronic lentiviral vector encoding the transcription factors PU.1, IRF8, and BATF3 (PIB) followed by IRES-eGFP. First, *in vivo* cDC1 reprogramming was tested by implantation of a mixture of transduced cancer cells and untransduced parental cells to assess antitumor immunity. Secondly, human cancer cells were reprogrammed in spheroids with immunosuppressive cells and in xenografts. Third, lentiviral (LV), adenoviral (Ad), and adenoviral-associated (AAV) vectors were tested to deliver PIB to tumors *in situ* as a cancer gene therapy. **(B)** C57BL/6J mice were injected subcutaneously with melanoma cells (B16, YUMM1.7, B2905) after transduction with PIB-eGFP or control eGFP and mixing 1:1 with parental cells (measured percentages by flow cytometry at day 3 are indicated) to induce tumor cell reprogramming *in vivo* along with tumor establishment. Anti-PD-1, anti-CTLA-4 or isotype control antibodies were administered by intraperitoneal injection at days 7, 10 and 13. Tumor growth and survival are shown (n=5). **(C)** Flow cytometry quantification of tumor antigen-specific IFNγ^+^CD8^+^ or IFNγ^+^CD4^+^ T cells from peripheral blood at day 14. T cells were isolated and re-stimulated *in vitro* using an antigen-agnostic approach with IFNγ-stimulated melanoma cell lines. **(D)** Tumor growth and survival of BATF3^KO^ mice after injection with PIB-eGFP- or eGFP-transduced BRAF^V600E^COX1/2^KO^ melanoma cells (n=5-6). **(E)** Quantification of tumor-antigen specific IFNγ^+^CD8^+^ or IFNγ^+^CD4^+^ T cells from peripheral blood with the antigen-agnostic approach applied in (C). **(F)** Survivor C57BL/6J mice that remained tumor-free for 100 days were re-challenged with YUMM1.7 cells. Age-matched naïve mice were used as controls and tumor growth and survival are shown (n=5). **(G)** Bilateral YUMM1.7 tumor growth after injection of 1:1 mixtures into the treated flank (right) and untransduced cells into the non-treated flank (left), as monotherapy (PIB-eGFP) or in combination with anti-PD-1 or anti-CTLA-4 (n=10). **(H)** Control Lewis lung adenocarcinoma (LLC) tumor growth within the same animals. **(I)** Representative pictures of animals with bilateral YUMM1.7 (treated and non-treated) and LLC tumors. Arrows indicate tumor locations and dashed lines tumor sizes. Data in panels C, E, G and H are shown as mean ± SD. Survival analyses in panel B, D and F were performed by log-rank Mantel-Cox test. Comparisons in panels C, E, G and H were analyzed using the Mann-Whitney test. ns - non-significant, *p<0.05, **p<0.01, ***p<0.001, ****p<0.0001.

First, we aimed at verifying reprogrammed cells’ capacity to drive systemic immunity in the absence of toll-like receptor (TLR) 3 stimulation *in vitro* (*23*), which would be critical for the success of a local immunotherapy approach mediated solely by the expression of PIB reprogramming factors. We employed a low immunogenic murine melanoma cell line B16-F10 (B16), characterized by low MHC expression and resistance to ICB treatment and the immunogenic line B2905, which models highly mutated melanoma tumors (*23*, *24*). First, to evaluate systemic immunity we established subcutaneous B16 tumors expressing the model antigen ovalbumin (OVA) bilaterally and a heterologous Lewis lung adenocarcinoma (LLC) tumor in the same wild-type (WT) C57BL/6J animals. Then, we injected *in vitro* reprogrammed B16-derived cells pulsed with OVA and the TLR3-agonist polyinosinic-polycytidylic acid (P(I:C)) into the right flank tumors in combination with systemic anti-PD-1 and anti-CTLA-4 administration. Reprogramming into cDC1-like cells was induced using a lentiviral polycistronic vector encoding for PIB followed by an internal ribosomal entry site (IRES) and enhanced green fluorescent protein (eGFP) to track transduced cells (Fig. 1A) (*23*). eGFP-transduced cells were used as controls for lentiviral-mediated immunogenicity. Interestingly, both B16-OVA tumors showed a clear reduction in tumor growth (fig. S1A), but not LLC tumors, demonstrating systemic and antigen-specific antitumor immunity (fig. S1B, C). Next, we asked whether induced immunity is dependent on P(I:C) stimulation and the artificial OVA antigen. Therefore, we established B2905 tumors and administered P(I:C)-stimulated or unstimulated B2905-derived reprogrammed cells. Both resulted in delayed tumor growth and similar medium survival (MS) (fig. S1D). These data illustrate the capacity of cDC1 reprogramming to induce antitumor immunity independently of exogenous stimulation and the presence of highly immunogenic model antigens.

To test the anti-tumor efficacy of *in vivo* cDC1 reprogramming we subcutaneously implanted a mixture of 88% PIB-eGFP-transduced B16 cells and 12% untransduced parental cells, or mixtures of eGFP-transduced and parental cells as a control, 16 hours after transduction (fig. S2A, B). This strategy allowed to separate the delivery of transcription factors from the *in vivo* reprogramming process. We observed complete responses (CR) in 30% of animals and delayed tumor growth in the other animals, thereby extending MS (43 vs. 19 days, p<0.0001). Interestingly, we detected vitiligo at the tumor regression site (fig. S2C), demonstrating the induction of a cytotoxic response against melanoma antigens (*23*). When combined with anti-PD-1 and anti-CTLA-4, we observed tumor regression in all animals (fig. S2B). We confirmed efficient delivery of the transcription factors to tumor cells and the absence of phenotypic reprogramming before implantation *in vivo* (fig. S2D-F). To dissect whether cDC1’ functional properties are critical for the observed potent antitumor immunity we compared cDC1 reprogramming with myeloid reprogramming mediated by PU.1 and C/EBPα to induce macrophage-like cells (*25*) (fig. S2A-F). *In vivo*, cDC1 reprogramming extended MS when compared to macrophage reprogramming (43 vs. 29.5 days, p<0.0001), especially when combined with ICB (p=0.0003), which resulted in 100% CRs (fig. S2B). This effect is consistent with the selective induction of high levels of MHC-I and MHC-II (fig. S2G, H) and cross-presentation capacity by PIB (fig. S2I). We next confirmed that *in vivo* cDC1 reprogramming combined with ICB induced systemic immunity by performing bilateral tumor challenges (fig. S3A, B). This resulted in tumor reduction of both treated and non-treated tumors and systemic expansion of cytotoxic T cells in peripheral blood recognizing the melanoma tumor antigens PMEL, TRP-2, and p15E (fig. S3A-C).

Next, we investigated *in vivo* cDC1 reprogramming as monotherapy or in combination with either anti-PD-1 or anti-CTLA-4 using B16, B2905 and the additional melanoma model YUMM1.7, which is resistant to ICB and also depends on cDC1 availability (*11*, *24*, *26*). We implanted a 1:1 mixture of transduced and parental melanoma cells (fig. S3D, E) and observed that monotherapy induced complete tumor regressions in YUMM1.7 (100% CR), B2905 (80% CR), and extended MS in B16 challenged animals from 17 to 31 days (Fig. 1B). cDC1 reprogramming synergized with anti-PD-1 or anti-CTLA-4 treatment leading to increased CRs in B16 and B2905, which also resulted in expansion of tumor antigen-specific IFNγ^+^CD8^+^ and IFNγ^+^CD4^+^ T cells in peripheral blood (Fig. 1C). To assess whether antitumor immunity requires endogenous cDC1, we used the immunogenic BRAF^V600E^COX1/2^KO^ melanoma model which grows in BATF3^KO^ mice due to the lack of endogenous cDC1 (*13*). We observed increased MS (96.5 vs. 19.5 days, p<0.001) (Fig. 1D), synergy with anti-PD-1 treatment (50% vs. 100% CR), and concomitant expansion of tumor antigen-specific T cells (Fig. 1E).

To address whether immune memory was induced, we re-challenged survivor WT or BATF3^KO^ animals that showed complete tumor regressions. While naïve mice developed tumors, survivor animals remained tumor-free (100% YUMM1.7; 66% BRAF^V600E^COX1/2^KO^) (Fig. 1F and fig. S3F). We then asked whether combination with ICB is required for systemic antitumor immunity and observed tumor-specific abscopal effects with the monotherapy or when combined with anti-PD-1 or anti-CTLA-4 (Fig. 1G-I). Taken together, these findings highlight that *in vivo* cDC1 reprogramming mediated by PU.1, IRF8 and BATF3 is (i) sufficient to elicit antitumor immunity, (ii) protects from distal tumor growth and (iii) tumor growth after re-challenge, and (iv) the effects are independent of endogenous cDC1s. Interestingly, both cDC1 and myeloid *in vivo* reprogramming systems elicited antitumor immunity, illustrating the potential of cellular reprogramming *in vivo* as a new modality for cancer immunotherapy.

### Remodeling of the tumor microenvironment

Given the ability of cDC1 to shape the TME (*9*, *27*), we addressed the impact of *in vivo* reprogramming on tumor morphology and immune composition by immunofluorescence and flow cytometry. At day 9 after initiation of *in vivo* reprogramming, we observed a global increase of immune cell (CD45^+^) infiltration and a reduction of transduced tumor cells (fig. S4A). Strikingly, we observed that *in vivo* reprogramming led to the formation of dense lymphocyte clusters with defined borders resembling tertiary lymphoid structures (TLS) in the parenchyma of tumors (Fig. 2A), which contained a B cell and CD4^+^ T cell zone and a spatially segregated CD8^+^ T cell zone that were juxtaposed with TLS-specific podoplanin^+^ stromal cells (Fig. 2B). At day 21, reprogrammed tumors were smaller and showed a 2.7- and 1.5-fold increase in CD45^+^ cell infiltration as monotherapy or when combined with anti-PD-1, respectively (Fig. 2C, D). Within the lymphoid compartment, B cell percentages increased by 24.4-fold, NK cells by 2.2-fold, and CD4^+^ T cells by 2.4-fold (Fig. 2E). Although the percentages of CD8^+^ T cells were similar between treated and untreated tumors, we found a substantial decrease of exhausted PD-1^+^CD8^+^ and PD-1^+^CD4^+^ T cells, (4- and 8-fold, respectively) (Fig. 2F). Conversely, we observed a 4.3-fold increase of central memory CD62L^+^CD44^+^CD8^+^ T cells, that are critical for long-term memory (*28*), and a 2.2-fold increase in effector CD44^+^CD4^+^ T cells (Fig. 2G). Interestingly, we observed a 2.3-fold increase in the percentages of proliferative Ki-67^+^CD8^+^ T cells, as well as a 1.7-fold increase in TCF-1^+^CD8^+^ and 2.5-fold increase in TCF-1^+^CD4^+^ T cells (Fig. 2H, I), which have been attributed a function in the persistent control of tumor growth (*13*). We observed a 10.5- and 1.5-fold decrease in regulatory CD8^+^ and CD4^+^ T cell (Treg) populations, indicating that *in vivo* cDC1 reprogramming within tumors induces a shift towards pro-inflammatory T cell populations (Fig. 2J, K). In the myeloid compartment, reprogramming increased XCR1^+^ cDC1s by 6.9-fold, SIRPα^+^ cDC2s by 2.1-fold, Siglec-H^+^ pDCs by 3.5-fold, Ly6C^+^ monocytes by 2-fold and decreased F4/80^+^ macrophages by 1.4-fold (fig. S4B). Interestingly, we observed higher expression of PD-L1 in myeloid cells, including dendritic cells, macrophages, and neutrophils, reflecting inflammation (*29*) (fig. S4C). Next, we profiled lymphoid populations in tumor-draining lymph nodes (tdLN) and observed expansion of CD4^+^ T cells (eGFP 23.6±2.3% vs. PIB-eGFP 29.0±4.9%), while no major changes in CD8^+^ T, B and NK cells were observed (Fig. 2L and fig. S4D). Regarding the functional state of induced cDC1-like cells, we could not detect their presence in lymph nodes (fig. S4E), expression of CCR7 (fig. S4F) or activation of migratory state-associated gene expression (fig. S4G). Instead, we detected activation of a resident cDC1 program (14) further evidenced by their persistence in the tumor for at least 9 days (fig. S4E, G).

**Figure 2.**
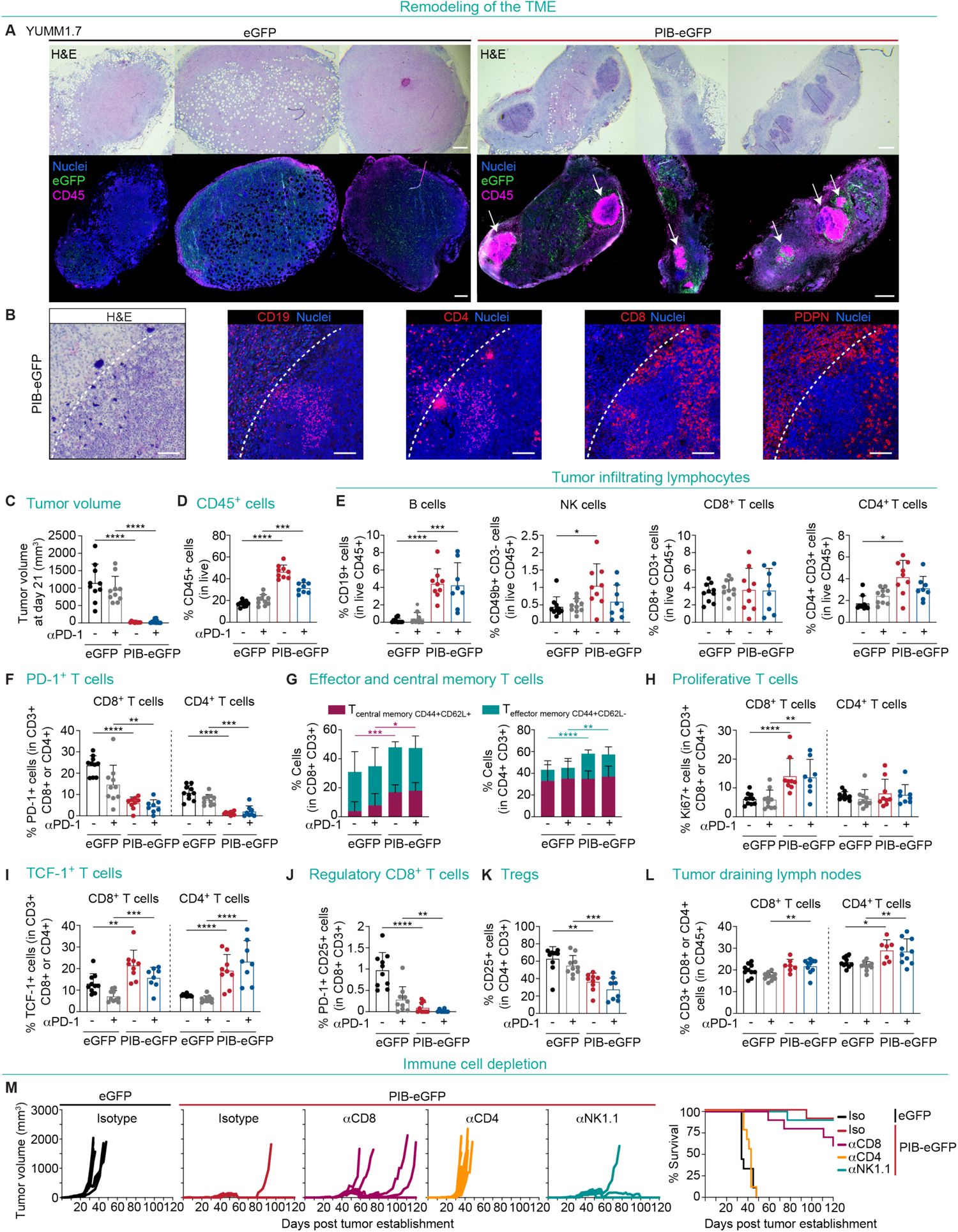
cDC1 reprogramming remodels the tumor microenvironment. **(A)** Hematoxylin and eosin (H&E) staining (top) and immunofluorescence (bottom) analysis of paraffin-embedded YUMM1.7 tumors 9 days after subcutaneous implantation of PIB-eGFP- or control eGFP-transduced cells (1:2 ratio of transduced to parental cells). Tumor sections were stained for eGFP (green, transduced cells), CD45 (purple, immune cells) and nuclei (blue, Syto 13) (n=3). Arrows indicate TLS-like structures. Scale bars are 500nm. **(B)** H&E (left) and immunofluorescence (right) of a TLS in PIB-eGFP tumors stained for CD19 (B cells), CD4 (CD4^+^ T cells), CD8 (CD8^+^ T cells) and PDPN (podoplanin^+^ stromal cells). Dashed lines indicate TLS border. Scale bars are 100µm. **(C)** Volumes of YUMM1.7 tumors 21 days after establishment and treatment with anti-PD-1 (grey and blue) or isotype control (black and red) antibodies at days 7, 10, and 13 (n=8-10). **(D)** Flow cytometry quantification of the percentages of tumor-infiltrating CD45^+^ cells and **(E)** CD19^+^ B cells, CD49b^+^CD3^-^ NK cells and CD8^+^ and CD4^+^ T cells. **(F)** Quantification of PD-1^+^CD8^+^ and PD-1^+^CD4^+^ T cells. **(G)** Percentages of CD44^+^CD62L^-^ effector memory and CD44^+^CD62L^+^ central memory CD8^+^ and CD4^+^ T cells. **(H)** Quantification of Ki67^+^ proliferative, **(I)** TCF-1^+^CD8^+^ and TCF-1^+^CD4^+^ T cells, **(J)** PD-1^+^CD25^+^ regulatory CD8^+^ T cells, **(K)** CD25^+^CD4^+^ Tregs. **(L)** Percentages of CD8^+^ and CD4^+^ T cells in the tumor-draining lymph nodes. **(M)** Mice were subjected to antibody-mediated depletion of CD8^+^ T cells (αCD8), CD4^+^ T cells (αCD4), NK cells (αNK1.1) or isotype controls and tumors established with a mixture of transduced YUMM1.7 cells. Tumor growth (left) and survival (right) are shown (n=10). Data in panel C-L are shown as mean ± SD. Comparisons in panel C-L were analyzed using the Mann-Whitney test. *p<0.05, **p<0.01, ***p<0.001, ****p<0.0001.

Global changes in immune composition prompted us to functionally investigate the contribution of individual effector cell populations for tumor control. We first confirmed that lymphocytes are essential for reprogramming-mediated tumor control by eliciting *in vivo* reprogramming in immunodeficient NOD.Cg-Prkdc^SCID^ Il2rg^tm1Wjl^/SzJ (NSG) animals (fig. S4H). Mice did not survive beyond day 30 post implantation despite showing extended MS when compared to eGFP control (day 18, p=0.01). This can be attributed to the reduced proliferation capacity of reprogrammed cells (fig. S4I) (*23*). Next, we depleted CD8^+^ T cells, CD4^+^ T cells and NK cells with antibodies in WT mice (fig. S4J) and observed that depletion of CD4^+^ T cells abolished tumor immune control, highlighting the critical role of CD4^+^ T cells in PIB-mediated antitumor immunity (Fig. 2M). While NK cell depletion did not show an impact in tumor growth, CD8^+^ T cell depletion reduced tumor control at later time points in 40% of animals (Fig. 2M). In summary, our data show that *in vivo* cDC1 reprogramming remodels the TME, induces TLS-like structures, reduces exhausted and regulatory populations and increases the infiltration of memory and stem-like T cells.

### Induction of polyclonal cytotoxic and memory T cell responses

To further characterize T cell responses elicited by *in vivo* cDC1 reprogramming, we profiled T cells from tumors, tdLN and peripheral blood using single cell RNA-sequencing (scRNA-seq) with T cell receptor (TCR) enrichment (Fig. 3A, B and fig. S5A). We segregated single cells expressing either *CD8a* or *CD4* (fig. S5B) and performed cluster annotation (fig. S5C). We identified 9 CD8^+^ T cell clusters (Fig. 3C, fig S5C, D and data file S1) and observed increased frequencies of intratumoral effector and effector memory cells in PIB-treated tumors, accompanied by a reduction in exhausted and terminally exhausted subsets (Fig. 3C). Reduced CD8^+^ T cell exhaustion was confirmed by decreased PD-1 expression (*Pdcd1,* fig. S5D), TIM-3 (*Havcr2)* and *Cd101* in the monotherapy setting as well as an additional decrease in *Lag3* expression when combined with anti-PD-1 (fig. S5E). Indeed, the frequencies of effector and stem-like CD8^+^ T cells were amplified when *in vivo* reprogramming was combined with anti-PD-1 treatment. These differences led us to ask whether CD8^+^ T cells follow different differentiation paths when exposed to reprogrammed cells. In control eGFP tumors, trajectory analysis showed that early activated and proliferating CD8^+^ T cells mainly differentiated towards effector, exhausted and terminally exhausted cells (Fig. 3D). Anti-PD-1 treatment also enhanced differentiation towards exhausted and terminally exhausted populations as previously reported (*30*), but *in vivo* reprogramming favored effector and memory fates resulting in fewer exhausted and terminally exhausted T cells (Fig. 3D). We also identified 11 CD4^+^ T cell clusters (Fig. 3E and fig S5F, G). In tumors treated with *in vivo* reprogramming, we observed a large cluster of cytotoxic CD4^+^ T cells that has been described to harbor the capacity to directly eradicate melanoma cells (*3*) (Fig. 3E). Interestingly, when combined with anti-PD-1, CD4^+^ T cells shifted towards T helper (Th) precursors, type 1 T helper (Th1), and stem-like fates indicating either differential kinetics or differential priming of these populations. We also observed decreased frequencies of exhausted and regulatory CD4^+^ T cells. Trajectory analysis showed that CD4^+^ T cells exposed to reprogrammed cells progressed mainly through a Th precursor to effector cytotoxic and Th1 fates (Fig. 3F).

**Figure 3.**
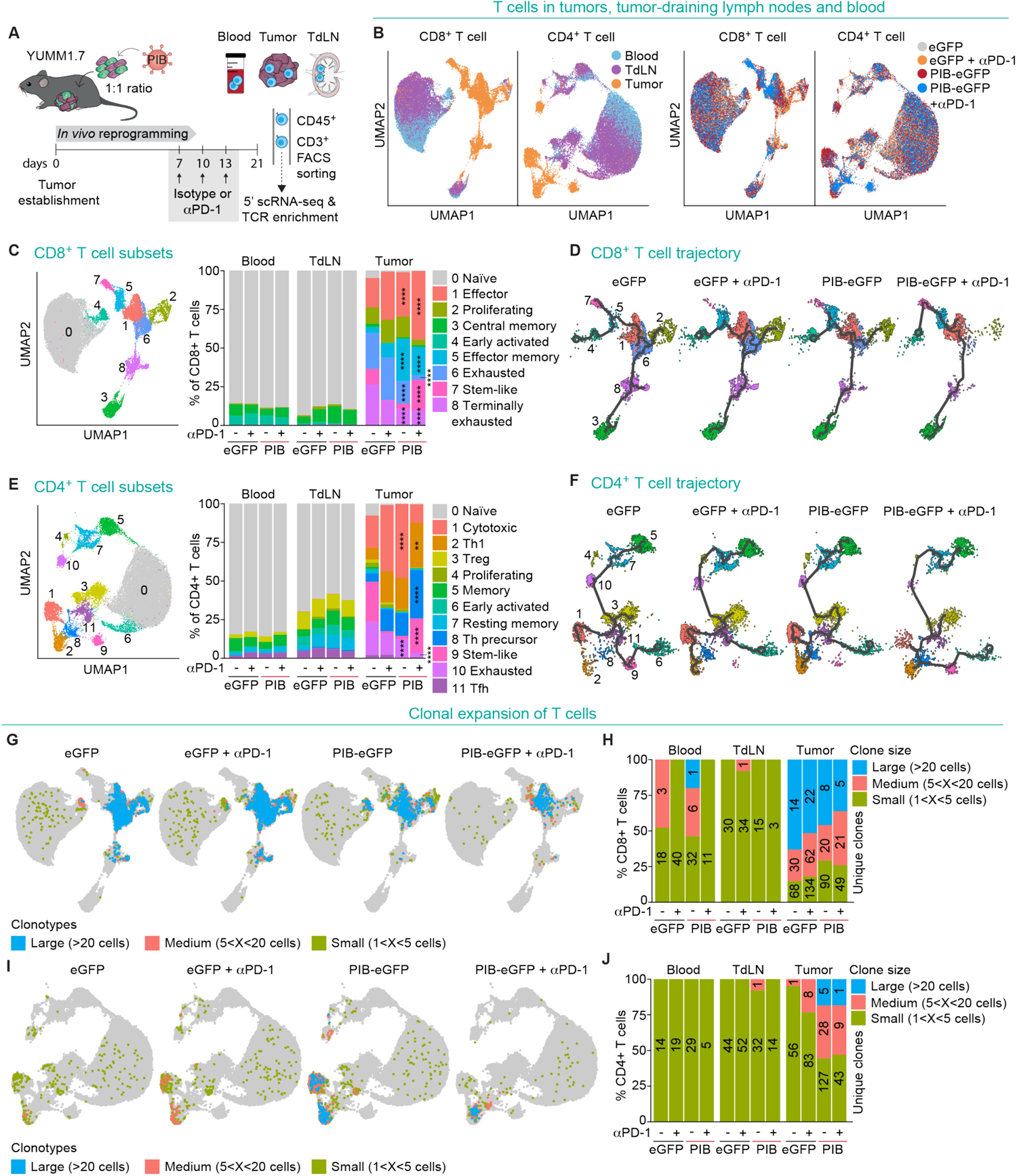
*In vivo* reprogrammed melanoma cells expand polyclonal CD4^+^ T cells. **(A)** Experimental design for 5’ single cell RNA-seq with TCR enrichment. YUMM1.7 tumors were established with a 1:1 mixture of PIB-eGFP or control eGFP-transduced and untransduced cells. Peripheral blood, tumor-draining lymph nodes (tdLN) and tumors were isolated 21 days after tumor establishment and CD45^+^CD3^+^ T cells were FACS-purified before loading on a 10x Chromium. Additional groups received intraperitoneally anti-PD-1 at days 7, 10 and 13 (n=5). **(B)** Principal component analysis of CD8^+^ and CD4^+^ T cells visualized by Uniform manifold approximation and projection (UMAP) plots from tumors, tdLN and blood (left) across treatment conditions (right). **(C)** UMAP plot showing color-coded CD8^+^ T cell subsets (left). Bar plots show the percentages of each CD8^+^ T cell subset in blood, tdLN and tumors (right). CD8^+^ T cell subsets are numbered from 0 to 8. **(D)** Trajectory analysis (black line) of CD8^+^ T cells across treatment conditions. **(E)** UMAP plot showing color-coded CD4^+^ T cell subsets (left). Bar plots (right) show the percentages of each CD4^+^ T cell subset in blood, tdLN and tumors. CD4^+^ T cell subsets are numbered from 0 to 11. **(F)** Trajectory analysis (black line) of CD4^+^ T cells across treatment conditions. **(G)** CD8^+^ T cells isolated from tumors, tdLN and blood were color-coded by clonotype size into small (between 1 and 5 cells), medium (between 5 and 20 cells), and large (>20 cells) clones and projected onto UMAP plots across treatment conditions. TCR sequences detected in only one single cell were excluded from this analysis. **(H)** Bar plots showing percentages of CD8^+^ T cells in blood, tdLN and their clonotype distribution. The numbers of unique clones are indicated within the bars. **(I)** Tumor, tdLN and blood-derived CD4^+^ T cell clonotype sizes projected onto UMAP plots and **(J)** bar plots showing percentages of CD4^+^ T cells and their clonotype distribution. Comparisons in C and E were performed using the exact Binomial test. Relevant statistical comparisons between intratumoral T cells for the conditions eGFP vs. PIB-eGFP and eGFP+anti-PD-1 vs. PIB-eGFP+anti-PD-1 are shown. All statistical comparisons can be found in data file S1. **p<0.01, ****p<0.0001.

To address if polyclonal expansion was induced, we analyzed full-length αβ TCR sequences obtained from single T cells with more than 1 cell per clonotype. First, we observed similar polyclonal expansion of CD8^+^ T cells in tumors treated with *in vivo* reprogramming and control tumors (118 and 112 unique clones, respectively) (Fig. 3G, H). Expanded clones in PIB-treated tumors when compared to eGFP tumors were primarily within the effector (44.4% vs. 30.1%) and effector memory CD8^+^ T cell subsets (4.8% vs. 0.1%) (fig. S6A, B). Interestingly, we observed the expansion of 160 unique CD4^+^ T cell clones while only 57 and 92 were found in control tumors and tumors treated with anti-PD-1 (Fig. 3I, J). In PIB-treated tumors, 59.6% of expanded clones were cytotoxic and 32.9% were Th1 CD4^+^ T cells (fig. S6C, D). In contrast, the combination with anti-PD-1 treatment reduced CD4^+^ T cell clonal expansion (53 unique clones). To validate the cytotoxic potential of CD4^+^ T cells from tumors undergoing *in vivo* reprogramming, we performed killing assays *in vitro* and observed efficient and specific killing of melanoma cells (fig. S6E-G). In the blood, we observed expansion of 29 CD4^+^ T cell clones when compared to 14 and 19 in control mice and anti-PD-1 (Fig. 3I, J). Interestingly, the majority of these systemically circulating clones were identified as T follicular helper (Tfh) cells, a subset of CD4^+^ T cells supporting B cell responses, germinal center and TLS formation (fig. S6C) (*27*). Together, these results further underscore the importance of polyclonal CD4^+^ T cells for tumor control corroborating with immune depletion experiments (Fig. 2M).

### Induction of cDC1 phenotype in human tumors *in vivo*

The signals from the environment can impact the reprogramming process (*16*, *17*), and thus, we assessed whether human cancer cells acquire an immunogenic cDC1 phenotype *in vivo*. We established human melanoma (A375 and A2058) and glioblastoma (T98G) tumors in NSG mice and performed phenotypic characterization of the *in vivo* reprogramming process by flow cytometry (Fig. 4A and fig. S7A). At day 9, we detected reprogrammed CD45^+^HLA-DR^+^ T98G (75.36±11.84%), A375 (69.93±11.45%) and A2058 (76.97±7.24%) cells, as well as partially reprogrammed cells expressing either CD45 or HLA-DR (Fig. 4B, C and fig. S7B). We observed a gradual increase in the percentage of reprogrammed cells from days 3 to 9, showing not only the initiation but also progression of the reprogramming process *in vivo* as observed *in vitro* (*23*). As a measure of reprogramming fidelity, we assessed the expression of the cDC1-specific surface markers XCR1, CLEC9A, and CD226 (Fig. 4D and fig. S7C) that were previously identified as late markers of successful cDC1 reprogramming (*22*, *23*). Interestingly, already at day 5 we observed enhanced expression of XCR1 *in vivo* when compared to *in vitro* in T98G (8.0±2.5% vs. 1.2±1.1%), A375 (12.1±6.6% vs. 0.4±0.2%) and A2058-derived cells (9.5±6.5% vs. 4.8±3.2%) (Fig. 4D and fig. S7C). Regarding the immunogenicity of induced cells *in vivo* at day 9, we detected a 4-, 2-, and 3-fold higher expression of HLA class I molecules in reprogrammed T98G, A375 and A2058 cells, respectively (Fig. 4E). Moreover, we confirmed the acquisition of the co-stimulatory molecule CD40 reflecting a mature antigen-presenting phenotype (Fig. 4F). Overall, we demonstrated that PIB overexpression in human xenograft models induces a cDC1 phenotype *in vivo* with enhanced fidelity and immunogenicity.

**Figure 4.**
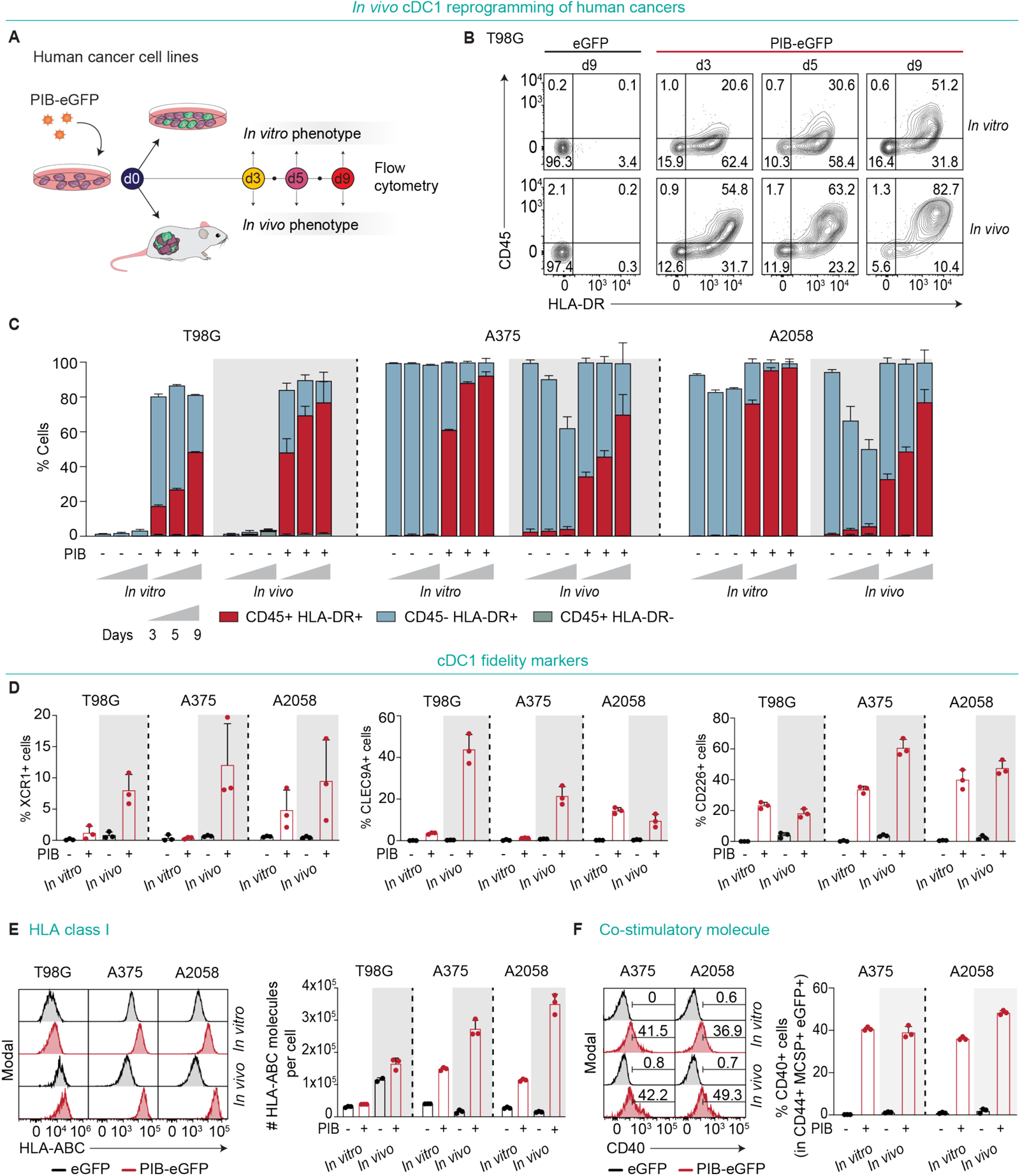
Induction of a cDC1 phenotype in human cancer cells *in vivo.* **(A)** Experimental design to assess phenotypic cDC1 reprogramming *in vivo*. Human cancer cell lines were transduced *in vitro* with PIB-eGFP, implanted in NSG mice and isolated at days 3, 5, and 9 for phenotypic profiling by flow cytometry. eGFP-transduced cells were used as controls and *in vitro* reprogrammed cells for comparison. Reprogramming efficiency was evaluated by flow cytometry as the percentage of CD45^+^HLA-DR^+^ cells (completely reprogrammed) and CD45^+^HLA-DR^-^ or CD45^-^HLA-DR^+^ cells (partially reprogrammed) gated in eGFP^+^ transduced cancer cells. **(B)** Representative flow cytometry plots and **(C)** quantification of reprogramming kinetics *in vitro* and *in vivo* of the glioblastoma cell line T98G (gated in CD44^+^eGFP^+^ cells), and melanoma lines A375 and A2058 cells (gated in CD44^+^MCSP^+^eGFP^+^ cells) (n=3). **(D)** Percentages of XCR1^+^ (left), CLEC9A^+^ (middle) and CD226^+^ (right) cells after 5 days of *in vitro* or *in vivo* reprogramming in CD44^+^eGFP^+^ cells for T98G and CD44^+^MCSP^+^eGFP^+^ for melanoma A375 and A2058. **(E)** Histograms (left) and quantification (right) of surface HLA-ABC molecules per cell gated in CD44^+^eGFP^+^ cells for T98G and CD44^+^MCSP^+^eGFP^+^ for melanoma A375 and A2058 (n=3). **(F)** Histograms (left) and quantification (right) of the percentages of cells expressing CD40 (n=3). Data in panels C-F are shown as mean ± SD.

### Reprogramming of human immunosuppressive tumor spheroids

To further support the feasibility of cDC1 reprogramming in tissues that include the presence of human immunosuppressive cells and soluble mediators of the TME, we first generated spheroids from human cancer cell lines and confirmed morphology and growth (Fig. 5A and fig. S8A-C). Using high-content fluorescence imaging and transduction of tumor cells with increasing multiplicities of infection (MOI) we detected increased transduction (mCherry^+^) and reprogramming in spheroids (CD45^+^ and HLA-DR^+^), and an overall decrease in the size of spheroids (Fig. 5B). This corresponds to the previously described loss of proliferation and tumorigenic potential accompanying cDC1 reprogramming (fig. S4H, I) (*23*). While reprogramming efficiency varied among human cancer types, these were similar or higher for each individual cell line reprogrammed in 3D when compared to 2D (Fig. 5C, D). T98G, IGR39, and A2058 cell lines reprogrammed more efficiently in spheroids (Fig. 5C, D), indicating that the 3D environment does not compromise but rather can favor reprogramming. To verify the establishment of a transcriptional cDC1 program and uncover potential differences between 2D and 3D, we performed scRNA-seq of reprogrammed cells at days 3, 7, and 9 (fig. S8D and data file S2). Transcriptomic analysis revealed faster upregulation of endogenous *SPI1*, *IRF8*, and *BATF3*, reprogramming markers (*PTPRC, HLA-DR*) and cDC1 genes (*C1orf54, CLEC9A, ZNF366*) in spheroid-derived cells (Fig. 5E). When compared to 2D cultures, reprogrammed cells in spheroids showed higher levels of transcriptional affiliation to cDC1s and rapid activation of the tumor-APC gene signature by day 3, which was established using commonly upregulated cDC1 genes during reprogramming of 18 human cancer cell lines (*23*) (Fig. 5F, G). In agreement, Reactome and Gene Ontology (GO) biological processes analysis of differentially expressed genes revealed the biggest differences at day 3, where terms associated with the immune system, GTPase signaling, and cell adhesion were enriched (fig. S8E). Interestingly, we detected increased expression of lymphotoxin beta (*LTB*) in spheroids at day 9 supporting TLS formation (*31*) (fig. S8F). Gene set enrichment analysis for immunogenic (TLR-induced maturation) or tolerogenic (homeostatic maturation) signatures (*32*) demonstrated that during reprogramming in 3D, PIB induced an immunogenic program despite the transient expression of homeostatic maturation genes at day 3 (fig. S8G). Indeed, reprogramming in 3D was accompanied by the activation of IFN, STING and NF-κB pathways (fig. S8H) reflecting a mature cDC1 program (*23*).

**Figure 5.**
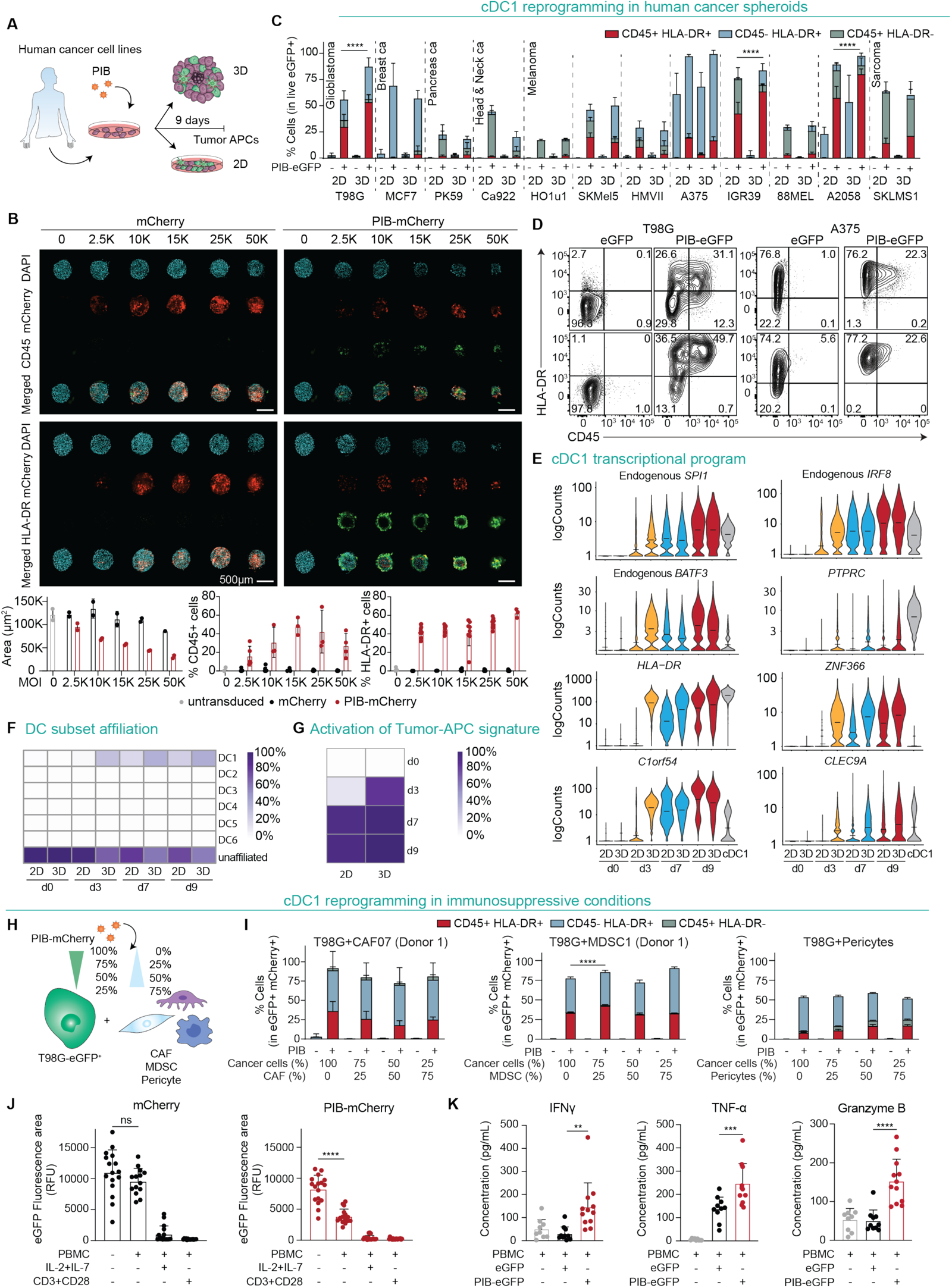
Reprogramming progresses in spheroids and in immunosuppressive tumor environments. **(A)** Experimental design to evaluate cDC1 reprogramming in human cancer spheroids. Cancer cells were transduced with PIB-eGFP or PIB-mCherry and used to form spheroids (3D) or cultured in monolayer (2D). **(B)** Confocal microscopy images and quantification of microtissue area and percentages of CD45^+^ and HLA-DR^+^ cells in T98G-derived spheroids after 9 days of reprogramming with PIB-mCherry at increasing multiplicities of infection (MOI) (n=2-10). **(C)** Flow cytometry quantification of cDC1 reprogramming efficiency in 12 human cancer cell lines in 2D and 3D at day 9 of reprogramming gated in transduced eGFP^+^ cells (n=4-12). Reprogramming efficiency was evaluated by flow cytometry as the percentage of CD45^+^HLA-DR^+^ cells (completely reprogrammed) and CD45^+^HLA-DR^-^ or CD45^-^HLA-DR^+^ cells (partially reprogrammed). **(D)** Representative flow cytometry plots showing phenotype of reprogrammed T98G and A375 cells in 2D and 3D compared to eGFP-transduced cells. **(E)** Reprogrammed and partially reprogrammed T98G cells were purified at reprogramming days 3, 7, and 9 and profiled by scRNA-seq. Violin plots show mRNA expression of endogenous transcription factors and cDC1 genes along the reprogramming time course in 2D and 3D. eGFP-transduced cells were used as day 0; donor peripheral blood cDC1 cells served as reference. **(F)** Integration of scRNA-seq data with data from published DC subsets (GSE94820) (*53*). Heatmap shows the percentage of cells transcriptionally affiliated with individual DC subsets. **(G)** Heatmap showing percentage of tumor-APC gene signature activation (*23*). **(H)** Experimental design to evaluate the effect of immunosuppression in cDC1 reprogramming using spheroids containing T98G-eGFP^+^ cells combined with cancer-associated fibroblasts (CAFs), myeloid-derived suppressor cells (MDSCs) or pericytes at indicated ratios. **(I)** Reprogramming efficiency gated in T98G-eGFP^+^ mCherry^+^ cells in spheroids with increasing proportions of CAFs (n=3-9, left), MDSC (n=3, middle), and pericytes (n=6-7, right). CAF07 and MDSC1 refer to cells from one individual donor. **(J)** Spheroid sizes as a measure of T cell cytoxicity against T98G-eGFP^+^ containing CAFs after 7 days of co-culture with non-activated HLA-A2-matched PBMCs or pre-activated with anti-CD3 and anti-CD28 antibodies or IL-2 and IL-7. Relative fluorescence units (RFU) were quantified by the eGFP^+^ fluorescence area by imaging (n=14-18). **(K)** Quantification of cytokine release 24 hours after co-culture of non-activated HLA-A2-matched PBMCs from three donors with T98G-eGFP^+^ spheroids containing CAFs (n=9-12). Data in panels B, C and I-K are shown as mean ± SD. Comparisons in panels C and I were analyzed using two-way ANOVA and in panels J and K using one-way ANOVA followed by Tukey’s multiple comparison test. ns - non-significant; **p<0.01; ***p<0.001; ****p<0.0001.

To evaluate the impact of immunosuppressive human TME components in cDC1 reprogramming, we initiated reprogramming in spheroids containing increasing proportions of cancer-associated fibroblasts (CAFs), myeloid-derived suppressor cells (MDSC), or pericytes (Fig. 5H and fig. S9A). Surprisingly, the presence of CAFs, MDSCs, or pericytes did not impair cDC1 reprogramming efficiency (Fig. 5I and fig. S9B, C), while the addition of anti-inflammatory cytokines IL-6, TGF-β, VEGF and immuno-regulatory GM-CSF only marginally reduced reprogramming (fig. S9D). Finally, we addressed the antigen presentation function of reprogrammed cancer cells in spheroids containing CAFs by co-culture with HLA-A2-matched or unmatched PBMCs quantifying spheroid size (as a measure of cytotoxicity) and T cell reactivity markers (as a measure of T cell activation). While control spheroids were not targeted by non-activated PBMCs, spheroids containing reprogrammed cells showed a reduction in size (Fig. 5J and fig. S9E) and elicited secretion of the PBMC-derived cytotoxic cytokines IFNγ, TNF-α and Granzyme B (Fig. 5K and fig. S9F). Together, these results demonstrate that cDC1 reprogramming in spheroids accelerates the kinetics of reprogramming eliciting an immunogenic cDC1 cell state, which goes in line with the findings of *in vivo* reprogramming in mice (Fig. 4B, C). Moreover, our findings indicate that the progression of cDC1 reprogramming and function lead to efficient immune cell activation and cytotoxicity within the human TME and is not affected by immunosuppression.

### Delivery of cDC1 reprogramming factors to tumors with adenoviral vectors

We next aimed to identify a platform to deliver PIB factors to tumors and elicit *in situ* cDC1 reprogramming. We compared the lentiviral vector (LV-PIB-eGFP) with non-integrative and replication-deficient adenoviral (Ad-PIB-eGFP) and adeno-associated viral vectors (AAV-PIB-eGFP) in monolayers, spheroids, and tumors *in situ* (Fig. 6A). First, we observed that the three viral vectors transduce mouse and human cancer cell lines (fig. S10A, B) while reprogramming efficiency and MHC-I expression was higher with Ad and LV when compared to AAV vectors (fig. S10C-E). We then profiled reprogramming kinetics and detected rapid phenotypic changes at day 3 with LV and Ad vectors (fig. S10F). In contrast, AAV showed low reprogramming efficiency at early time points, which did not reach the efficiencies of LV or Ad vectors (fig. S10F). Ad-mediated reprogramming also induced the expression of CD40 and CD226, confirming the acquisition of an immunogenic cDC1-like phenotype (fig. S10G, H).

**Figure 6.**
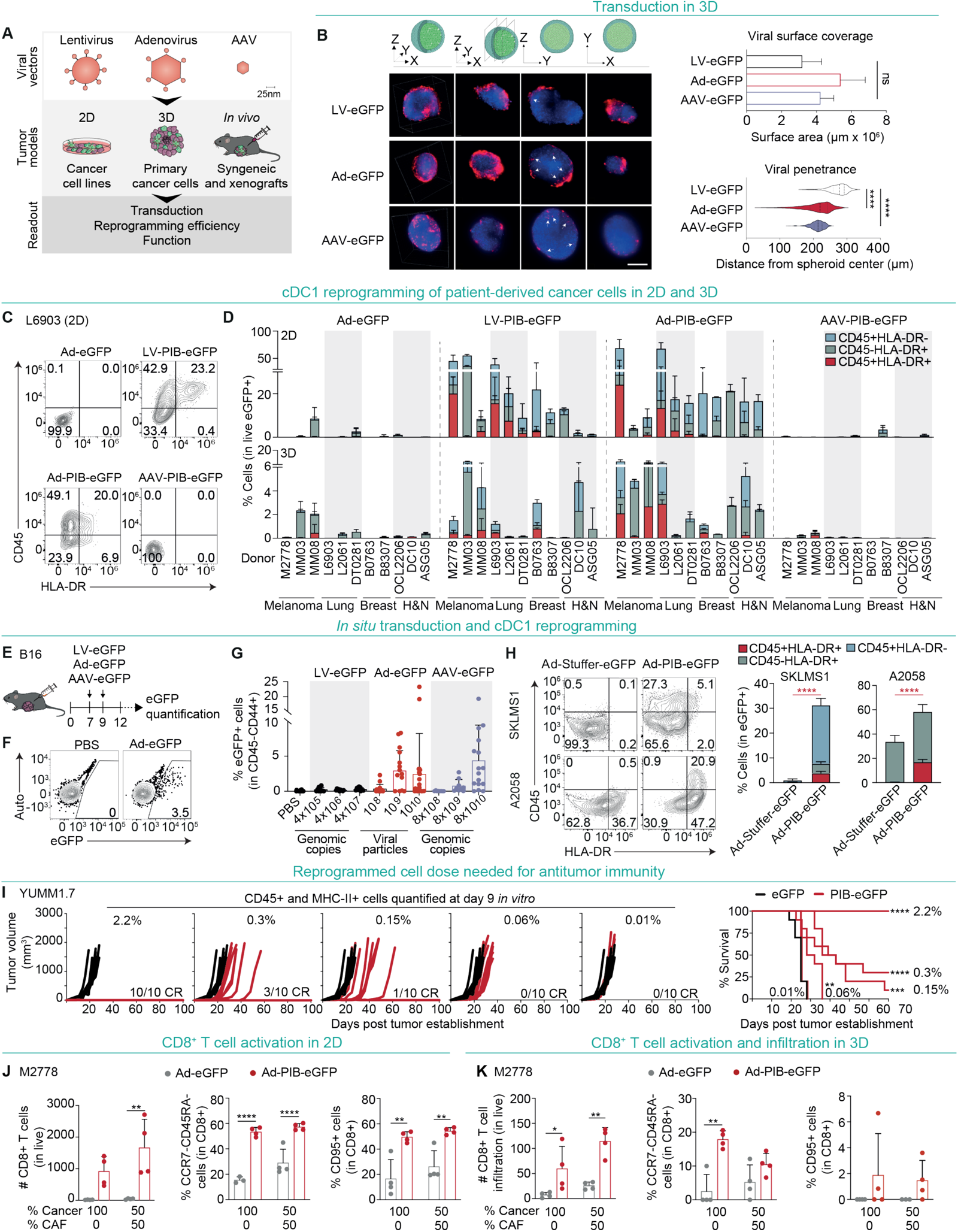
Efficient delivery of cDC1 reprogramming factors with adenoviral vectors. **(A)** Experimental design to prioritize a viral vector for delivery of PIB factors to tumors. Lentiviral (LV), adenoviral (Ad), and adeno-associated viral (AAV) transduction and reprogramming efficiencies were quantified using mouse and human cancer cell lines and patient-derived cancer cells in monolayer (2D), spheroids (3D), and tumors *in vivo*. **(B)** Light sheet microscopy pictures (left) and quantification (right) of T98G tumor spheroid transduction and penetration with eGFP-encoding LV, Ad and AAV vectors. Fixed tumor spheroids were stained against eGFP (red) and nuclei with DAPI (blue). The illustration (upper) visualizes the 3D image construction of the spheroid below. 3D image construction was performed by imaging and stacking of planes using fluorescence microscopy. Illustrated axes indicate the direction of the imaged planes. Viral surface coverage and penetrance were quantified by Zeiss Arivis image analysis software. Scale bar is 200 µm. **(C)** Representative flow cytometry plots and **(D)** quantification of reprogramming efficiency in patient-derived cancer cells 9 days post transduction in 2D or 3D with PIB-encoding LV, Ad and AAV vectors (LV-PIB-eGFP, Ad-PIB-eGFP, AAV-PIB-eGFP) (n=2-3). H&E, head and neck cancers. **(E)** Experimental design to evaluate transduction efficiency *in situ* using subcutaneous B16 tumors in C57BL/6J mice. Tumors were injected with 3 doses of LV-eGFP, Ad-eGFP, AAV-eGFP vectors or PBS at day 7 and 9 and isolated at day 12 for analysis (n=9-25). **(F)** Representative flow cytometry plots with transduction efficiency of Ad-eGFP when compared to PBS gated in CD45^-^CD44^+^ cells and **(G)** quantification of eGFP^+^ cells of tumors transduced with the 3 viral vectors or PBS. Quantification of viral particles is shown. **(H)** Human SKLMS1 and A2058 tumors were established in NXG mice and injected 4 times intratumorally with Ad-PIB-eGFP or Ad-Stuffer-eGFP at day 7, 9, 11 and 13 and analyzed at day 16 by flow cytometry. Representative flow cytometry plots (left) and quantification (right) of reprogramming efficiency gated in eGFP^+^ cells (n=8-10). Comparisons between CD45^+^HLA-DR^+^ populations (red) were used for statistical analysis. **(I)** YUMM1.7 tumors were established with decreasing doses of PIB-eGFP-transduced cells mixed with parental cell line. Percentages of reprogrammed cells (CD45^+^ and MHC-II^+^) in cell mixtures were quantified by flow cytometry at day 9 post transduction from parallel *in vitro* cultures. Tumor growth (left) and survival (right) are shown (n=10). The number of complete responses (CR) over the total number of mice per group is indicated. **(J)** Quantification of CD8^+^ T cell numbers (left), and percentages of effector CCR7^-^CD45RA^-^CD8^+^ (middle) and cytotoxic CD95^+^CD8^+^ T cells (right) after 8 days of co-culture with Ad-eGFP or Ad-PIB-eGFP transduced M2778 cells with (50%) or without (100%) CAFs in 2D. **(K)** Quantification of CD8^+^ T cell numbers within spheroids (left), and percentages of effector CCR7^-^CD45RA^-^CD8^+^ (middle) and cytotoxic CD95^+^CD8^+^ T cells (right). Data in panels B, D, G, H, J, and K are shown as mean ± SD. Comparisons in panels B were analyzed using One-Way ANOVA followed by Dunn’s multiple comparison test. Comparisons in panels H, J and K were analyzed using Mann Whitney test. Survival analysis in panel I was performed by log-rank Mantel-Cox test. ns - non-significant; **p<0.01; ***p<0.001; ****p<0.0001.

Next, we compared the capacity of the viral vectors to penetrate and transduce tissues (Fig. 6B). Ads and AAVs showed higher spheroid penetrance capacity (212.0±41.83µm and 212.6±18.54µm from spheroid center, respectively), in comparison to LVs (273.7±26.02µm). Ads and LVs showed the highest reprogramming efficiencies in 3D (fig. S10I). Importantly, in patient-derived cancer cells Ads were comparable to LVs in inducing cDC1 reprogramming and expression of HLA-ABC and CD40 in 2D and 3D (Fig. 6C, D and fig. S10J). To evaluate tumor transduction capacity *in situ*, we performed two consecutive intratumoral injections of LV-eGFP, Ad-eGFP, and AAV-eGFP vectors into B16 tumors and quantified the percentages of eGFP^+^ tumor cells (Fig. 6E). We observed high *in situ* transduction capacity with Ad-eGFP and AAV-eGFP when 10^9^ (2.94±2.9%) and 10^10^ (2.44±5.8%) viral particles or 8x10^10^ genomic copies (GC) were administered, respectively (Fig. 6F, G). In contrast, the transduction capacity of LV-eGFP was low even at the highest dose (0.3±0.2%) (Fig. 6G). We then injected intratumorally Ad-PIB-eGFP or a control Ad vector (Ad-Stuffer-eGFP) into two human xenograft models (A2058 melanoma and SKLMS1 sarcoma) and detected transduction (eGFP^+^ 5.0±2.7% SKLMS1; 0.3±0.2% A2058), *in situ* reprogramming (CD45^+^HLA-DR^+^ 3.9±0.7% SKLMS1; 16.6±2.5% A2058) and increased HLA-ABC expression (Fig. 6H and fig. S10K). In agreement with the required persistence *in vivo*, reprogrammed cells with adenoviral vectors could be maintained *in vitro* until day 20 independently of FLT3L signaling (fig. S10L). Next, we assessed the dose required to elicit antitumor immunity. An estimated 0.06% of reprogrammed cells corresponding to 0.3% eGFP^+^ cells were sufficient to reduce tumor growth and increase MS from 25 to 30 days (Fig. 6I and fig. S10M). Reassuringly, we observed 10% and 100% CRs with 0.15% and 2.2% of reprogrammed cells, respectively, which indicate that low cDC1 doses are sufficient for anti-tumor immunity.

To complement the validation of Ad-mediated PIB delivery we evaluated antigen presentation function in patient-derived cancer spheroids and profiled antitumor efficacy *in vivo*. Reprogrammed patient-derived melanoma cells induced activation and expansion of HLA-matched CD8^+^ T cells, giving rise to effector CCR7^-^CD45RA^-^ and cytotoxic CD95^+^CD8^+^ T cells (Fig. 6J), which also resulted in higher T cell infiltration into spheroids (Fig. 6J and 6K). *In vivo,* a ratio of 1:1 Ad-PIB transduced and parental B16 cells reduced tumor growth and extended MS (43 vs 19 days, p<0.001) (fig. S11A), increased p15E-specific CD8^+^ T cells in peripheral blood (fig. S11B) and induced control of contralateral tumor growth (fig. S11C). Overall, these data demonstrate that Ad vectors combine fast and efficient reprogramming with tumor transduction capacity *in situ*, providing a delivery platform for a cancer gene therapy approach based on cDC1 reprogramming.

### Systemic and durable antitumor immunity driven by gene therapy approach

To evaluate efficacy of a gene therapy based on *in situ* cDC1 reprogramming, we established B16 tumors and administered 4 intratumoral injections of either Ad-PIB, Ad-Stuffer or PBS at days 7, 9, 11 and 13 combined with anti-PD-1 and anti-CTLA-4 (Fig. 7A). Strikingly, we observed 50% CRs in mice treated with Ad-PIB, which remained tumor-free for 100 days (Fig. 7B). Two of these mice also developed vitiligo (fig. S11D). Within tumors, we detected an enrichment of CD45^+^ cells, primarily CD8^+^ T cells (Fig. 7C), which negatively correlated with the tumor volume (Fig. 7D and fig. S11E). Ad-PIB-treated tumors were enriched for effector T-bet^+^PD-1^-^CD8^+^ T cells and showed reduction in terminally exhausted T-bet^-^PD-1^+^CD8^+^ T cells (Fig. 7E). The ratio of pro-inflammatory CD4^+^ T helper cells to regulatory T cells was also elevated in Ad-PIB-treated mice, confirming the shift towards a pro-inflammatory response in the gene therapy setting (Fig. 7F). The intratumoral frequencies of myeloid cells remained unaltered (fig. S11F), but we again detected increased PD-L1 expression (Fig. 7G). Moreover, in both tdLN and non-draining lymph nodes (ndLN), we detected higher frequencies of p15E-specific CD8^+^ T cells and effector memory CD44^+^CD62L^-^CD8^+^ T cells (Fig. 7H, I and fig. S11G).

**Figure 7.**
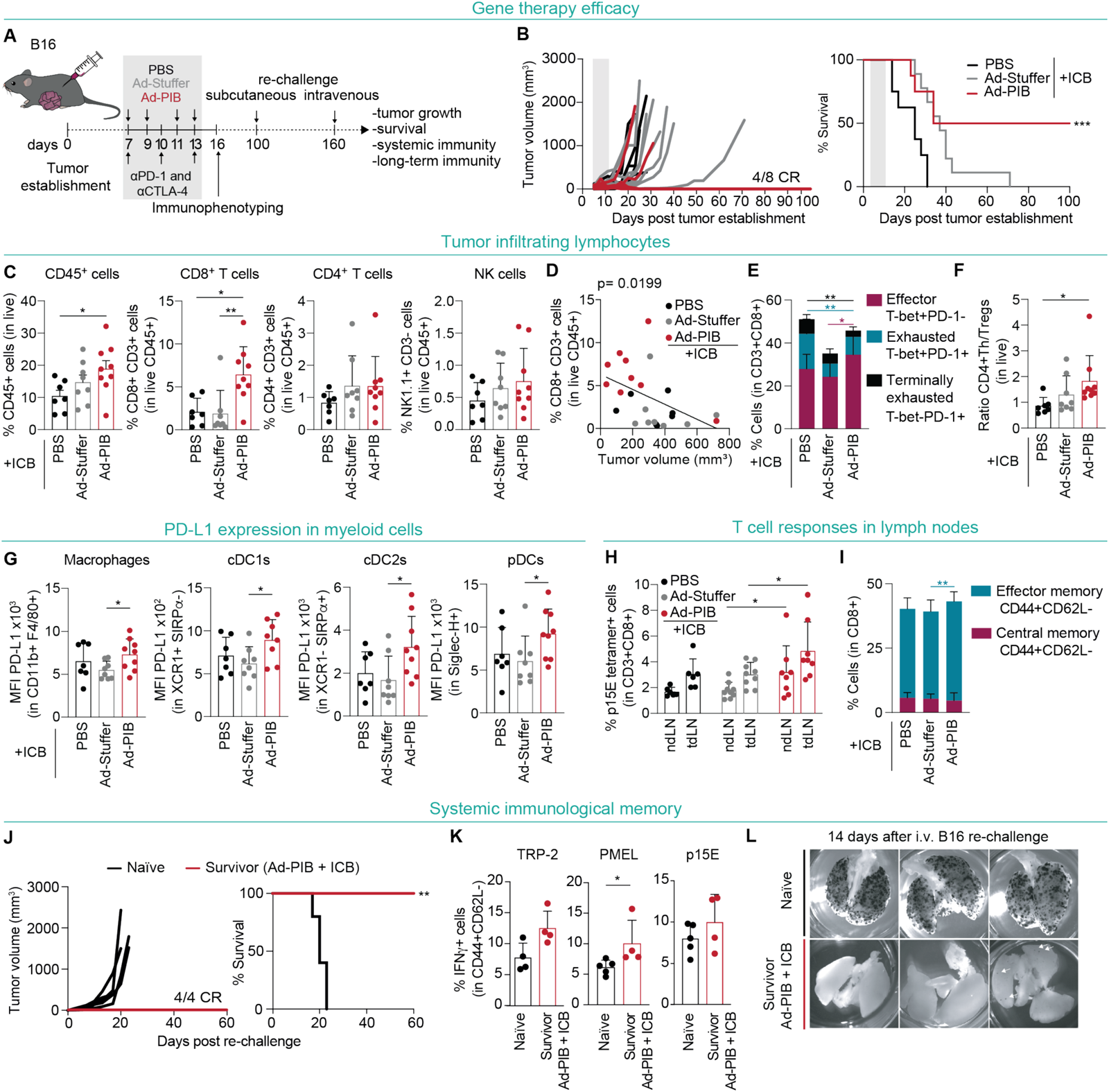
cDC1 reprogramming gene therapy elicits systemic and long-term antitumor immunity. **(A)** Experimental design to assess antitumor efficacy of Ad-PIB gene therapy in C57BL/6J mice with subcutaneous B16 melanoma tumors. Tumors were injected 4 times with Ad-PIB (red), non-coding Ad vector control (Ad-Stuffer, grey), or PBS (black) at day 7, 9, 11, and 13 after tumor establishment. Anti-PD-1 and anti-CTLA-4 (ICB) were administered intraperitoneally at day 7, 10, and 13. Survivor mice were further subcutaneously re-challenged with B16 cells at day 100 and intravenously at day 160. Grey box indicates the time of treatment. **(B)** Tumor growth (left) and survival (right) (n=8-10). The number of complete responses (CR) over the total number of mice per group is indicated. **(C)** Flow cytometry quantification of tumor-infiltrating lymphoid cells at day 16 (n=7-10). **(D)** Correlation of CD8^+^ T cell infiltration and tumor size. **(E)** Percentages of intratumoral T-bet^+^PD-1^-^ effector, T-bet^+^PD-1^+^ exhausted, and T-bet^-^PD-1^-^ terminally exhausted CD8^+^ T cells. Comparisons between the indicated color-coded populations were used for statistical analysis. **(F)** Ratio of intratumoral T-bet^+^CD44^+^CD4^+^ T helper (Th) cells and CD44^+^CD25^+^ T regulatory (Treg) cells. **(G)** PD-L1 expression in myeloid cells measured by mean fluorescence intensity (MFI). **(H)** Flow cytometry quantification of tumor antigen p15E-specific CD8^+^ T cells in tumor-draining lymph nodes (tdLN) and non-draining lymph nodes (n=7-10). **(I)** Percentages of CD44^+^CD62L^-^ effector memory and CD44^+^CD62L^+^ central memory CD8^+^ T cells in tdLN. **(J)** Survivor mice and naïve control mice were re-challenged subcutaneously with B16 cells. Tumor growth (left) and survival (right) are shown (n=4-5). **(K)** Flow cytometry quantification of tumor antigen-specific T cells from peripheral blood at day 14 after *in vitro* re-stimulation with peptides TRP-2-, PMEL- and p15E. Percentages of TRP-2-, PMEL- and p15E-specific IFNγ^+^CD44^+^CD62L^-^ effector memory CD8^+^ T cells (n=4-5). **(L)** Survivor mice were further re-challenged intravenously with B16 cells. Images of lungs from survivor and naïve mice 14 days after re-challenge (n=4). Data in panels C, E-I, and K are shown as mean ± SD. Comparisons in panel C, E, G, and I were analyzed using One-Way ANOVA followed by Dunn’s multiple comparison test. Comparisons in panels H and K were analyzed using the Mann Whitney test. Survival analyses in panel B and J was performed by log-rank Mantel-Cox test. *p<0.05, **p<0.01; ***p<0.001.

To test whether treatment with the Ad-PIB gene therapy induces immunological memory, we re-challenged survivor mice subcutaneously with parental B16 cells (Fig. 7A). While naïve mice developed tumors within 20 days, survivors remained tumor-free for another 60 days (Fig. 7J). In the peripheral blood of survivors, we also detected central memory CD62L^+^CD44^+^CD8^+^ T cells specific for the melanoma antigens PMEL and p15E (fig. S11H) and cytotoxic effector memory IFN-γ^+^CD62L^-^CD44^+^CD8^+^ T cells specific for TRP-2, PMEL, and p15E (Fig. 7K). Finally, we tested whether the gene therapy-induced systemic immune memory confers protection against metastatic tumor growth. Thus, we took advantage of the metastatic properties of B16 cells to colonize the lung and injected them intravenously at day 160 (Fig. 7A). Remarkably, survivor mice previously re-challenged subcutaneously also did not develop metastatic *foci* in the lungs (Fig. 7L and fig. S11I). Reassuringly, in mice that showed long-term tumor-free survival (>200 days) when compared to naïve mice, we could not detect differences in autoantibody levels in the plasma or off-target toxicity in internal organs (fig. S12). Ultimately, these results support a cDC1 reprogramming gene therapy modality for cancer immunotherapy which triggers systemic tumor-antigen specific T cell responses that lead to long-term and safe antitumor immunity.

## Discussion

In this study, we demonstrate that PU.1, IRF8 and BATF3-mediated cell fate reprogramming of tumor cells into immunogenic cDC1-like cells *in situ* represents a new immunotherapeutic modality for the treatment of solid tumors. Our results show that delivery of PIB factors to tumors *in situ* and tumor spheroids drives a cDC1 cell fate at phenotypic, transcriptomic, and functional levels, which results in TME remodeling, induction of tumor-specific T cell responses and leads to systemic and long-lasting antitumor immunity.

*In vivo* reprogramming results in cells with more mature phenotypes, improved function and increased fidelity measured by molecular affiliation to their natural counterparts (*17*), even when induced at anatomical locations that do not constitute their natural niche. For instance, β-cells reprogrammed in the liver and intestine also showed improved reprogramming fidelity and insulin secretion (*33*, *34*). In this study, we show that eliciting cDC1 reprogramming in spheroids and *in vivo* accelerated reprogramming and improved fidelity. This may be due to the availability of general tissue factors that are not supplied *in vitro* or specific cues available in the TME. Tissue-related cues such as hypoxia or mechanotransduction were shown to improve the efficiency of iPSC reprogramming (*35*, *36*). Hypoxia and mechanotransduction, through integrin and GTPase signaling, induce increased chromatin accessibility and transcription factor binding (*35*, *36*). Cell-to-cell contact and GTPase activity were enriched terms in cells reprogrammed in spheroids, which may explain enhanced reprogramming. Within the TME of solid tumors, hypoxia and immunosuppression are drivers of resistance to ICB, CAR-T cell therapy or dendritic cell vaccination (*37*). In contrast, TME-derived signals may be beneficial for cDC1 reprogramming. Tumors can also interrupt the maturation of cDC1s systemically by downregulation of IRF8 expression (*38*). The lack of sensitivity to immunosuppression may come from the enforced IRF8 expression during reprogramming compensating for extrinsic immunosuppressive signaling.

Previous studies have emphasized the role of cDC1s in antitumor immunity by activating CD8^+^ T cells and licensing NK cells (*9*). Although CD4^+^ T cell priming has been mainly attributed to cDC2s, recent studies suggest that cDC1s bear greater capacity to process cell-associated antigens for MHC class II presentation and prime CD4^+^ T cells (*39–41*). By replenishing cDC1-like cells within tumors we identified CD4^+^ T cells as critical effector cells driving tumor regression by polyclonal expansion. CD8^+^ T cell polyclonality was described as a requirement for positive treatment outcome in patients receiving PD-1 blockade therapy (*2*). Given the recently described cross-talk between cDC1, CD8^+^ and CD4^+^ T cells, which were associated with positive outcome (*41*), it is likely that these triad interactions may also occur between reprogrammed cancer cells and T cells locally in the TME eliciting a response without the requirement for migration to lymph nodes. While the importance of CD4^+^ T cells in YUMM1.7 tumors was clear, the magnitude may depend on the tumor model, as regressing B16 tumors showed higher CD8^+^ T cell infiltration. Nonetheless, in both melanoma models the ratio of pro-inflammatory CD4^+^ T helper cells to Tregs was increased, implying that *in vivo* cDC1 reprogramming drives a potent CD4^+^ T cell response. In addition to cell-mediated immunity, cDC1s can also orchestrate B cell responses and humoral immunity (*42*). Intratumoral B cells are also a reflection of TLS formation, which contain TCF-1^+^ stem-like, memory T cells and Tfh cells that support B cell maturation (*43*). TLS formation in melanoma tumors can be used as a predictive biomarker for survival and response to immunotherapy (*43*). We observed the induction of TLS-like structures by *in vivo* reprogramming of melanoma cells, leading to an increase of intratumoral TCF-1^+^ T cells, B cells, and circulating Tfh cells. *In vivo* cDC1 reprogramming, thus, provides a strategy to induce the formation of TLS *de novo* in tumors, which in addition to the therapeutic potential offers a system to dissect the mechanisms underlying TLS neogenesis (*31*).

To enable clinical translation, we tested a gene therapy concept by employing non-integrative viral vectors to deliver PIB to tumors and induce cDC1 reprogramming *in situ*. Currently, only adenoviral vectors are approved for non-lytic intratumoral gene therapies, due to their high *in situ* transduction efficiency, safety profile, large cargo space, and fast transgene expression onset (*44*, *45*). Indeed, transduction and reprogramming data in spheroids, syngeneic and xenografts models support the superiority of adenoviral delivery of reprogramming factors to tumor cells. Interestingly, it was sufficient to generate 2% of transduced cells (that correspond to 0.15% of CD45^+^ and MHC-II^+^ cells) to induce tumor growth delay and regression, providing a major advantage over other intratumoral immunotherapies such as oncolytic viruses, suicide gene approaches, and expression of co-stimulatory molecules or cytokines, which require high tumor cell transduction. Nonetheless, increased numbers of reprogrammed tumor cells correlated with tumor growth delay and survival rates, suggesting that the transduction rate *in situ* is a critical parameter for efficacy. A limitation of our study is that we have not discerned if efficacy is determined only by the numbers of partially or completely reprogrammed cells or by their reprogramming fidelity. In the future, we envision testing transduction enhancers in the adenoviral capsid (*46*) and in the formulation such as syn-3 to maximize delivery (*45*), as well as epigenetic adjuvants to increase reprogramming efficiency (*47*). These experiments will help clarifying the relative contribution of reprogrammed cell number versus cell quality *in vivo*. To further develop an even more scalable immunotherapy it will also be interesting to test non-viral delivery methods such as RNA moieties, which are currently employed for the delivery of cancer vaccines (*48*), or a specific cocktail of small molecules, shown to be sufficient for human iPSC reprogramming (*49*).

In this study, we showed that *in vivo* cDC1 reprogramming induced long-term and durable tumor control in models with varied profiles of immunogenicity, T cell infiltration, mutational burden, and responsiveness to ICB. Notwithstanding the efficacy as monotherapy, it is important to note the synergy with ICB for treatment of immune-deserted and low immunogenic tumors (e.g., B16). One of the reasons for this can be associated with the increased PD-L1 expression observed on myeloid cells in the tumor. Importantly, the combination with ICB did not reveal obvious autoimmune reactions in survivor mice, however this may require further future exploration in aged mice or in models prone to develop autoimmunity in response to ICB (*50*). In addition to combination with ICB, cDC1 reprogramming *in situ* could potentially synergize with other modalities, e.g., adoptive T cells, by enhancing target antigen presentation and supporting infiltration of engineered T cells into solid tumors, or agonistic CD40 antibodies given the critical role of CD4^+^ T cells that might depend on CD40-CD40L interaction and the previously reported role in TLS-like structure induction in glioma (*51*). As the sequence of immunotherapy treatments has been shown to impact treatment efficacy (*52*), it will be interesting to evaluate this approach in the clinical setting as a neoadjuvant therapy. This could establish immunological competence before tumor resection and support the elimination of residual tumor cells and metastatic lesions.

Overall, we provide proof-of-principle that cDC1 reprogramming *in situ* represents a tumor origin-agnostic, off-the-shelf yet personalized immunotherapy modality able to orchestrate systemic and durable antitumor immunity. cDC1 reprogramming *in situ* offers the advantages of a precision cell therapy while overcoming the challenges of *ex vivo* cell manipulation. This study paves the way for first-in-human trials and lays the foundation for a new class of immunotherapies based on the unique function of immune cell subsets generated *in vivo* through cellular reprogramming.

## Material and Methods

### Study design

Here, we aimed to investigate whether *in situ* direct cell fate reprogramming of tumor cells into immunogenic cDC1-like cells within the TME elicits antitumor immunity to provide proof-of-principle for a new cancer immunotherapy modality. To this end, we divided this study into three parts: First, we evaluated the immunogenicity of reprogrammed cells generated *in vivo* using syngeneic mouse models of melanoma. To evaluate the efficiency of *in vivo* reprogramming independently of the delivery system, we transplanted mixtures of PIB-transduced cells with parental tumor cells. We characterized systemic antitumor immune responses as monotherapy or in combination with ICB by monitoring tumor growth and survival, inducing contralateral tumors and re-challenging survivors with tumor cells. We characterized changes in tumor-infiltrating and peripheral immune cells by flow cytometry and scRNA-seq with TCR enrichment. Secondly, we characterized reprogramming in human cancer spheroids and xenograft models. We evaluated reprogramming efficiencies in the presence of immunosuppressive TME components by flow cytometry and scRNA-seq. Thirdly, we compared lentiviral, adenoviral, and adeno-associated viral platforms to deliver PIB to tumors and evaluate antitumor efficacy using an adenoviral-based gene therapy approach.

### Mice

Animal care and experimental procedures were performed in accordance with the Swedish federal regulations after approval from the Swedish Board of Agriculture. B6.129S(C)-Batf3tm1Kmm/J (BATF3^KO^, The Jackson Laboratory) and C57BL/6-Tg(TcraTcrb)1100Mjb/J (OT-I, The Jackson Laboratory) mice were bred in-house. C57BL/6J, NOD.Cg-Prkdc^SCID^IL2rg^tm1^Wjl/SzJ (NSG, The Jackson Laboratory), NOD-Prkdc^SCIT^IL2rg^tm1^/Rj (NXG, Janvier Labs) and B6.SJL-Ptprc^a^Pepc^b^/BoyCrl females aged 6-8 weeks were purchased from Charles River or Janvier-Labs. Animals were housed in a controlled temperature environment (23±2 °C) and a fixed 12-hour light/dark cycle, having free access to food and water. Mice were age-matched, gender-matched and within the same gender randomly assigned to treatment or control groups in all experiments. Numbers of mice for *in vivo* experiments were determined based on previous expertise, and power analysis was not performed. Mice were sacrificed by cervical dislocation when endpoints were reached. Investigators were not blinded during experimental procedures or the assessment of outcomes.

### Cell culture

Mouse B16-F10, LLC, MC38 and human A375, A2058, HO1u1, IGR39, MCF7, PK59, SKLMS1, SKMel5, Ca922, 88MEL, T98G cancer cell lines, cancer-associated fibroblasts (CAF) and human embryonic kidney (HEK) 293T were maintained in Dulbecco’s modified Eagle’s medium (DMEM) supplemented with 10% (v/v) fetal bovine serum (FBS), 2 mM GlutaMAX, 1mM sodium pyruvate and 100 U/ml penicillin and 100 mg/ml streptomycin (DMEM complete). B16-F10 expressing Ovalbumin (B16-OVA) were maintained in DMEM complete supplemented with 0.4 mg/ml geneticin (Gibco). Mouse Panc02, B2905, MB49, BRAF^V600E^COX1/2^KO^ cancer cell lines, mouse CD103^+^ bone marrow-derived dendritic cells (BM-DC), primary mouse and human T cells were cultured in RPMI 1640 medium supplemented with 10% (v/v) FBS, 2 mM GlutaMAX, 1 mM sodium pyruvate, 50 mM 2-mercaptoethanol and 100 U/ml penicillin and 100 mg/ml streptomycin (RPMI complete). YUMM1.7 melanoma cells were cultured in DMEM/F-12 with 10% (v/v) FBS, 2 mM GlutaMAX, 0.1 mM non-essential amino acids, 1 mM sodium pyruvate and 100 U/ml penicillin and 100 mg/ml streptomycin (DMEM/F-12 complete). MDSCs were differentiated from monocytes obtained from PBMCs of healthy donors and cultured in RPMI complete. Human pericytes were cultured in Pericyte medium (ScienCell). Fibroblasts were expanded on tissue-culture plates coated with 0.1% gelatin. All cells were dissociated from tissue-culture plates using TrypLE Express for 5-10 minutes at 37°C, split at 80% confluency and maintained in a humid environment at 37°C and 5% CO_2_. Reagents used for cell culture were purchased from Thermo Fisher Scientific, STEMCELL Technologies, and Nordic Biolabs. Detailed information on cells used in the study and culture conditions are provided in data file S3.

### Primary patient samples

Human tumor specimens were obtained according to the Helsinki Declaration and the European Network of Research Ethics Committees. Primary cancer cells derived from melanoma, lung cancer, breast cancer, head and neck cancer (tonsil and tongue), and CAF cultures were either purchased from Amsbio, BioIVT, VitroBiopharma or provided by the National Center of Cancer Immune Therapy CCIT-DK or the Skåne University Hospital. Human brain vascular pericytes were purchased from ScienCell. Primary tumor tissue was processed according to a standardized digestion protocol as previously described (*23*). In brief, after receiving tumor tissue in cold PBS, samples were cut into pieces, fat and muscle tissue were removed and tumor fragments were further mechanically and enzymatically digested following the gentleMACS Octo Dissociator protocol (Miltenyi) using the 37°C_h_TDK_3 program. For tonsil and tongue cancer samples, tumor tissue was cut into pieces and enzymatically digested with 80 µg/ml of collagenase D (Sigma-Aldrich) and 25 µg/ml of DNAse I (Sigma-Aldrich) for 20 minutes at 37°C. During incubation, the mixture was inverted every 5 minutes. Single cell suspensions were obtained by passing digestion mixtures through a 70 µm strainer and then seeded on 0.1% gelatin-coated tissue culture plates. Culture conditions for primary cancer cells are detailed in data file S3.

### Molecular cloning

Polycistronic lentiviral vector expressing the mouse or human transcription factors PU.1, IRF8 and BATF3 separated by 2A self-cleaving peptide sequences under the control of a constitutive SFFV promoter, followed by IRES2-eGFP was cloned previously (*22*, *23*). To generate mCherry expressing vectors, we used the empty backbone pRRL.PPT-SFFV-MCS-IRES2 (SFFV-MCS) (*22*, *23*) and inserted the coding sequence for mCherry by infusion cloning downstream the IRES sequence to generate pRRL.PPT-SFFV-MCS-IRES2-mCherry (SFFV-mCherry). Thereafter, we cloned the polycistronic cassette for human PIB into the multiple cloning site (MCS) and generated pRRL.PPT-SFFV-PU.1-P2A-IRF8-T2A-BATF3-IRES2-mCherry (PIB-mCherry). To generate a lentiviral polycistronic construct for myeloid reprogramming, the coding sequences of mouse PU.1 and C/EBPα (PC) separated by a T2A sequence were cloned first into the MCS of the pFUW-tetO-MCS vector (*22*) followed by subcloning of the polycistronic cassette into the MCS of the pRRL.PPT-SFFV-MCS-IRES2-eGFP vector (PC-eGFP). Adenoviral vectors (Ad) and adeno-associated viral (AAV) vectors were cloned and produced at VectorBuilder. Replication-deficient adenoviral vectors pAd5-SFFV-PU.1-P2A-IRF8-T2A-BATF3 (Ad-PIB) and pAd5-SFFV-PU.1-P2A-IRF8-T2A-BATF3-CMV-eGFP (Ad-PIB-eGFP) with an eGFP sequence under the control of constitutive cytomegalovirus (CMV) promoter were generated. pAd5-CMV-eGFP (Ad-eGFP), pAd5-SFFV-Stuffer (Ad-Stuffer) and pAd5-SFFV-Stuffer-CMV-eGFP (Ad-Stuffer-eGFP) were cloned and used as controls. The stuffer sequence was derived from the genome of *E. Coli* as a non-coding sequence and designed to have the same base pair length as polycistronic PIB. For replication-deficient AAV vectors, pAAV6-SFFV-PU.1-P2A-IRF8-T2A-BATF3 (AAV-PIB) was cloned. To generate eGFP expressing AAV vectors, the stop codon from BATF3 was removed and the eGFP sequence cloned downstream, separated by a F2A sequence (AAV-PIB-eGFP). pAAV6-CMV-eGFP (AAV-eGFP) was used as control. Sequences were verified by Sanger sequencing. Plasmids and primers used for cloning and sequencing are listed in data file S3.

### Tumor establishment

To establish tumors, cancer cells were harvested with TrypLE Express, live cells counted by Trypan blue staining using an automated hemocytometer and injected subcutaneously into the right flanks of recipient mice in 100 µl of ice-cold PBS. Before injection, mice were anesthetized by an intraperitoneal injection of ketamine (135 mg/kg) and xylazine (3 mg/kg). For tumor growth and survival experiments, 1x10^5^ B16-F10, YUMM1.7 or 1x10^6^ B2905 in C57BL/6J mice, 1x10^5^ YUMM1.7 in NSG mice, or 1x10^5^ BRAF^V600E^COX1/2^KO^ cells in BATF3^KO^ mice were used. In bilateral tumor settings, 2x10^5^ B16-OVA, B16-F10 or YUMM1.7 were injected subcutaneously into the right flank and 1x10^5^ B16-OVA, B16-F10 or YUMM1.7 into the left flank. LLC tumors were formed by subcutaneous injection of 1x10^6^ cells into the upper right neck area. For immunophenotyping and immunofluorescence analysis of tumors established with mixtures of *in vitro* transduced and parental cells, a total of 1x10^6^ B16-F10 or YUMM1.7 cells were injected in C57BL/6J mice. For establishing xenograft models, we injected 5x10^6^ of human A375, A2058, T98G in NSG or A2058 and SKLMS1 cell lines into NXG mice. C57BL/6J, NSG and NXG mice were 6-12-week-old age-matched females and BATF3^KO^ mice were males and females 6-12 weeks old. Tumor volumes were monitored with a digital caliper and calculated using the formula V = L*W*H/2. Survival was determined by predefined endpoints such as tumor size reaching 1500 mm^3^, tumor ulceration, or signs of animal suffering. Animals were randomized for tumor establishment and again before treatment.

### Immune checkpoint blockade treatment

For single or combinatorial treatment with ICB, mice received 200 μg of anti-PD-1 (clone RMP1-14, BioXCell) and/or 200 μg of anti-CTLA-4 (clone 9H10, BioXCell) or rat 200 µg IgG2a (clone 2A3, BioXCell) and IgG2b (clone LTF-2, BioXCell) isotype control antibodies diluted in 100 µl PBS intraperitoneally at days 7, 10, and 13 after tumor establishment.

### *In vivo* reprogramming of tumor cells

To evaluate the immunogenicity of *in vivo* reprogrammed cells in syngeneic mouse melanoma models, we transduced cancer cells *in vitro* with lentiviral (PIB, PC) or adenoviral particles (Ad-PIB), and 16 hours post-transduction mixed with untransduced parental cancer cells in defined ratios and injected subcutaneously into the right flank of mice. Unless stated otherwise, cells were mixed at a 1:1 ratio of transduced and untransduced parental cancer cells. As controls, we used empty viral vectors (lentivirus control: eGFP, adenovirus control: Ad-Stuffer). Transduction with lentiviral vectors was performed in the presence of polybrene (8 µg/ml, Sigma-Aldrich). The MOI used for transduction and induction of reprogramming by lentivirus ranged between 5.5x10^7^ and 5.0x10^8^ GC per cell. Cell mixtures were also kept *in vitro* to estimate the percentages of transduced cells by eGFP expression at day 3 and reprogramming efficiency by CD45 and MHC-II expression at day 9 by flow cytometry. To establish a dose-response between the amount of reprogrammed cells with induced antitumor immunity, transduced cells were serially diluted parental cancer cells (1:1, 1:2, 1:4, 1:10 and 1:100 ratio) before subcutaneous injection into mice. For adenoviral-mediated reprogramming *in vivo*, cells were transduced with non-eGFP encoding vectors (Ad-PIB, control: Ad-Stuffer) at an MOI of 2,500 infective units (IFU) per cell. To characterize the *in vivo* reprogramming efficiency of human cancer cells, we used the human cancer cell lines T98G, A375, A2058 in NSG mice. Cells were transduced with lentiviral vectors (PIB-eGFP, control: eGFP) mixed with untransduced cells and injected subcutaneously 16 hours post transduction and kept *in vitro* for phenotypic profiling by flow cytometry. At days 3, 5 and 9 post tumor establishment, tumors were isolated and dissociated into single cell suspensions for flow cytometry analysis for tumor or melanoma markers (CD44, MCSP), reprogramming markers (CD45, HLA-DR), antigen presentation (HLA-ABC), co-stimulatory molecule (CD40) and cDC1 markers (XCR1, CLEC9A, CD226).

### Delivery of viral vectors *in situ*

To deliver viral vectors to tumors *in situ*, LV, Ad, AAV vectors were diluted in ice-cold PBS to reach a final volume of 30 µl and intratumorally injected when the size of tumors reached 30–90 mm^3^. Tumors that did not reach the required sizes were excluded from the experiment. To quantify *in vivo* transduction efficiency in B16 tumors, eGFP-encoding vectors were administered at day 7 and 9 post tumor establishment. 4x10^5^, 4x10^6^ and 4x10^7^ GCs of LV-eGFP, 10^8^, 10^9^ and 10^10^ viral particles (VPs) of Ad-eGFP and 8x10^8^, 8x10^9^ and 8x10^10^ GCs of AAV-eGFP were administered per injection. At day 12, tumor tissue was isolated and dissociated into single cell suspensions for flow cytometry analysis of transduction efficiency through quantification of eGFP^+^ cells within live CD44^+^CD45^-^ cells. To assess *in situ* reprogramming efficiency in human xenograft models or efficacy in the B16 model combined with ICB treatment, 10^10^ VPs of Ad-PIB-eGFP or Ad-Stuffer-eGFP were injected intratumorally at day 7, 9, 11 and 13 post tumor establishment. In human xenograft models vectors encoded also eGFP (Ad-PIB-eGFP, Ad-Stuffer-eGFP). At day 16 after human xenograft establishment in NXG mice, tumors were isolated, dissociated, and reprogramming efficiency quantified by flow cytometry.

### Single cell RNA sequencing with TCR enrichment of T cells

5’ scRNA-seq with TCR enrichment was performed on FACS-sorted CD45^+^CD3^+^ T cells isolated from tumors, tdLN and peripheral blood of animals 21 days after tumor establishment with subcutaneous injection of PIB-eGFP or eGFP-transduced YUMM1.7 cells mixed at a 1:1 ratio with untransduced parental cells. Tumors were processed into single cell suspensions and tdLN were mechanically dissociated with a plunger against a 50 µm cell strainer and collected in FACS buffer for staining. Blood samples were collected into K2-EDTA coated microvette tubes (Sarstedt) and further processed to remove erythrocytes through red blood cell lysis using BD Pharm Lyse lysing buffer (BD Bioscience). Single cell suspensions were pooled from 5 animals per treatment group and stained with anti-CD45 and anti-CD3 antibodies. 3,000-10,000 T cells were FACS-sorted, resuspended in PBS containing 0.04% bovine serum albumin (BSA, STEMCELL Technologies) and loaded on a 10x Chromium (10x Genomics) without multiplexing.

### Spheroid reprogramming

To test reprogramming of human cancer cell line-derived spheroids, cells were transduced with lentiviral vectors encoding for PIB-eGFP or PIB-mCherry 4 hours before spheroid formation using the forced-floating method. To assess reprogramming efficiency, spheroids were dissociated by incubation with TrypLE for 20 minutes followed by vigorous pipetting and analyzed for CD45 and HLA-DR expression by flow cytometry. As controls, cells were transduced with lentiviral vectors encoding eGFP or mCherry. Alternatively, reprogrammed cells were detected in non-dissociated spheroids using fluorescent microscopy. To analyze cDC1 reprogramming in spheroids at the transcriptional level, eGFP^+^ cells expressing CD45 and/or HLA-DR were FACS-purified at day 3, 7 and 9 of reprogramming and loaded on a 10x Chromium (10x Genomics) for scRNA-seq. Reprogramming under immunosuppressive cytokine conditions was performed using DMEM complete medium supplemented with human IL-6 (10-40 ng/ml; Peprotech), TGF-ϕ3 (50-200 ng/ml; Miltenyi), VEGF (25-100 ng/ml; Miltenyi), or GM-CSF (50-200 ng/ml; Miltenyi) during the 9 days of reprogramming. Medium with cytokines was replaced every two days. For reprogramming in heterotypic spheroids, T98G-eGFP^+^ cells were transduced with PIB-mCherry and mixed with CAFs, MDSCs or pericytes at decreasing percentages of cancer cells (100%, 75%, 50% and 25%). For immunohistochemistry, confocal imaging, ATP release assay, and co-cultures with PBMCs, spheroids were aggregated by centrifugation of 300 transduced cancer cells and 1,000 CAFs and maintained over 3 days without the addition of Matrigel (Corning). For transduction and reprogramming of primary cancer cell-derived spheroids with LV, Ad, AAV vectors, spheroids were generated with 2.5% Matrigel and after 3 days incubated with LV (65,000 GC/ml), Ad (5,000 IFU/ml) or AAV (250,000 GC/ml) vectors encoding for PIB-eGFP or eGFP. For transduction with lentiviral vectors, complete medium was supplemented with 8 μg/ml of polybrene. Spheroids were dissociated at day 9 for flow cytometry profiling.

### Statistical analysis

All statistical analyses were performed using GraphPad Prism or R software. Data was subjected to a normality test before using ANOVA, two-way ANOVA, Kruskal-Wallis or Mann-Whitney test and t-test. Statistical significance of two groups was determined using an unpaired two-tailed Mann-Whitney test or t-test. Group comparisons were performed using ANOVA and corrected by Dunn’s or Tukey’s multiple comparison test. To estimate statistically significant differences in the survival in multiple groups we used the log-rank Mantel-Cox test. Unless stated otherwise in the figure legends, data are shown as mean ± SD and n represents the total number of animals in *in vivo* experiments or biological replicates in *in vitro* experiments and experiments. Randomization was performed using the Microsoft Office Excel function (=RANDBETWEEN). Sample sizes were based on previous experience. Significance was considered with *p < 0.05; **p < 0.01; ***p < 0.001; ****p < 0.0001. Statistical tests and parameters for each experiment are reported in the respective figure legend.

## Acknowledgments

We thank the members of the Cell Reprogramming in Hematopoiesis and Immunity Laboratory for discussions and S. Pedreiro for technical assistance. We thank K. Zaret (University of Pennsylvania) for critical reviewing and suggestions on the manuscript. We thank the Center for Translational Genomics, SciLifeLab Lund for providing sequencing services, S. Strömblad from the Lund University Bioimaging Center and L. Olsson from the SpatialOmics Lund University core facility for assistance with immunofluorescence. We thank J. Idoyaga (Stanford University) and her group members for sharing the CCR7 staining protocol. We would also like to thank the Lund Stem Cell Center FACS facility for cell sorting assistance and the Centre for Comparative Medicine for animal facilities. We thank F. Granucci (University of Milano Bicocca, Italy), S. Hugues (University of Geneva, Switzerland), Elsa Kress (Antineo, France), G. Merlino (National Cancer Institute Maryland, USA), S. Kobold (Ludwig Maximilian University of Munich, Germany) and R. Marais (Cancer Research Manchester Institute, UK) for sharing mouse cell lines and F. Aguilo (Umeå University), K. Miharada (Kumamoto University, Japan) and Y. Nakamura (RIKEN, Japan) and G. B. Jönsson (Lund University) for providing human cancer cell lines. We also thank the healthy donors and the Center for Clinical Immunology and Transfusion Medicine at Skåne University Hospital for providing leukocyte concentrates (ethical permit: 2022:11).

## Author contributions

E.A., F.Å, M.S.N., A.R., O.Z., X.C., N.R., C.V., M.R., L.-A.L., T.S., X.H., and E.R., conducted reprogramming experiments *in vitro* and in spheroids and analyzed the data. E.A., F.Å., X.C. and T.B. conducted reprogramming experiments *in vivo* and analyzed the data. E.A., F.Å, A.R., N.R., I.A., C.V., and M.R. designed and performed functional experiments *in vitro*. E.A., F.Å, X.C., and T.B. performed immunophenotyping by flow cytometry. E.A., M.S.N., O.Z., A.R., N.R., C.V., M.R. and M.d.R.T performed immunohistochemistry analysis. I.K. M.S.N, O.Z. and E.A performed scRNA-seq analysis from spheroids and T cells. E.A. and O.Z. cloned lentiviral constructs. E.R. and F.F.R. designed Ad and AAV vectors. M.V.S., Ö.M., D.A., M.L., L.G., I.M.S., provided primary human cancer samples and discussed clinical applicability. C.-F.P., F.F.R., and C.F.P. provided input for research design, interpretation of results and edited the manuscript. C.-F.P., F.F.R., and C.F.P. attracted funding. E.A., F.F.R. and C.-F.P. conceptualized the study and wrote the manuscript.

## Data and materials availability

The sequencing data generated in this study are available from Gene Expression Omnibus (GEO) under accession codes GSE255385 (mouse scRNA-seq and TCR), GSE255536 (human spheroid scRNA-seq). Published datasets re-analyzed in this study are available under accession codes GSE94820, GSE71171, GSE224941, GSE184527 and GSE103618. Constructs and vectors used for reprogramming are available from Asgard Therapeutics under a material transfer agreement with the company. All other data needed to evaluate the conclusions in the paper are present in the paper or Supplementary Materials.

## References

1. P. Sharma, S. Hu-Lieskovan, J. A. Wargo, A. Ribas, Primary, Adaptive, and Acquired Resistance to Cancer Immunotherapy. Cell 168, 707–723 (2017).

2. C. Puig-Saus, B. Sennino, S. Peng, C. L. Wang, Z. Pan, B. Yuen, B. Purandare, D. An, B. B. Quach, D. Nguyen, H. Xia, S. Jilani, K. Shao, C. McHugh, J. Greer, P. Peabody, S. Nayak, J. Hoover, S. Said, K. Jacoby, O. Dalmas, S. P. Foy, A. Conroy, M. C. Yi, C. Shieh, W. Lu, K. Heeringa, Y. Ma, S. Chizari, M. J. Pilling, M. Ting, R. Tunuguntla, S. Sandoval, R. Moot, T. Hunter, S. Zhao, J. D. Saco, I. Perez-Garcilazo, E. Medina, A. Vega-Crespo, I. Baselga-Carretero, G. Abril-Rodriguez, G. Cherry, D. J. Wong, J. Hundal, B. Chmielowski, D. E. Speiser, M. T. Bethune, X. R. Bao, A. Gros, O. L. Griffith, M. Griffith, J. R. Heath, A. Franzusoff, S. J. Mandl, A. Ribas, Neoantigen-targeted CD8+ T cell responses with PD-1 blockade therapy. Nature 615, 697–704 (2023).

3. E. G. Bawden, T. Wagner, J. Schröder, M. Effern, D. Hinze, L. Newland, G. H. Attrill, A. R. Lee, S. Engel, D. Freestone, M. de Lima Moreira, E. Gressier, N. McBain, A. Bachem, A. Haque, R. Dong, A. L. Ferguson, J. J. Edwards, P. M. Ferguson, R. A. Scolyer, J. S. Wilmott, C. M. Jewell, A. G. Brooks, D. E. Gyorki, U. Palendira, S. Bedoui, J. Waithman, K. Hochheiser, M. Hölzel, T. Gebhardt, CD4+ T cell immunity against cutaneous melanoma encompasses multifaceted MHC II-dependent responses. Sci. Immunol. 9, eadi9517 (2024).

4. J. Larkin, V. Chiarion-Sileni, R. Gonzalez, J.-J. Grob, P. Rutkowski, C. D. Lao, C. L. Cowey, D. Schadendorf, J. Wagstaff, R. Dummer, P. F. Ferrucci, M. Smylie, D. Hogg, A. Hill, I. Márquez-Rodas, J. Haanen, M. Guidoboni, M. Maio, P. Schöffski, M. S. Carlino, C. Lebbé, G. McArthur, P. A. Ascierto, G. A. Daniels, G. V. Long, L. Bastholt, J. I. Rizzo, A. Balogh, A. Moshyk, F. S. Hodi, J. D. Wolchok, Five-Year Survival with Combined Nivolumab and Ipilimumab in Advanced Melanoma. N. Engl. J. Med. 381, 1535–1546 (2019).

5. S. Adams, S. Loi, D. Toppmeyer, D. W. Cescon, M. De Laurentiis, R. Nanda, E. P. Winer, H. Mukai, K. Tamura, A. Armstrong, M. C. Liu, H. Iwata, L. Ryvo, P. Wimberger, H. S. Rugo, A. R. Tan, L. Jia, Y. Ding, V. Karantza, P. Schmid, Pembrolizumab monotherapy for previously untreated, PD-L1-positive, metastatic triple-negative breast cancer: cohort B of the phase II KEYNOTE-086 study. Ann. Oncol., 405–411 (2019).

6. M. R. Patel, G. S. Falchook, K. Hamada, L. Makris, J. C. Bendell, A phase 2 trial of trifluridine/tipiracil plus nivolumab in patients with heavily pretreated microsatellite-stable metastatic colorectal cancer. Cancer Med. 10, 1183–1190 (2021).

7. D. A. Reardon, A. Omuro, A. A. Brandes, J. Rieger, A. Wick, J. Sepulveda, S. Phuphanich, P. de Souza, M. S. Ahluwalia, M. Lim, G. Vlahovic, J. Sampson, OS10.3 Randomized Phase 3 Study Evaluating the Efficacy and Safety of Nivolumab vs Bevacizumab in Patients With Recurrent Glioblastoma: CheckMate 143. Neuro. Oncol. 19, iii21–iii21 (2017).

8. S. Spranger, D. Dai, B. Horton, T. F. Gajewski, Tumor-Residing Batf3 Dendritic Cells Are Required for Effector T Cell Trafficking and Adoptive T Cell Therapy. Cancer Cell 31, 711–723.e4 (2017).

9. K. C. Barry, J. Hsu, M. L. Broz, F. J. Cueto, M. Binnewies, A. J. Combes, A. E. Nelson, K. Loo, R. Kumar, M. D. Rosenblum, M. D. Alvarado, D. M. Wolf, D. Bogunovic, N. Bhardwaj, A. I. Daud, P. K. Ha, W. R. Ryan, J. L. Pollack, B. Samad, S. Asthana, V. Chan, M. F. Krummel, A natural killer–dendritic cell axis defines checkpoint therapy– responsive tumor microenvironments. Nat. Med. 24, 1178–1191 (2018).

10. M. Hubert, E. Gobbini, C. Couillault, T.-P. V. Manh, A.-C. Doffin, J. Berthet, C. Rodriguez, V. Ollion, J. Kielbassa, C. Sajous, I. Treilleux, O. Tredan, B. Dubois, M. Dalod, N. Bendriss-Vermare, C. Caux, J. Valladeau-Guilemond, IFN-III is selectively produced by cDC1 and predicts good clinical outcome in breast cancer. Sci. Immunol. 5, eaav3942 (2020).

11. H. Salmon, J. Idoyaga, A. Rahman, M. Leboeuf, R. Remark, S. Jordan, M. Casanova-Acebes, M. Khudoynazarova, J. Agudo, N. Tung, S. Chakarov, C. Rivera, B. Hogstad, M. Bosenberg, D. Hashimoto, S. Gnjatic, N. Bhardwaj, A. K. Palucka, B. D. Brown, J. Brody, F. Ginhoux, M. Merad, Expansion and Activation of CD103+ Dendritic Cell Progenitors at the Tumor Site Enhances Tumor Responses to Therapeutic PD-L1 and BRAF Inhibition. Immunity 44, 924–938 (2016).

12. L. Heger, L. Hatscher, C. Liang, C. H. K. Lehmann, L. Amon, J. J. Lühr, T. Kaszubowski, R. Nzirorera, N. Schaft, J. Dörrie, P. Irrgang, M. Tenbusch, M. Kunz, E. Socher, S. E. Autenrieth, A. Purbojo, H. Sirbu, A. Hartmann, C. Alexiou, R. Cesnjevar, D. Dudziak, XCR1 expression distinguishes human conventional dendritic cell type 1 with full effector functions from their immediate precursors. Proc. Natl. Acad. Sci. 120, e2300343120 (2023).

13. P. Meiser, M. A. Knolle, A. Hirschberger, G. P. de Almeida, F. Bayerl, S. Lacher, A.-M. Pedde, S. Flommersfeld, J. Hönninger, L. Stark, F. Stögbauer, M. Anton, M. Wirth, D. Wohlleber, K. Steiger, V. R. Buchholz, B. Wollenberg, C. E. Zielinski, R. Braren, D. Rueckert, P. A. Knolle, G. Kaissis, J. P. Böttcher, A distinct stimulatory cDC1 subpopulation amplifies CD8+ T cell responses in tumors for protective anti-cancer immunity. Cancer Cell 41, 1498–1515.e10 (2023).

14. I. Heras-Murillo, I. Adán-Barrientos, M. Galán, S. K. Wculek, D. Sancho, Dendritic cells as orchestrators of anticancer immunity and immunotherapy. Nat. Rev. Clin. Oncol. 21, 257–277 (2024).

15. D. Srivastava, N. DeWitt, In Vivo Cellular Reprogramming: The Next Generation. Cell 166, 1386–1396 (2016).

16. Q. Zhou, J. Brown, A. Kanarek, J. Rajagopal, D. A. Melton, In vivo reprogramming of adult pancreatic exocrine cells to β-cells. Nature 455, 627–632 (2008).

17. L. Qian, Y. Huang, C. I. Spencer, A. Foley, V. Vedantham, L. Liu, S. J. Conway, J. Fu, D. Srivastava, In vivo reprogramming of murine cardiac fibroblasts into induced cardiomyocytes. Nature 485, 593–598 (2012).

18. O. Torper, D. R. Ottosson, M. Pereira, S. Lau, T. Cardoso, S. Grealish, M. Parmar, In Vivo Reprogramming of Striatal NG2 Glia into Functional Neurons that Integrate into Local Host Circuitry. Cell Rep. 12, 474–481 (2015).

19. K. Yao, S. Qiu, Y. V Wang, S. J. H. Park, E. J. Mohns, B. Mehta, X. Liu, B. Chang, D. Zenisek, M. C. Crair, J. B. Demb, B. Chen, Restoration of vision after de novo genesis of rod photoreceptors in mammalian retinas. Nature 560, 484–488 (2018).

20. Z. Guo, L. Zhang, Z. Wu, Y. Chen, F. Wang, G. Chen, In Vivo Direct Reprogramming of Reactive Glial Cells into Functional Neurons after Brain Injury and in an Alzheimer’s Disease Model. Cell Stem Cell 14, 188–202 (2014).

21. F. F. Rosa, C. F. Pires, I. Kurochkin, A. G. Ferreira, A. M. Gomes, L. G. Palma, K. Shaiv, L. Solanas, C. Azenha, D. Papatsenko, O. Schulz, C. R. E. Sousa, C. F. Pereira, Direct reprogramming of fibroblasts into antigen-presenting dendritic cells. Sci. Immunol. 3, eaau4292 (2018).

22. F. F. Rosa, C. F. Pires, I. Kurochkin, E. Halitzki, T. Zahan, N. Arh, O. Zimmermannová, G. Ferreira, H. Li, S. Karlsson, S. Scheding, C. F. Pereira, Single-cell transcriptional profiling informs efficient reprogramming of human somatic cells to cross-presenting dendritic cells. Sci. Immunol. 7, eabg5539 (2022).

23. O. Zimmermannova, A. G. Ferreira, E. Ascic, M. Velasco Santiago, I. Kurochkin, M. Hansen, Ö. Met, I. Caiado, I. E. Shapiro, J. Michaux, M. Humbert, D. Soto-Cabrera, H. Benonisson, R. Silvério-Alves, D. Gomez-Jimenez, C. Bernardo, M. Bauden, R. Andersson, M. Höglund, K. Miharada, Y. Nakamura, S. Hugues, L. Greiff, M. Lindstedt, F. F. Rosa, C. F. Pires, M. Bassani-Sternberg, I. M. Svane, C.-F. Pereira, Restoring tumor immunogenicity with dendritic cell reprogramming. Sci. Immunol. 8, eadd4817 (2023).

24. E. Pérez-Guijarro, H. H. Yang, R. E. Araya, R. El Meskini, H. T. Michael, S. K. Vodnala, K. L. Marie, C. Smith, S. Chin, K. C. Lam, A. Thorkelsson, A. J. Iacovelli, A. Kulaga, A. Fon, A. M. Michalowski, W. Hugo, R. S. Lo, N. P. Restifo, S. K. Sharan, T. Van Dyke, R. S. Goldszmid, Z. Weaver Ohler, M. P. Lee, C. P. Day, G. Merlino, Multimodel preclinical platform predicts clinical response of melanoma to immunotherapy. Nat. Med. 26, 781– 791 (2020).

25. M. H. Linde, A. C. Fan, T. Köhnke, A. C. Trotman-Grant, S. F. Gurev, P. Phan, F. Zhao, N. L. Haddock, K. A. Nuno, E. J. Gars, M. Stafford, P. L. Marshall, C. G. Dove, I. L. Linde, N. Landberg, L. P. Miller, R. G. Majzner, T. Y. Zhang, R. Majeti, Reprogramming Cancer into Antigen-Presenting Cells as a Novel Immunotherapy. Cancer Discov. 13, 1164–1185 (2023).

26. G. Ghislat, A. S. Cheema, E. Baudoin, C. Verthuy, P. J. Ballester, K. Crozat, N. Attaf, C. Dong, P. Milpied, B. Malissen, N. Auphan-Anezin, T. P. Vu Manh, M. Dalod, T. Lawrence, NF-kB–dependent IRF1 activation programs cDC1 dendritic cells to drive antitumor immunity. Sci. Immunol. 6, eabg3570 (2021).

27. M. Cohen, A. Giladi, O. Barboy, P. Hamon, B. Li, M. Zada, A. Gurevich-Shapiro, C. G. Beccaria, E. David, B. B. Maier, M. Buckup, I. Kamer, A. Deczkowska, J. Le Berichel, J. Bar, M. Iannacone, A. Tanay, M. Merad, I. Amit, The interaction of CD4+ helper T cells with dendritic cells shapes the tumor microenvironment and immune checkpoint blockade response. *Nat*. Cancer 3, 303–317 (2022).

28. G. Micevic, A. Daniels, K. Flem-Karlsen, K. Park, R. Talty, M. McGeary, H. Mirza, H. N. Blackburn, E. Sefik, J. F. Cheung, N. I. Hornick, L. Aizenbud, N. S. Joshi, H. Kluger, A. Iwasaki, M. W. Bosenberg, R. A. Flavell, IL-7R licenses a population of epigenetically poised memory CD8 + T cells with superior antitumor efficacy that are critical for melanoma memory. Proc. Natl. Acad. Sci. 120, e2304319120 (2023).

29. S. A. Oh, D. C. Wu, J. Cheung, A. Navarro, H. Xiong, R. Cubas, K. Totpal, H. Chiu, Y. Wu, L. Comps-Agrar, A. M. Leader, M. Merad, M. Roose-Germa, S. Warming, M. Yan, J. M. Kim, S. Rutz, I. Mellman, PD-L1 expression by dendritic cells is a key regulator of T-cell immunity in cancer. *Nat*. cancer 1, 681–691 (2020).

30. S. C. Wei, J. H. Levine, A. P. Cogdill, Y. Zhao, N. A. A. S. Anang, M. C. Andrews, P. Sharma, J. Wang, J. A. Wargo, D. Pe’er, J. P. Allison, Distinct Cellular Mechanisms Underlie Anti-CTLA-4 and Anti-PD-1 Checkpoint Blockade. Cell 170, 1120–1133.e17 (2017).

31. M. Ramachandran, A. Vaccaro, T. van de Walle, M. Georganaki, R. Lugano, K. Vemuri, D. Kourougkiaouri, K. Vazaios, M. Hedlund, G. Tsaridou, L. Uhrbom, I. Pietilä, M. Martikainen, L. van Hooren, T. Olsson Bontell, A. S. Jakola, D. Yu, B. Westermark, M. Essand, A. Dimberg, Tailoring vascular phenotype through AAV therapy promotes anti-tumor immunity in glioma. Cancer Cell 41, 1134–1151.e10 (2023).

32. L. Ardouin, H. Luche, R. Chelbi, S. Carpentier, A. Shawket, F. Montanana Sanchis, C. Santa Maria, P. Grenot, Y. Alexandre, C. Grégoire, A. Fries, T.-P. Vu Manh, S. Tamoutounour, K. Crozat, E. Tomasello, A. Jorquera, E. Fossum, B. Bogen, H. Azukizawa, M. Bajenoff, S. Henri, M. Dalod, B. Malissen, Broad and Largely Concordant Molecular Changes Characterize Tolerogenic and Immunogenic Dendritic Cell Maturation in Thymus and Periphery. Immunity 45, 305–318 (2016).

33. A. Banga, E. Akinci, L. V. Greder, J. R. Dutton, J. M. W. Slack, In vivo reprogramming of Sox9+ cells in the liver to insulin-secreting ducts. Proc. Natl. Acad. Sci. U. S. A. 109, 15336–15341 (2012).

34. C. Ariyachet, A. Tovaglieri, G. Xiang, J. Lu, M. S. Shah, C. A. Richmond, C. Verbeke, D. Melton, B. Z. Stanger, D. Mooney, R. A. Shivdasani, S. Mahony, Q. Xia, D. T. Breault, Q. Zhou, Reprogrammed Stomach Tissue as a Renewable Source of Functional β Cells for Blood Glucose Regulation. Cell Stem Cell 18, 410–421 (2016).

35. S. Park, J. Lee, K. S. Ahn, H. W. Shim, J. Yoon, J. Hyun, J. H. Lee, S. Jang, K. H. Yoo, Y. Jang, T. Kim, H. K. Kim, M. R. Lee, J. Jang, H. Shim, H. Kim, Cyclic Stretch Promotes Cellular Reprogramming Process through Cytoskeletal-Nuclear Mechano-Coupling and Epigenetic Modification. Adv. Sci. 10, 2303395 (2023).

36. Y. Yoshida, K. Takahashi, K. Okita, T. Ichisaka, S. Yamanaka, Hypoxia Enhances the Generation of Induced Pluripotent Stem Cells. Cell Stem Cell 5, 237–241 (2009).

37. C.-H. Tsai, Y.-M. Chuang, X. Li, Y.-R. Yu, S.-F. Tzeng, S. T. Teoh, K. E. Lindblad, M. Di Matteo, W.-C. Cheng, P.-C. Hsueh, K.-C. Kao, H. Imrichova, L. Duan, H. Gallart-Ayala, P.-W. Hsiao, M. Mazzone, J. Ivanesevic, X. Liu, K. E. de Visser, A. Lujambio, S. Y. Lunt, S. M. Kaech, P.-C. Ho, Immunoediting instructs tumor metabolic reprogramming to support immune evasion. Cell Metab. 35, 118–133.e7 (2023).

38. M. A. Meyer, J. M. Baer, B. L. Knolhoff, T. M. Nywening, R. Z. Panni, X. Su, K. N. Weilbaecher, W. G. Hawkins, C. Ma, R. C. Fields, D. C. Linehan, G. A. Challen, R. Faccio, R. L. Aft, D. G. Denardo, Breast and pancreatic cancer interrupt IRF8-dependent dendritic cell development to overcome immune surveillance. Nat. Commun. 9, 1250 (2018).

39. Y. Valdez, W. Mah, M. M. Winslow, L. Xu, P. Ling, S. E. Townsend, Major Histocompatibility Complex Class II Presentation of Cell-associated Antigen Is Mediated by CD8α+ Dendritic Cells In Vivo. J. Exp. Med. 195, 683–694 (2002).

40. D. J. Theisen, J. T. Davidson, C. G. Briseño, M. Gargaro, E. J. Lauron, Q. Wang, P. Desai, V. Durai, P. Bagadia, J. R. Brickner, W. L. Beatty, H. W. Virgin, W. E. Gillanders, N. Mosammaparast, M. S. Diamond, L. D. Sibley, W. Yokoyama, R. D. Schreiber, T. L. Murphy, K. M. Murphy, WDFY4 is required for cross-presentation in response to viral and tumor antigens. Science 362, 694–699 (2018).

41. S. T. Ferris, V. Durai, R. Wu, D. J. Theisen, J. P. Ward, M. D. Bern, J. T. Davidson, P. Bagadia, T. Liu, C. G. Briseño, L. Li, W. E. Gillanders, G. F. Wu, W. M. Yokoyama, T. L. Murphy, R. D. Schreiber, K. M. Murphy, cDC1 prime and are licensed by CD4+ T cells to induce anti-tumour immunity. Nature 584, 624–629 (2020).

42. Y. Kato, T. M. Steiner, H.-Y. Park, R. O. Hitchcock, A. Zaid, J. L. Hor, S. Devi, G. M. Davey, D. Vremec, K. M. Tullett, P. S. Tan, F. Ahmet, S. N. Mueller, S. Alonso, D. M. Tarlinton, H. L. Ploegh, T. Kaisho, L. Beattie, J. H. Manton, D. Fernandez-Ruiz, K. Shortman, M. H. Lahoud, W. R. Heath, I. Caminschi, Display of Native Antigen on cDC1 That Have Spatial Access to Both T and B Cells Underlies Efficient Humoral Vaccination. J. Immunol. 205, 1842–1856 (2020).

43. R. Cabrita, M. Lauss, A. Sanna, M. Donia, M. Skaarup Larsen, S. Mitra, I. Johansson, B. Phung, K. Harbst, J. Vallon-Christersson, A. van Schoiack, K. Lövgren, S. Warren, K. Jirström, H. Olsson, K. Pietras, C. Ingvar, K. Isaksson, D. Schadendorf, H. Schmidt, L. Bastholt, A. Carneiro, J. A. Wargo, I. M. Svane, G. Jönsson, Tertiary lymphoid structures improve immunotherapy and survival in melanoma. Nature 577, 561–565 (2020).

44. R. E. Sobol, K. B. Menander, S. Chada, D. Wiederhold, B. Sellman, M. Talbott, J. J. Nemunaitis, Analysis of Adenoviral p53 Gene Therapy Clinical Trials in Recurrent Head and Neck Squamous Cell Carcinoma. Front. Oncol. 11, 645745 (2021).

45. S. A. Boorjian, M. Alemozaffar, B. R. Konety, N. D. Shore, L. G. Gomella, A. M. Kamat, T. J. Bivalacqua, J. S. Montgomery, S. P. Lerner, J. E. Busby, M. Poch, P. L. Crispen, G. D. Steinberg, A. K. Schuckman, T. M. Downs, R. S. Svatek, J. Mashni, B. R. Lane, T. J. Guzzo, G. Bratslavsky, L. I. Karsh, M. E. Woods, G. Brown, D. Canter, A. Luchey, Y. Lotan, T. Krupski, B. A. Inman, M. B. Williams, M. S. Cookson, K. A. Keegan, G. L. Andriole, A. I. Sankin, A. Boyd, M. A. O’Donnell, D. Sawutz, R. Philipson, R. Coll, V. M. Narayan, F. P. Treasure, S. Yla-Herttuala, N. R. Parker, C. P. N. Dinney, Intravesical nadofaragene firadenovec gene therapy for BCG-unresponsive non-muscle-invasive bladder cancer: a single-arm, open-label, repeat-dose clinical trial. Lancet Oncol. 22, 107– 117 (2021).

46. A. A. Stepanenko, A. O. Sosnovtseva, M. P. Valikhov, A. A. Chernysheva, S. A. Cherepanov, G. M. Yusubalieva, Z. Ruzsics, A. V. Lipatova, V. P. Chekhonin, Superior infectivity of the fiber chimeric oncolytic adenoviruses Ad5/35 and Ad5/3 over Ad5-delta-24-RGD in primary glioma cultures. Mol. Ther. Oncolytics 24, 230–248 (2022).

47. D. Huangfu, K. Osafune, R. Maehr, W. Guo, A. Eijkelenboom, S. Chen, W. Muhlestein, D. A. Melton, Induction of pluripotent stem cells from primary human fibroblasts with only Oct4 and Sox2. Nat. Biotechnol. 26, 1269–1275 (2008).

48. L. A. Rojas, Z. Sethna, K. C. Soares, C. Olcese, N. Pang, E. Patterson, J. Lihm, N. Ceglia, P. Guasp, A. Chu, R. Yu, A. K. Chandra, T. Waters, J. Ruan, M. Amisaki, A. Zebboudj, Z. Odgerel, G. Payne, E. Derhovanessian, F. Müller, I. Rhee, M. Yadav, A. Dobrin, M. Sadelain, M. Łuksza, N. Cohen, L. Tang, O. Basturk, M. Gönen, S. Katz, R. K. Do, A. S. Epstein, P. Momtaz, W. Park, R. Sugarman, A. M. Varghese, E. Won, A. Desai, A. C. Wei, M. I. D’Angelica, T. P. Kingham, I. Mellman, T. Merghoub, J. D. Wolchok, U. Sahin, Ö. Türeci, B. D. Greenbaum, W. R. Jarnagin, J. Drebin, E. M. O’Reilly, V. P. Balachandran, Personalized RNA neoantigen vaccines stimulate T cells in pancreatic cancer. Nature 618, 144–150 (2023).

49. J. Guan, G. Wang, J. Wang, Z. Zhang, Y. Fu, L. Cheng, G. Meng, Y. Lyu, J. Zhu, Y. Li, Y. Wang, S. Liuyang, B. Liu, Z. Yang, H. He, X. Zhong, Q. Chen, X. Zhang, S. Sun, W. Lai, Y. Shi, L. Liu, L. Wang, C. Li, S. Lu, H. Deng, Chemical reprogramming of human somatic cells to pluripotent stem cells. Nature 605, 325–331 (2022).

50. K. Adam, A. Iuga, A. S. Tocheva, A. Mor, A novel mouse model for checkpoint inhibitor-induced adverse events. PLoS One 16, 1–14 (2021).

51. L. van Hooren, A. Vaccaro, M. Ramachandran, K. Vazaios, S. Libard, T. van de Walle, M. Georganaki, H. Huang, I. Pietilä, J. Lau, M. H. Ulvmar, M. C. I. Karlsson, M. Zetterling, S. M. Mangsbo, A. S. Jakola, T. Olsson Bontell, A. Smits, M. Essand, A. Dimberg, Agonistic CD40 therapy induces tertiary lymphoid structures but impairs responses to checkpoint blockade in glioma. Nat. Commun. 12, 4127 (2021).

52. J. Wei, W. Montalvo-Ortiz, L. Yu, A. Krasco, S. Ebstein, C. Cortez, I. Lowy, A. J. Murphy, M. A. Sleeman, D. Skokos, Sequence of αPD-1 relative to local tumor irradiation determines the induction of abscopal antitumor immune responses. Sci. Immunol. 6, eabg0117 (2021).

53. A.-C. Villani, R. Satija, G. Reynolds, S. Sarkizova, K. Shekhar, J. Fletcher, M. Griesbeck, Butler, S. Zheng, S. Lazo, L. Jardine, D. Dixon, E. Stephenson, E. Nilsson, I. Grundberg, D. McDonald, A. Filby, W. Li, P. L. De Jager, O. Rozenblatt-Rosen, A. A. Lane, M. Haniffa, A. Regev, N. Hacohen, Single-cell RNA-seq reveals new types of human blood dendritic cells, monocytes, and progenitors. Science 356, eaah4573 (2017).

54. A. G. Ferreira, O. Zimmermannova, I. Kurochkin, E. Ascic, F. Åkerström, C.-F. Pereira, Reprogramming Mouse and Human Cancer cells to Antigen Presenting Cells. Bio-Protocol. 13, e4881 (2023).

55. C. T. Mayer, P. Ghorbani, A. Nandan, M. Dudek, C. Arnold-Schrauf, C. Hesse, L. Berod, P. Stüve, F. Puttur, M. Merad, T. Sparwasser, Selective and efficient generation of functional Batf3-dependent CD103+ dendritic cells from mouse bone marrow. Blood 124, 3081–3091 (2014).

56. W. Wang, Z. Jiao, T. Duan, M. Liu, B. Zhu, Y. Zhang, Q. Xu, R. Wang, Y. Xiong, H. Xu, L. Lu, Functional characterization of myeloid-derived suppressor cell subpopulations during the development of experimental arthritis. Eur. J. Immunol. 45, 464–473 (2015).

57. J. Alquicira-Hernandez, A. Sathe, H. P. Ji, Q. Nguyen, J. E. Powell, scPred: accurate supervised method for cell-type classification from single-cell RNA-seq data. Genome Biol. 20, 264 (2019).

