## Supplemental Material for "A cancer immunotherapy modality based on dendritic cell reprogramming *in vivo*"

**This PDF file includes:**

Supplementary Methods
Figures S1 to S12
References (23, 32, *54*-*57*)

### Supplementary Methods

#### Viral production

Transfer plasmids encoding PU.1, IRF8 and BATF3 followed by IRES-eGFP (PIB-eGFP), eGFP, PIB-mCherry, mCherry, mOrange and bicistronic PU.1 and C/EBPα followed by IRES-eGFP (PC-eGFP) were used to produce lentiviral vectors. In experiments using lentiviral vectors for *in vitro* transduction, lentivirus was produced using the second-generation system as previously described (*23, 54*). In brief, human embryonic kidney (HEK) 293T cells were seeded in 15 cm plates to reach ~80% confluency and transfected with 7.5 µg packaging plasmid (psPAX2), 2.5 µg VSV-G-encoding envelope plasmid (pMD2), and 10 µg transfer plasmid combined with 60 µl of 1 mg/ml polyethyleneimine (PEI) in Opti-MEM. Virus-containing supernatants were collected after 36, 48, and 72 hours, filtered using 0.45 µm low protein binding cellulose acetate filters and concentrated 100-fold with Lenti-X Concentrator (Takara) before storage at -80°C. Alternatively, virus-containing supernatants were ultracentrifuged for 90 minutes at 4°C with 25,000 g in a SW 32 Ti Swinging-Bucket Rotor (Beckman Coulter). Lentiviral vector pellets were resuspended overnight in DMEM medium and stored in aliquots at -80°C. Lentiviral titers were quantified with the Lenti-X qRT-PCR titration kit (Takara) following the manufacturer's protocol.

In experiments using lentiviral vectors for *in situ* transduction, *in vivo* grade lentiviral particles were produced at VectorBuilder based on a third-generation system. In brief, HEK 293T cells were transfected with eGFP encoding transfer plasmid, envelope plasmid encoding VSV-G and two packaging plasmids encoding Gag/Pol and Rev. The supernatants were collected, and cell debris removed via centrifugation and filtration. Lentiviral particles were subsequently concentrated using polyethylene glycol (PEG) precipitation and further purified through sucrose cushion ultracentrifugation. Lentiviral titers were determined by quantifying the lentiviral p24 Gag protein using ELISA.

Adenoviral (Ad) and adeno-associated viral (AAV) vectors encoding for PIB (Ad-PIB, AAV-PIB) or PIB and eGFP (Ad-PIB-eGFP, AAV-PIB-eGFP), eGFP (Ad-eGFP, AAV-eGFP) or a non-coding stuffer sequence with or without eGFP (Ad-Stuffer, Ad-Stuffer-eGFP) were produced at VectorBuilder. Adenoviral vectors were packaged and amplified in HEK 293A cells. In brief, adenoviral plasmids containing PIB, PIB-eGFP, Stuffer or eGFP were first linearized by restriction digestion with Pacl. The linearized plasmid DNA was then transfected into HEK 293A expressing the adenovirus gene E1 to produce recombinant adenovirus. Adenoviral particles released into the culture medium were harvested and concentrated using cesium chloride (CsCl) gradient ultracentrifugation. The viral titer was determined by spectrophotometry (OD260) to quantify the number of viral particles and measured for the number of infective units (IFU) by immunocytochemistry staining for transduced cells via the detection of adenovirus-specific hexon protein.

For AAV production, HEK 293T cells were co-transfected with a helper plasmid encoding adenovirus genes (E4, E2A, and VA), an AAV helper plasmid encoding AAV rep and cap genes, and a transfer plasmid containing PIB-eGFP or eGFP. After incubation, AAV particles were harvested from cell lysates and supernatant and concentrated using polyethylene glycol (PEG) precipitation. Subsequently, AAV particles were further purified and concentrated through CsCl gradient ultracentrifugation. The viral titer was determined by quantitative PCR (qPCR) targeting the inverted terminal repeat sequence of AAV. Sodium dodecyl-sulfate polyacrylamide gel electrophoresis (SDS-PAGE) and silver staining were used to determine AAV purity. All viral vectors used for *in vivo* application successfully passed sterility and mycoplasma testing.

**Flow cytometry and FACS**

Surface marker analysis was performed on dissociated cells from *in vitro* 2D cultures or single cell suspensions of digested tissue from spheroids or tissues. Cells were stained with adequate antibodies diluted in phosphate-buffered saline (PBS) supplemented with 2% FBS (FACS buffer) at 4°C for 20-30 minutes in the presence of 1% mouse or rat serum, for human and mouse cells, respectively, to block unspecific binding. CCR7 staining was performed at 37°C for 30 minutes before staining for additional surface markers. Annexin V staining was performed using the Annexin V Apoptosis Detection Kit (Thermo Fisher Scientific) following manufacturer's recommendations. In brief, cells were first stained with antibodies against surface markers, washed with PBS and binding buffer, and then incubated with fluorophore-conjugated Annexin V at room temperature for 15 minutes. Tetramer staining was performed at room temperature for 30 minutes before surface marker staining and cell fixation using 4% paraformaldehyde (PFA, Thermo Fisher Scientific) for 20 minutes at 4°C without permeabilization. Intracellular staining for cytokines or proliferation marker Ki67 was performed using the Cytofix/Cytoperm Fixation/Permeabilization Kit (BD Biosciences) following the manufacturer's recommendation. Intranuclear transcription factor staining was performed using the True-Nuclear Transcription Factor Buffer Set (Biolegend) following the manufacturer's recommendation. For flow cytometry analysis requiring cell fixation, cells were stained with fixable viability dye (FVD) 450 or 520 (Thermo Fisher Scientific) before surface marker staining and fixation. For analysis without cell fixation, dead cells were stained by addition of 4′,6-diamidino-2-phenylindole (DAPI) or 7-aminoactinomycin D (7-AAD) to the cell suspension after surface marker staining and before acquisition. Flow cytometry analysis was performed on LSR Fortessa, LSR Fortessa X20, FACSymphony A1 and Beckman Coulter Life Sciences CytoFlex Benchtop flow cytometers. FACS-sorting was performed on a BD FACS Aria III or on a FACSymphony S6 sorter, using a 100 μm nozzle. FACS data were analyzed using FlowJo v.10.0.7 (FlowJo LLC). Gates were determined according to fluorescence minus one (FMO) controls. Antibodies used in this study are listed in data file S3.

***In vitro* transduction and reprogramming of tumor cells**

*In vitro* reprogramming mediated by lentiviral vectors was performed as previously described (*23, 54*). In brief, 0.5-1x$10^6$ cells were plated per tissue culture 6-well or 10-cm plate and incubated overnight with 5.5E+07 and 5.0E+08 genomic copies (GC) per cell in the presence of 8 μg/ml polybrene. Adenoviral and adeno-associated viral vectors were used at a multiplicity of infection of 5,000 IFU per cell for Ad vectors and 250,000 GC per cell for AAV vectors in tissue culture

12-well, 6-well or 10-cm plates. Transduction was performed in 5 ml in a 10-cm plate, 1 ml per well in a 6-well plate, or 0.5 ml per well in a 12-well plate. After 16h, virus-containing medium was replaced with fresh medium and cells were maintained until day 9 in culture with regular medium changes every 2-3 days and splitted 1:6 when 80% confluency was reached. Transduction efficiency was measured by flow cytometry using eGFP expression. cDC1 reprogramming efficiency was measured by flow cytometry analysis of CD45 and MHC-II/HLA-DR expression within live or live eGFP$^+$/mCherry$^+$ cells. Macrophage reprogramming efficiency was measured by CD45 and CD11b expression within live eGFP$^+$ cells. In addition, the expression of MHC-I/HLA-ABC, the co-stimulatory molecule CD40 and cDC1 markers XCR1, CLEC9A and CD226 was quantified by flow cytometry. Quantification of MHC-I/HLA-ABC surface molecules per cell was performed using the PE Phycoerythrin Fluorescence Quantitation Kit (BD Biosciences) following manufacturer's instructions. For monitoring the persistence of adenovirus-mediated eGFP expression and reprogramming, cells were maintained in complete media or with the addition of 100 ng/ml soluble FLT3 ligand (FLT3L, Miltenyi) until day 20 after transduction.

**MACS enrichment**

To purify *in vitro* reprogrammed mouse cancer cells by magnetic-activated cell sorting (MACS, Miltenyi) with high yields, we followed a protocol as previously described (*54*). Briefly, cells were resuspended in cold FACS buffer to reach a concentration of $10^7$ cells/ml and incubated with rat serum for 15 minutes, followed by 5 minutes incubation with biotinylated antibodies. To purify cDC1-like cancer cells, we used biotinylated CD45 and MHC-II antibodies. To purify macrophage-like cancer cells, we used biotinylated CD45 and CD11b antibodies. Cells were washed twice with FACS buffer before incubation with magnetic anti-biotin microbeads for 15 minutes. All incubations were performed on ice and labelled cells were purified using LS columns (Miltenyi) according to the manufacturer's recommendations. To purify naïve ovalbumin-specific CD8$^+$ T cells from spleens of OT-I mice, we isolated and homogenized spleens by plunging against a 40 µm cell strainer. Red blood cells were lysed with Pharm lyse lysing buffer (BD Biosciences) for 8 minutes protected from light at room temperature and naïve CD8$^+$ T cells were purified by MACS using the naïve mouse CD8$^+$ T cell isolation kit (Miltenyi Biotec).

To purify human HLA-A2$^+$ or HLA-A2$^-$ CD8$^+$ T cells, peripheral blood mononuclear cells (PBMCs) from HLA-A2$^+$ and HLA-A2$^-$ donors were centrifuged at 350g for 5 minutes at room temperature, resuspended in FACS buffer and CD8$^+$ T cells were isolated using the human CD8$^+$ T cell isolation kit (Miltenyi) according to manufacturer's recommendations.

**Tumor treatments with transfer of reprogrammed cDC1-like cells generated *in vitro***

To treat established tumors with *in vitro* generated cDC1-like cancer cells, we injected intratumorally 1-2x$10^5$ MACS-purified cells per tumor at day 7, 10 and 13 post tumor establishment as previously described (*54*). In brief, B16-derived or B2905-derived cDC1-like cells were generated by transduction of cancer cells with lentiviral vectors encoding for PIB-eGFP and maintained *in vitro* until day 5 of reprogramming. At day 5, CD45$^+$ and MHC-II$^+$

reprogrammed cells were purified by MACS (Miltenyi), counted, and resuspended in ice-cold PBS. Injection of eGFP-transduced cells and PBS were included as controls. For bilateral B16-OVA tumor growth experiments, B16-derived cells were pulsed 24h before injection with 1 mg/ml ovalbumin protein and 10 µg/ml toll-like receptor (TLR) 3 agonist polyinosine-polycytidylic acid (P(I:C)), washed extensively, and injected into right-flank B16-OVA tumors. For B2905 tumor growth experiments, B2905-derived cDC1-like cells were stimulated 24h before injection with 10 µg/ml P(I:C) or left unstimulated, washed extensively, and injected intratumorally at day 7, 10 and 13.

**Generation of bone marrow-derived CD103$^+$ dendritic cells**

Mouse CD103$^+$ bone marrow-derived dendritic cells (BM-DCs) were generated as previously described (*55*). In brief, bone marrow (BM) cells were harvested from long bones of the leg (tibias and femurs) by crushing. Cells were then harvested in PBS supplemented with 2% FBS, filtered with a 40 µm cell strainer and plated in Petri dishes at a density of 1.5x$10^6$ cells per mL of RPMI complete medium supplemented with 5 ng/ml GM-CSF (Miltenyi) and 200 ng/ml Fltl3L (Miltenyi) for 16 days (*55*).

**Cross-presentation assay**

Naïve ovalbumin-specific CD8$^+$ T cells from spleens of OT-I mice were enriched using a naïve mouse CD8$^+$ T cell isolation kit (Miltenyi Biotec). Enriched CD8$^+$ T cells were labeled with CellTrace Violet (CTV, Thermo Fisher Scientific) according to manufacturer's protocol. Parental B16 cells that do not overexpress OVA were transduced with lentiviral particles encoding PIB-eGFP or PC-eGFP and reprogrammed for 9 days into cDC1-like or macrophage-like cancer cells, respectively. MACS-purified reprogrammed cancer cells, eGFP-transduced cancer cells, and CD103$^+$ BM-DCs were incubated overnight at 37°C with ovalbumin protein (100 µg/ml) in the presence of P(I:C) (10 µg/ml) and extensively washed. Then, 1x$10^4$ cells were incubated with 1x$10^5$ naïve CTV-labeled OT-I CD8$^+$ T cells in 96-well U-bottom plate. After 3 days of co-culture, T cells were collected, stained, and analyzed by flow cytometry. T cell proliferation was determined by dilution of CTV staining and upregulation of CD44 expression. The threshold for data plotting was fixed at 100 events within live CD8$^+$ T cell gating.

**T cell re-stimulation of mouse peripheral blood**

Peripheral CD8$^+$ and CD4$^+$ T cells from treated and control animals were isolated from peripheral blood by tail vein puncture and collected into K2-EDTA coated microvette tubes (Sarstedt). Erythrocytes were lysed using Pharm lyse lysing buffer (BD Bioscience). After lysis, live cells were resuspended in RPMI complete medium and plated in 96-well U-bottom plates at 2x$10^5$ cells per well for antigen-agnostic or antigen-specific T cell re-stimulation.

For antigen-agnostic T cell re-stimulation, 1x$10^5$ cancer cells were plated per well and stimulated with 100 ng/ml mouse IFNγ (Miltenyi) 24 hours before start of the co-culture with peripheral blood T cells. Before addition of T cells, medium supplemented with IFNγ was removed, cancer cells were washed with PBS and T cells were added in RPMI complete medium. After 1 hour, brefeldin A (BD Biosciences) was added to block secretion of intracellular proteins and the co-culture continued for 5 hours. For antigen-specific T cell re-stimulation, $1x10^4$ $CD103^+$ BM-DCs were plated into 96-well U-bottom plates with 10 μg/ml of tumor antigen-derived peptides TRP-2 (MBL), PMEL (AnaSpec) and p15E (MBL) 24 hours before addition of T cells. BM-DC and T cell co-cultures were performed for 16 hours, then brefeldin A was added to the medium and cells were incubated for additional 5 hours. Intracellular production of IFNγ in T cells was analyzed by flow cytometry following cell fixation and intracellular antibody staining.

**Tumor re-challenge**

After 100 days of tumor-free survival with *in vivo* reprogramming or Ad-PIB gene therapy treatment, C57BL/6J survivor mice were re-challenged with $1x10^5$ B16-F10 or YUMM1.7 cells and $BATF3^{KO}$ survivor mice with $1x10^5$ $BRAF^{V600E}COX1/2^{KO}$ cells by subcutaneous injection into the same flank of the regressed tumor.

For metastatic challenge after 160 days of tumor-free survival with Ad-PIB gene therapy treatment, C57BL/6J survivor mice were injected with $1x10^5$ B16-F10 cells in 100 μl PBS into the lateral tail vein. 21 days after intravenous challenge, animals were sacrificed by cervical dislocation and lungs were carefully removed. Lungs were inflated and placed in 10% formalin (Sigma-Aldrich). Lung tissues were imaged by Leica Light Microscope and metastatic *foci* quantified by Fiji software (ImageJ2, Ver. 2.14.0/1.54f). An intensity threshold was set to identify and count dark metastatic foci. The area of *foci* was calculated against the total lung area to obtain coverage of *foci*.

**Immunohistochemistry and immunofluorescence**

For mouse tumor tissue images, tumors were excised at day 9 post tumor establishment with a 1:2 mixture of eGFP- or PIB-eGFP-transduced and untransduced parental cells. Tissues were fixed in 10% formalin for 24 hours, dehydrated for 72 hours by incubation with increasing ethanol concentrations (70%, 95% and absolute ethanol), and embedded in paraffin after xylene treatment. 5 μm thick tissue sections were cut and mounted onto BOND Plus slides (Leica). The paraffin-embedded sections were de-paraffinized and stained with Hematoxylin and Eosin (H&E) or used for immunofluorescence imaging. For immunofluorescence analysis, antigen retrieval was performed for the tissue sections using a steamer at 99°C in TRIS/EDTA buffer (Biotechne) for CD45 and eGFP staining or using low pH 10 mM Sodium Citrate buffer (Thermo Fischer Scientific) for CD19, CD4, CD8 and podoplanin (PDPN) staining. After antigen-retrieval, sections were blocked for 1 hour and antibody staining was performed in a humidified chamber. Stainings for eGFP and CD45 were performed for 2 hours at room temperature with fluorophore-conjugated antibodies (Novus Biologicals) anti-eGFP-AF594 (5 μg/ml) and anti-CD45-AF647 (20 μg/ml) and nucleic staining with Syto 13 (50 μM) (Thermo Fisher Scientific). Fluorescence images for CD45 and eGFP staining were acquired using the scanner of GeoMx (Nanostring). For assessing tertiary lymphoid structure (TLS) formation, consecutive tissue sections were stained overnight at 4°C with the primary antibodies anti-CD19 (20 μg/ml), anti-CD4 (0.7 μg/ml), anti-CD8 (0.5 μg/ml), anti-PDPN (20 μg/ml) and visualized with secondary goat anti-Rabbit IgG-AF568 for CD8 and PDPN or tyramide signal amplification (Thermo Fisher Scientific) for CD4 and CD19 following the manufacturer's recommendations. Sections were mounted with Fluoromount-G mounting media containing DAPI (Thermo Fisher Scientific). For immunohistochemistry analysis of spheroids, tissues were fixed with 4% PFA and embedded in 1.7% agarose gel. Paraffin embedding, cutting under microscope control, and antibody staining with H&E, Ki67, vimentin, FAP (R&D Systems), EGFR (DBS), and cytokeratin (Dako) was performed at Sophistolab AG. Image acquisition for spheroids and TLS structures in tumor tissue was performed with a Leica DMi8 microscope (Leica) using a DMC4500 Camera with a 10x or 40x objective.

**Tumor and lymph node dissociation**

Established tumors and lymph nodes were isolated at indicated time points by excision and chopped into pieces of 2 mm diameter for dissociation. To generate single cell suspensions, tumor tissue was further mechanically and enzymatically processed for 30-60 minutes at 37°C using a digestion solution of 1 mg/ml Collagenase D (Sigma-Aldrich) and 10 μg/ml DNase I (Sigma-Aldrich) in RPMI complete medium while shaking. The resulting cell suspension was passed through a 70 μm filter and divided for flow cytometry analysis using antibody staining panels. For analysis of lymphoid cells in tumor-draining lymph nodes (tdLN) and non-draining lymph nodes (ndLN), tissues were smashed with a plunger against a 50 μm cell strainer and collected in FACS buffer for flow cytometry analysis.

**Immunophenotyping of tumors and lymph nodes**

For YUMM1.7 tumors established with PIB-eGFP- or eGFP-transduced cell mixtures and treated with anti-PD-1 or isotype control, tumors and tdLN were isolated for immunophenotyping at day 21 post tumor establishment. For B16 tumors treated with Ad-PIB gene therapy and anti-PD-1 and anti-CTLA4, tumors, tdLN and ndLN were isolated 9 days after first intratumoral Ad-PIB treatment and 16 days post tumor establishment. Tumors and lymph nodes were dissociated as described previously and stained for flow cytometry analysis using the following staining panels: the lymphoid panel included antibodies for CD45, CD3, CD8, CD4, NK1.1, CD49b, CD19, CD44, CD62L, PD-1, CD25, Ki67, T-bet and TCF-1. The myeloid panel included antibodies for CD45, CD11c, MHC-II, F4/80, Ly6C, XCR1, CD11b, Ly6G, SIRPα and Siglec-H. Lymphoid populations were defined according to: $CD8^+$ T cells ($CD45^+CD3^+CD8^+$), $CD4^+$ T cells ($CD45^+CD3^+CD4^+$), NK cells ($CD45^+CD49b^+$), B cells ($CD45^+CD19^+$), effector memory T cells ($CD45^+CD3^+CD44^+CD62L^-$ $CD8^+$ or $CD4^+$), central memory T cells ($CD45^+CD3^+CD44^+CD62L^+$ $CD8^+$ or $CD4^+$), proliferative T cells ($CD45^+CD3^+Ki67^+$ $CD8^+$ or $CD4^+$ T cells), stem-like T cells ($CD45^+CD3^+TCF\text{-}1^+$ CD8+ or $CD4^+$ T cells), regulatory $CD8^+$ T cells ($CD45^+CD3^+CD8^+CD44^+CD25^+$), Treg ($CD45^+CD3^+CD4^+CD25^+$), exhausted T cells ($CD45^+CD3^+PD\text{-}1^+T\text{-}bet^+$ $CD8^+$ or $CD4^+$), terminally exhausted T cells ($CD45^+CD3^+PD\text{-}1^+T\text{-}bet^-$ $CD8^+$ or $CD4^+$), effector T cells ($CD45^+CD3^+PD\text{-}1^-T\text{-}bet^+$ $CD8^+$ or $CD4^+$) and T helper cells

($CD45^+CD3^+$T-bet$^+CD4^+$). Myeloid populations were defined according to: Neutrophils ($CD45^+$MHC-II$^-$$CD11b^+Ly6G^+$), monocytes ($CD45^+$MHC-II$^-$$CD11b^+Ly6C^+$), macrophages ($CD45^+$MHC-II$^+$$CD11c^+F4/80^+$), cDC1 ($CD45^+$MHC-II$^+$$CD11c^+F4/80^-$$SIRPa^-$Siglec-H$^-$$XCR1^+$), cDC2 ($CD45^+$MHC-II$^+$$CD11c^+F4/80^-$$SIRPa^+$Siglec-H$^-$$XCR1^-$) and pDC ($CD45^+$MHC-II$^+$$CD11c^+F4/80^-$Siglec-H$^+$).

**Persistence and migration of transduced cells**

For assessing the persistence of transduced tumor cells in tumors and migration to tdLN, YUMM1.7 cells expressing mCherry (YUMM1.7-mCherry) were transduced with PIB-eGFP or eGFP, mixed 1:1 with parental cells and injected subcutaneously in CD45.1 mice. At days 1, 2, 3, 4, 9 and 15 after tumor establishment, tumors and tdLN were isolated and dissociated into single cell suspensions as described above for flow cytometry analysis. Transduced YUMM1.7 cells were quantified by the percentage of GFP-expressing cells gated in live mCherry+ cells and $CD45.1^-$ cells to exclude host immune cells.

**Immune cell depletion**

For depletion of $CD8^+$ T cells, $CD4^+$ T cells or NK cells, mice were injected intraperitoneally with 100 μL of depleting anti-CD8a (clone 53-6.7, BioXCell), anti-CD4 (clone GK1.5, BioXCell), anti-NK1.1 (clone PK136, BioXCell) or rat isotype control IgG2a (clone 2A3, BioXCell) and IgG2b (clone LTF-2, BioXCell) antibodies (400 μg/mouse). Antibodies were injected two days before, the same day, and two days after tumor establishment. Depleting antibody injections were continuously repeated every 3-4 days. Depletion was confirmed by flow cytometry analysis of NK cells, $CD8^+$ and $CD4^+$ T cells from peripheral blood at the same day of tumor establishment.

**Analysis of single cell RNA sequencing with TCR enrichment of T cells**

Single cell RNA-seq libraries were prepared using the Chromium Single Cell V(D)J Reagent Kit (10x Genomics). Indexed sequencing libraries were quantified with a High Sensitivity DNA analysis kit (Agilent) and Agilent Bioanalyzer. Indexed libraries were pooled in an equimolar ratio and sequenced with an Illumina NextSeq 500 platform. 10x Genomics cellranger Single Cell Software v7.1.0 was used for demultiplexing, alignment (mm10), filtering, UMI counting, single cell 5′ end gene counting, TCR assembly, annotation of paired VDJ and performing quality control using the manufacturer's parameters. The sparse expression matrix generated by the cellranger analysis pipeline was used as input to the Seurat library v4.3.0, and cells and genes passing quality control thresholds were included according to the following criteria: 1) number of genes detected in each single cell greater than 200 and lower than 5,000; 2) percentage of counts in mitochondrial genes less than 7.5%. We normalized data using "LogNormalize" with the scale factor of 10,000 and identified 2,500 variable features. We used the first 30 principal components for subsequent UMAP and clustering analyses. Next, the data was segregated into CD4 and CD8 dataset, according to gene expression of *Cd4* and *Cd8a*. Each data set was reanalyzed independently using 2,000 variable features and 15 principal components for UMAP and clustering analysis. We identified 9 clusters for $CD8^+$ and 12 clusters for $CD4^+$ T cell datasets. For differential expression analysis between cell types, we used FindAllMarkers function with Wilcox Rank Sum test with the following parameters: logfc.threshold = 0.25, min.pct = 0.25. In addition, scaling was performed and resulting genes were visualized using DoHeatmap function and used for annotation of cell types. To calculate overrepresentation of T cell subsets in a particular treatment condition we used the exact binomial test. Additionally, we performed differential expression analysis between differently treated cells for each cluster using the parameters defined above. To order cells on pseudotime, we used Monocle 3 library v1.3.1. The data was separated by treatment and run on corresponding UMAP with default settings except for the following parameter: use_partition=FALSE that assumes all cells in the dataset descend from a common transcriptional ancestor. The root of the trajectory was selected automatically among early activated cells for CD4 and CD8 cells. The VDJ data generated by the cellranger analysis pipeline was used as input to scRepertoire library v1.8.0 and matched with corresponding Seurat objects. We removed any chain without values and multi chains by selecting the 2 corresponding chains with the highest expression for a single cell barcode.

**Cytotoxicity assays with $CD4^+$ T cells**

$CD4^+$ T cells were isolated from tumors established by subcutaneous injection of PIB-eGFP or eGFP-transduced and untransduced YUMM1.7 cells (1:1 ratio) at day 21 post implantation. Tumor tissue was isolated and digested into single cell suspensions. $CD4^+$ T cells were isolated using the mouse $CD4^+$ T cell isolation kit (Miltenyi) according to the manufacturer's instructions. 24 hours before co-culture, $1x10^4$ target tumor cells were seeded on tissue culture 96-well plates in quadruplicates. Target cells were either PIB-eGFP or eGFP-transduced YUMM1.7 cells mixed with untransduced cells (1:1) at day 9 of reprogramming. In addition, PIB-eGFP or eGFP-transduced non-target LLC cells mixed with untransduced cells were included to control for unspecific killing. Isolated live $CD4^+$ T cells were counted and added to target cells in a 1:1 target to effector ratio and co-cultured for 24 hours. Apoptotic and dead cells were quantified by flow cytometry after staining with the Annexin V kit. Apoptotic cells were defined as $annexin^+7\text{-}AAD^-$ and dead cells as $annexin^+7\text{-}AAD^+$ cells and quantified by gating in transduced ($eGFP^+$) and untransduced ($eGFP^-$) cells.

**Spheroid formation**

Human cancer cell line-derived spheroids were formed using the forced-floating method. Each well of an ultra-low-attachment U-bottom 96-well plate (Perkin-Elmer) was seeded with $2x10^4$ cells in 200 µL of respective medium supplemented with 2.5% Matrigel (Corning) followed by centrifugation at 1,000 g for 10 minutes to force cell aggregation. The plates were incubated at 37°C for 96 hours after which tumor spheroid formation can be observed. To form the multi-component heterotypic spheroids, eGFP-transduced T98G (T98G-$eGFP^+$) cells and cancer-associated fibroblasts (CAFs), MDSCs or pericytes were counted, combined at desired ratios, and

$2x10^4$ cells were seeded as described above. 100 μL of medium containing 2.5% Matrigel was replaced every 3-5 days of the experiment. 50% of media was carefully replaced to avoid organoid disruption. For immunohistochemistry, confocal imaging, ATP release assay, and co-cultures with PBMC, the spheroids were aggregated from a suspension of 300 cancer cells and 1,000 CAF over a period of 3 days without the addition of Matrigel. Alternatively, for human cancer cell line-derived spheroid co-cultures with PBMCs, spheroids were formed with MACS-enriched $CD45^+$ T98G cells at day 7 of reprogramming and mixed with CAFs before addition of PBMCs. For formation of patient-derived cancer spheroids, cancer cells were resuspended in 200 µl of DMEM or PneumaCult complete medium containing 2.5% Matrigel and dispensed to single wells of a 96 Thermo Scientific Nunclon Sphera 3D culture system plate. When indicated, patient-derived cancer spheroids were formed with CAFs. Prior to co-culturing CAFs with cancer cells, CAFs were transduced with the lentiviral vector SFFV-mOrange to seperate CAFs based on fluorescent mOrange expression from cancer cells. Spheroids were kept in these conditions for a maximum of 3 days before viral transduction.

**Myeloid-derived suppressor cell differentiation**

To prepare MDSCs, we first isolated PBMCs from peripheral blood of healthy donors using Lymphoprep tubes (Thermo Fisher Scientific). Erythrocytes were removed using Pharm Lyse (BD Biosciences) and monocytes were enriched via positive selection using CD14 microbeads (Miltenyi) by magnetic-activated cell sorting (MACS) according to the manufacturer's protocol. $CD14^+$ monocytes were cultured in RPMI complete supplemented with 1 µg/ml of TLR2 agonist Pam3CSK4 (1 µg/ml; Invivogen) previously shown to promote MDSC polarization (*56*). Acquisition of an MDSC phenotype was confirmed via flow cytometry quantification of the markers CD163, CD206, and 25F9.

**ATP Release Assay**

Viability and growth of spheroids was measured the day after aggregation (day 0) and after 9 days in culture using the CellTiter-Glo 3D cell viability assay (Promega), based on the evaluation of ATP luminescence. Serial dilutions of the ATP-Standard (Sigma FLAAS-1VL) were performed to generate the standard curve. The luminescence readout was acquired using Spark Multimode Microplate Reader (TECAN Life Sciences).

**Evaluation of reprogramming efficiency using confocal microscopy**

After 9 days of reprogramming, human cancer cell line-derived spheroids were fixed with 4% PFA, permeabilized with 0.5% Triton X-100 in PBS and blocked with Normal Serum Block (Biolegend). Spheroids were then incubated overnight with anti-CD45 (Abcam) or anti-HLA-DR (Biolegend) primary antibodies in incubation buffer (PBS supplemented with 10% FBS and 0.2% Triton X-100). The following day, spheroids were incubated with Cy5-conjugated goat anti-mouse IgG (Abcam) secondary antibody and DAPI in the antibody buffer for 6 hours. Finally, clearing with the ScaleS4 solution (PerkinElmer) was performed. High-content confocal image acquisition was performed using the Confocal Quantitative Image Cytometer CQ1 (Yokogawa). Images were further analyzed using the CellPathfinder (Yokogawa) high content analysis software through the identification and quantification of individual nuclear and membrane staining. The size of spheroids was determined through the detection of DAPI fluorescence and processed with the CellPathfinder software. The analysis was performed using images of the middle section of the spheroids.

**Single cell RNA sequencing of human cancer cells reprogrammed in 2D and 3D**
scRNA-seq was performed in T98G cells from monolayer culture or isolated from spheroids after the transduction with PIB-eGFP or control vector eGFP. At day 3, 7 and 9 of reprogramming, 5,000 to 10,000 transduced eGFP$^{+}$ cells expressing CD45 and/or HLA-DR, were FACS-purified and resuspended in PBS containing 0.04% bovine serum albumin (BSA). Day 0 controls were purified from eGFP-transduced cultures based on eGFP positivity at day 9 and processed similarly. Cells were loaded on a 10x Chromium (10x Genomics) without multiplexing. Single cell RNA libraries were prepared using the Chromium Single Cell 3$^{+}$ v2 Reagent Kit (10x Genomics). Indexed sequencing libraries were quantified with a High Sensitivity DNA analysis kit (Agilent) and Agilent Bioanalyzer. Indexed libraries were pooled in an equimolar ratio and sequenced with an Illumina NextSeq 500 platform.

**Analysis of single cell RNA sequencing of human cancer cells reprogrammed in 2D and 3D**
10x Genomics cellranger Single Cell Software v7.1.0 was used for demultiplexing, alignment (hg38), UMI counting and single cell 3′ end gene counting. The sparse expression matrix generated by cellranger analysis pipeline was used as input to Seurat library v4.3.0, and cells and genes passing quality control thresholds were included according to the following criteria: 1) total number of unique molecular identifiers detected per sample greater than 3 lower median absolute deviations (MADs); 2) number of genes detected in each single cell greater than 3 MADs; 3) percentage of counts in mitochondrial genes less than 10%. We normalized data using "LogNormalize" with the scale factor of 10,000 and identified 7000 variable features. For differential expression analysis between day 0 and corresponding days, we used FindMarkers function with Wilcox Rank Sum test with the following parameters: logfc.threshold = 0.1, min.pct = 0.25. Gene ontology (GO) and Reactome pathway enrichment analyses were performed using enrichR library v3.1 (https://cran.rproject.org/web/packages/enrichR/index.html). We calculated the percentage of overlapping genes between a tumor-APC signature gene list established previously (*23*) and differentially expressed genes in 2D and 3D settings. We used scPred with default svm radial model and publicly available DC single cell expression data for DC subset affiliation (*54, 57*). Training of classifiers with available DC data was performed using 7,000 variable genes. We then classified normalized expression levels from the reprogramming dataset. We used a probability threshold of 0.95 to classify cells into classes. GSEA was performed using singleseqgset R library (https://arc85.github.io/singleseqgset/) with default parameters. The immunogenic and tolerogenic gene signatures were based on combination of genes in Cluster 49

and Cluster 61 (TLR-induced maturation) and Cluster 103 (Homeostatic maturation) previously defined (*32*). The resulting enrichment scores were scaled in each sample individually. Additionally, GSEA between reprogrammed cells in spheroids at day 9 and non-reprogrammed cells (transduced with eGFP) was performed against IFN-γ, STING, TLR pathways, which were curated based on human KEGG pathways and NF-kB signaling pathway from Biocarta. The analysis was performed using UMI counts with default parameters, P = 1 for calculation of enrichment statistics, normalized enrichment scores and rank by calculating difference of means scaled by the standard deviation.

**Spheroid co-cultures with PBMC**

Donor-derived human PBMCs (STEMCELL Technologies) were resuspended in RPMI complete medium at the concentration of $1x10^5$ cells/ml. When stated, medium was supplemented with IL-2 (50 U/ml; Peprotech) and IL-7 (10 ng/ml; Peprotech) to maintain T cells in culture, or with a combination of anti-CD3 (125 ng/ml; Biolegend) and anti-CD28 (250 ng/ml; Biolegend) to induce T cell pre-activation. Subsequently, $5x10^3$ non-activated or pre-activated PBMCs were added to the each well containing eGFP$^+$ T98G spheroids aggregated with reprogrammed CD45$^+$ or mCherry-transduced T98G-eGFP$^+$ cells mixed with CAFs. T98G cells were HLA typed by antibody staining as HLA-A2 positive (*23*). PBMCs from donors were either HLA-A2 positive (matched) or negative (unmatched). PBMCs and the T98G cancer cell line were HLA-matched. Changes in size of spheroids were evaluated after 9 days of co-culture with PBMCs by assessing eGFP fluorescence using Spark Multimode Microplate Reader (TECAN Life Sciences). Using PBMCs in co-culture with spheroids containing reprogrammed or eGFP-transduced T98G-eGFP$^+$ cells mixed with CAFs, the supernatant was collected and processed with LEGENDplex Human CD8/NK Panel kit (Biolegend) according to the manufacturer´s protocol. Cytokine concentrations were determined using the LEGENDPlex Data Analysis software (Biolegend).

**Tumor spheroid clearing and staining for light sheet microscopy**

Spheroids were fixed by overnight incubation with 200-600 μL of 4% PFA and then blocked by overnight incubation with blocking solution (PBS with 0.2% Triton-X, 5% BSA, 10% DMSO). Incubation with primary antibody (Rabbit Anti-eGFP antibody - abcam290; 1/400) was performed overnight in PBS solution containing 0.2% Tween, 0.01 mg of of heparin, 5% DMSO and 3% BSA, washed 4-5 times in PBS supplemented with 0.02% Tween and 0.01 mg of heparin (PTwH buffer) and kept overnight in PTwH buffer. Incubation with secondary antibody (Goat anti-Rabbit Alexa Fluor 647, Invitrogen; 1/400) was performed overnight in PTwH buffer supplemented with 3% BSA, washed 4-5 times in PTwH buffer and kept overnight in the same buffer. Then, spheroids were casted in 1% low gelling temperature agarose (Merck – A9414), dehydrated by sequential incubations in methanol (MeOH)/PBS solutions containing 20, 40, 60 and 80% MeOH for 1 hours, followed by overnight incubation in 100% MeOH. On the following day, dehydrated spheroids were incubated in 67% DCM (Dichloro methane, Sigma-Aldrich, 270997-100ML) and 33% MeOH for 3 hours with shaking, followed by 2 incubations of 15 minutes with 100% DCM with shaking to wash away the MeOH. Next, spheroids were incubated in ethyl cinnamate (Sigma-Aldrich, 112372) overnight and fresh ethyl cinnamate was added for another 24 hours to prevent sample oxidization. All steps were performed at room temperature. Imaging was performed using an Ultramicroscope Blaze light sheet microscope (Miltenyi).

**Patient-derived cancer cell co-cultures with HLA-matched T cells**

Patient-derived melanoma cells transduced with Ad-PIB-eGFP or Ad-eGFP in monolayer or spheroids at reprogramming day 9 were used for co-culture experiments. For 2D cultures, transduced cells were collected in 5 ml tubes, resuspended to a concentration of $0.5x10^6$ cells per ml of RPMI complete medium supplemented with 1 mM of MART1 ELAGIGILTV (JPT, SP-MHCI-0006) and CMV peptide NLVPMVATV (JPT SP-MHCI-0005) and incubated for 3 hours at 37°C and 5% $CO_2$. Purified HLA-A2-matched $CD8^+$ T cells were added to a 48-well plate and peptide-loaded cells were washed in complete RPMI and gently loaded on top of $CD8^+$ T cells to a final ratio of 1 transduced melanoma cell to 10 $CD8^+$ T cells in a final volume of 500 µl. For spheroids, medium was replaced by fresh RPMI media containing 1 mM of MART1 ELAGIGILTV and CMV NLVPMVATV peptides, and spheroids were incubated for 3 hours at 37°C and 5% CO2. Peptide-loaded spheroids were then gently washed in complete RPMI and $0.5x10^6$ $CD8^+$ T cells loaded on top using a final volume of 200 µl. After 3 and 5 days of co-culture, culture medium was replaced by fresh complete RPMI medium containing 50 U/ml of IL-2 and 10 µg/ml of IL-7. After 8 days of co-culture, cells from 2D and 3D cultures were processed for flow cytometry analysis.

**ELISA quantification of autoantibodies**

Plasma was collected from 3 survivor mice at day 210 (30 weeks) post subcutaneous B16 tumor establishment and subsequent treatment with Ad-PIB and ICB (anti-PD-1 and anti-CTLA4) and 3 naïve control mice (21 weeks) for the quantification of autoantibodies. One of the survivor mice showed vitiligo. Blood was isolated by tail vein puncture into K2-EDTA coated microvette tubes (Sarstedt) and centrifuged at 1,000 g for 15 minutes to remove cellular components and collect plasma. Levels of autoantibodies in the plasma were then quantified using the anti-SSA, anti-SSB, anti-CENP-B and anti-dsDNA ELISA kits (Signosis) following manufacturer's instructions. In brief, a standard curve for each test was generated ranging from 250-3.9 ng/ml for anti-SSB and anti-CENP-B and from 125-1.9 ng/ml for anti-dsDNA and anti-SSA. The read-out was performed using the GloMax Discover microplate reader (Promega) at 450 nm absorbance.

**Histological evaluation of organs**

Mouse brain, heart, lung, kidney, colon, and stomach were excised from the same 6 mice and at the same time-point as for quantification of autoantibodies. Tissues were fixed, dehydrated, and embedded in paraffin after xylene treatment as described above. 4 µm thick tissue sections were cut and mounted onto BOND Plus slides (Leica). The paraffin-embedded sections were de-paraffinized and stained with H&E. The histological evaluation was conducted by ImaGene-iT AB. Organ histology was examined using H&E-stained sections under a bright-field microscope

(Olympus IX70). Assessment included scoring the presence of recruited immune cells and other atypic findings for each individual animal. Severity of findings was graded on a scale from none to high (0-3), and the distribution of pathology within sections was quantified as a percentage. Severity scores were then multiplied by the percentage of pathology distribution, resulting in scores ranging from 0 to 300. For each animal, four kidney sections from both the left and right kidneys, and two sections from the brain, heart, lung kidney, colon, and stomach were examined. Representative organ images were generated by slide scanning entire sections (Hamamatsu S210, Hamamatsu, Japan) from which low and high magnification digital images were captured using the ndpi software (Ndp viewer, Nanozoomer, Hamamatsu).

### Supplementary Figures

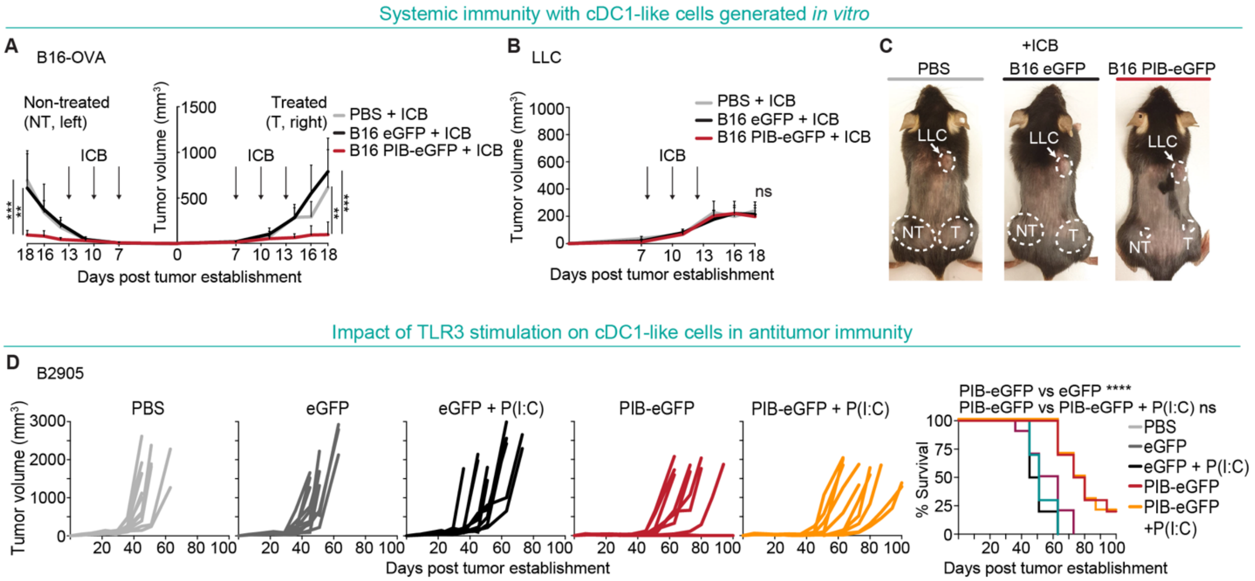


**Figure S1. Reprogrammed cancer cells *in vitro* elicit systemic antitumor immunity.**

**(A-C)** Bilateral B16-F10 (B16) expressing ovalbumin (B16-OVA) murine melanoma and heterologous Lewis lung adenocarcinoma (LLC) tumors were established by subcutaneous injection. B16-F10 cells (B16, not expressing OVA) were transduced with lentiviral vectors encoding PU.1, IRF8, BATF3 (PIB) and reprogrammed for 5 days *in vitro*. $1x10^5$ MACS-enriched B16-derived cDC1-like cells were then injected intratumorally at day 7, 10 and 13 into B16-OVA tumors into the right flank after overnight incubation with full-length OVA protein and toll-like receptor (TLR) 3-agonist polyinosinic-polycytidylic acid (P(I:C)). Additionally, immune checkpoint blockade (ICB, anti-PD-1 and anti-CTLA-4) or isotype control antibodies (IgG2a and IgG2b) were administered intraperitoneally at day 7, 10 and 13. Arrows indicate time-points of injection. Intratumoral injections of eGFP-transduced cells or PBS were included as controls (n=10). (A) Tumor growth of bilateral B16-OVA tumors and (B) control LLC tumors within the same animals. (C) Representative animals with bilateral B16-OVA (treated and non-treated) and control LLC tumors showing abscopal effects for melanoma but not LLC tumors. Arrows indicate tumor locations and dashed lines tumor sizes. **(D)** B2905 murine melanoma tumors were established subcutaneously and $1x10^5$ MACS-enriched B2905-derived cDC1-like cells were injected intratumorally at day 7, 10 and 13. 24 hours before intratumoral injection, cells were stimulated with P(I:C) or left unstimulated. Injections of eGFP-transduced B2905 cells stimulated with P(I:C) or unstimulated, or PBS were included as controls. Tumor growth (left) and survival (right) of mice are shown (n=10). Data in panel A and B are shown as mean ± SD. Comparisons in panel A and B were analyzed using the Kruskal-Wallis test followed by Dunn's multiple comparison test. Survival analysis in panel D was performed by log-rank Mantel-Cox test. ns - non-significant, $^{**}p<0.01$, $^{***}p<0.001$, $^{****}p<0.0001$.

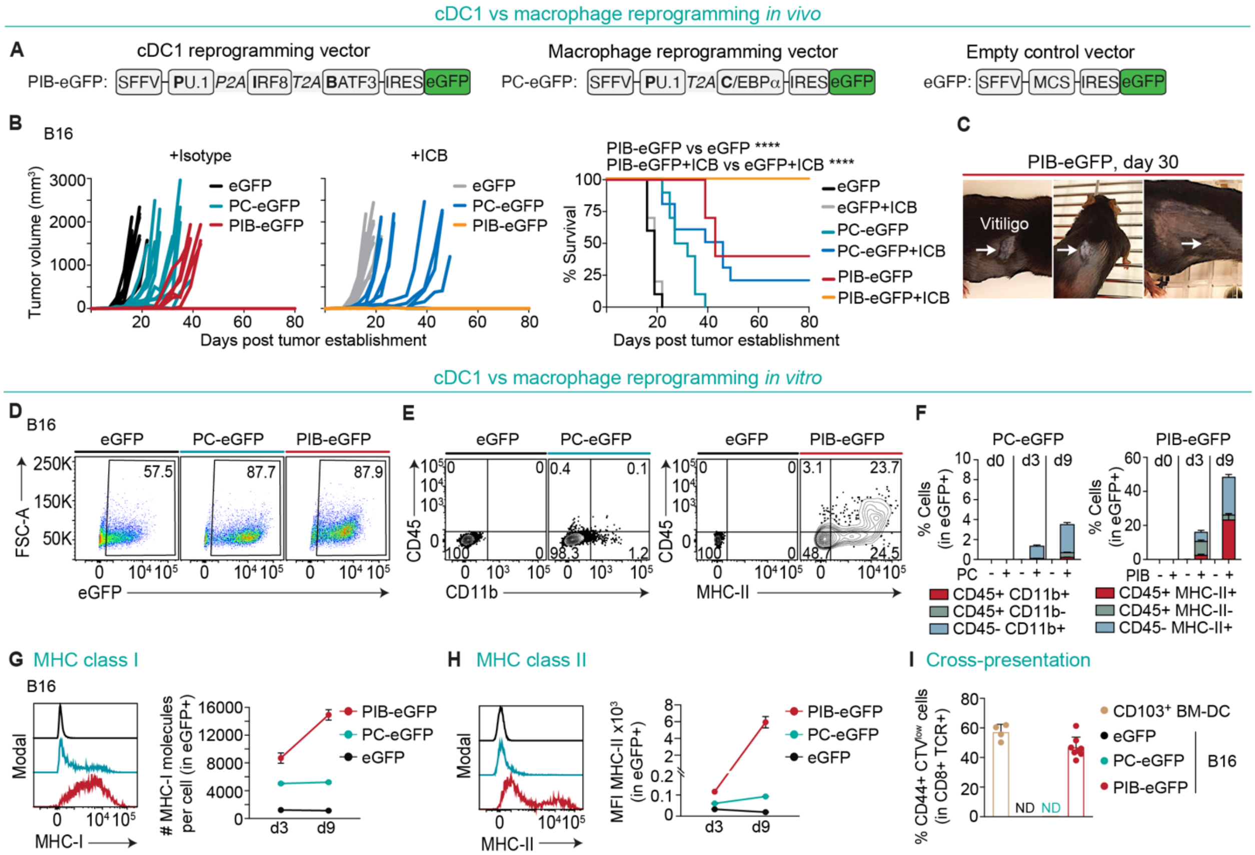


**Figure S2. *In vivo* cDC1 reprogramming elicits potent antitumor immunity.**

**(A)** Lentiviral vectors encoding PIB-eGFP to induce cDC1 reprogramming or PU.1 and C/EBPα (PC) to induce macrophage reprogramming under a constitutive SFFV promoter followed by an IRES-eGFP were used in B16 murine melanoma cells. Transcription factors were separated by 2A sequences. Empty vector containing a multiple cloning site (MCS) and IRES-eGFP was used as transduction control. **(B-H)** 88% of B16 cells were transduced with lentiviral vectors encoding PIB-eGFP or PC-eGFP *in vitro*, mixed with 12% of parental B16 cells and a total of $1x10^5$ cells per mouse were injected subcutaneously to induce tumor cell reprogramming *in vivo* along with tumor establishment. Injection of eGFP-transduced cells mixed with parental cells were included as controls. Additionally, cell mixtures were maintained *in vitro* to quantify transduced cells at day 3 and reprogramming efficiencies until day 9. ICB (anti-PD-1 and anti-CTLA-4) or isotype control (IgG2a and IgG2b) antibodies were administered intraperitoneally at day 7, 10, and 13 post tumor establishment. (B) Tumor growth (left) and survival of mice (right) are shown (n=10). (C) Representative animals 30 days after receiving PIB-eGFP-transduced B16 cells showing vitiligo (arrow) on tumor regression site. (D) Flow cytometry analysis of PIB-eGFP, PC-eGFP or eGFP-transduced cells 3 days after transduction and mixing with untransduced cells in parallel *in vitro*. (E) Flow cytometry analysis and (F) quantification of macrophage reprogramming efficiency measured by CD45 and CD11b marker expression (left) and cDC1 reprogramming efficiency measured by CD45 and MHC-II marker expression (right) at the time point of subcutaneous implantation (16 hours after transduction (d0)) and after 3 (d3) and 9 days (d9) of reprogramming *in vitro*. Reprogramming efficiency was quantified within transduced $eGFP^+$ cancer cells (n=3-9). (G) Flow cytometry analysis and quantification of the number of surface MHC-I molecules per cell and (H) MHC-II expression in B16 cells by mean fluorescence intensity (MFI) after 3 and 9 days of *in vitro* reprogramming (n=3). **(I)** Quantification of proliferating $CTV^{low}CD44^+$ ovalbumin-specific $CD8^+$ T cells (OT-I) after co-culture with $CD103^+$ bone marrow-derived dendritic cells (BM-DC), eGFP-transduced B16 cells (not expressing ovalbumin), MACS-enriched $CD45^+$ and $MHC\text{-}II^+$ B16-derived cDC1-like cells or MACS-enriched $CD45^+$ and $CD11b^+$ B16-derived macrophage-like cells (PC). $CD103^+$ BM-DC, eGFP-transduced or reprogrammed B16 cells were pulsed for 24 hours with full-length ovalbumin protein. To start the co-culture, ovalbumin-containing media was washed away extensively, and fresh media added together with OT-I cells and co-cultures were performed for 72 hours (n=4-7). Data in panel F-I are shown as mean ± SD. Survival analysis in panel B was performed by log-rank Mantel-Cox test. ****p<0.0001.

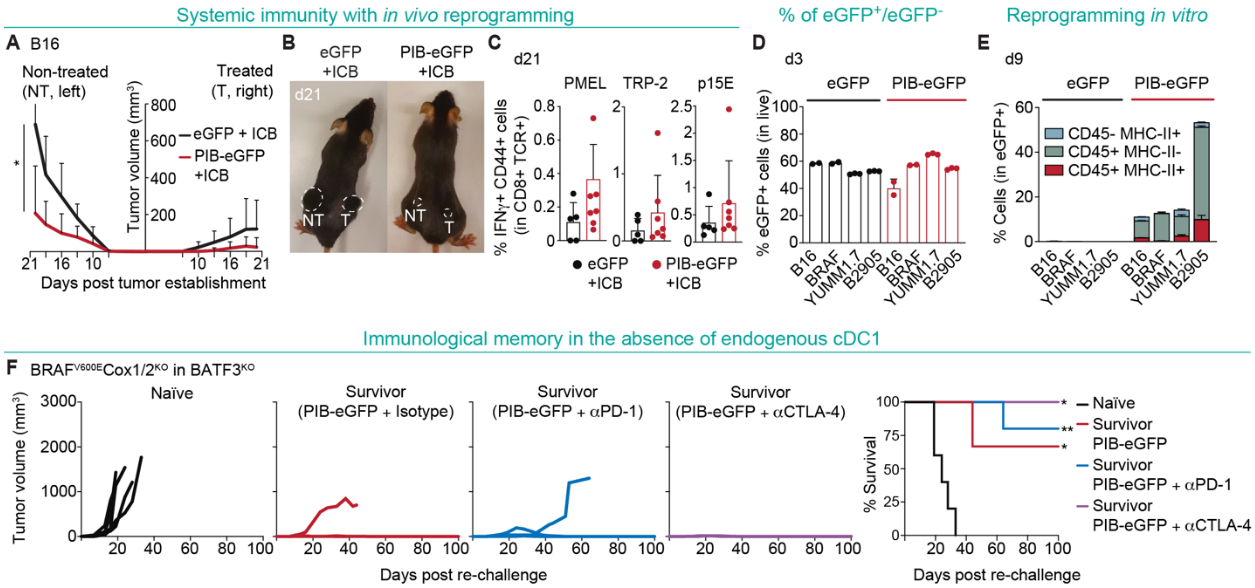


**Figure S3. cDC1 reprogramming *in vivo* induces abscopal effects and immunological memory independently of endogenous cDC1.**

**(A-C)** Bilateral B16 tumors were established by subcutaneous injection of $2x10^5$ PIB-eGFP- or eGFP-transduced B16 cells (1:1 mixtures with parental B16 cells) in the right flank and $1x10^5$ untransduced B16 cells in the left flank. ICB (anti-PD-1 and anti-CTLA-4) were administered intraperitoneally at day 7, 10, and 13 post tumor establishment (n=8). (A) Tumor growth of bilateral B16 tumors (treated and non-treated) is shown. (B) Representative animals with bilateral B16 tumors showing abscopal effects at day 21 when treated with *in vivo* reprogramming. Dashed circles highlight tumor sizes. (C) Flow cytometry quantification of tumor antigen-specific $CD44^+IFN\gamma^+CD8^+$ T cells from peripheral blood at day 21 after *in vitro* re-stimulation with TRP-2-, PMEL- and p15E peptides. **(D, E)** B16, YUMM1.7 and B2905 melanoma cells were transduced with PIB-eGFP- or eGFP-encoding lentiviral particles *in vitro* (1:1 mixture with parental cells) and injected subcutaneously into wild-type (WT) C57BL/6J animals and $BRAF^{V600E}COX1/2^{KO}$ into $BATF3^{KO}$ mice. (D) Flow cytometry quantification of transduced cells by eGFP expression from parallel *in vitro* cultures at day 3 and (E) reprogrammed cells by CD45 and MHC-II surface expression at day 9 (n=3). cDC1 reprogramming efficiency was evaluated by the percentage of $CD45^+$ $MHC\text{-}II^+$ cells (completely reprogrammed) and $CD45^+$ $MHC\text{-}II^-$ or $CD45^-$ $MHC\text{-}II^+$ cells (partially reprogrammed) within $eGFP^+$ transduced cancer cells. **(F)** Survivor cDC1-deficient $BATF3^{KO}$ mice that remained 100 days tumor-free were subcutaneously re-challenged with $1x10^5$ parental $BRAF^{V600E}COX1/2^{KO}$ cells. Age-matched naïve mice were used as controls and tumor growth (left) and survival (right) are shown (n=2-5). Data in panel A and C-E are shown as mean ± SD. Comparisons in panel A were analyzed using the Mann-Whitney test. Survival analysis in panel F was performed by log-rank Mantel-Cox test. $*p<0.05$, $**p<0.01$.

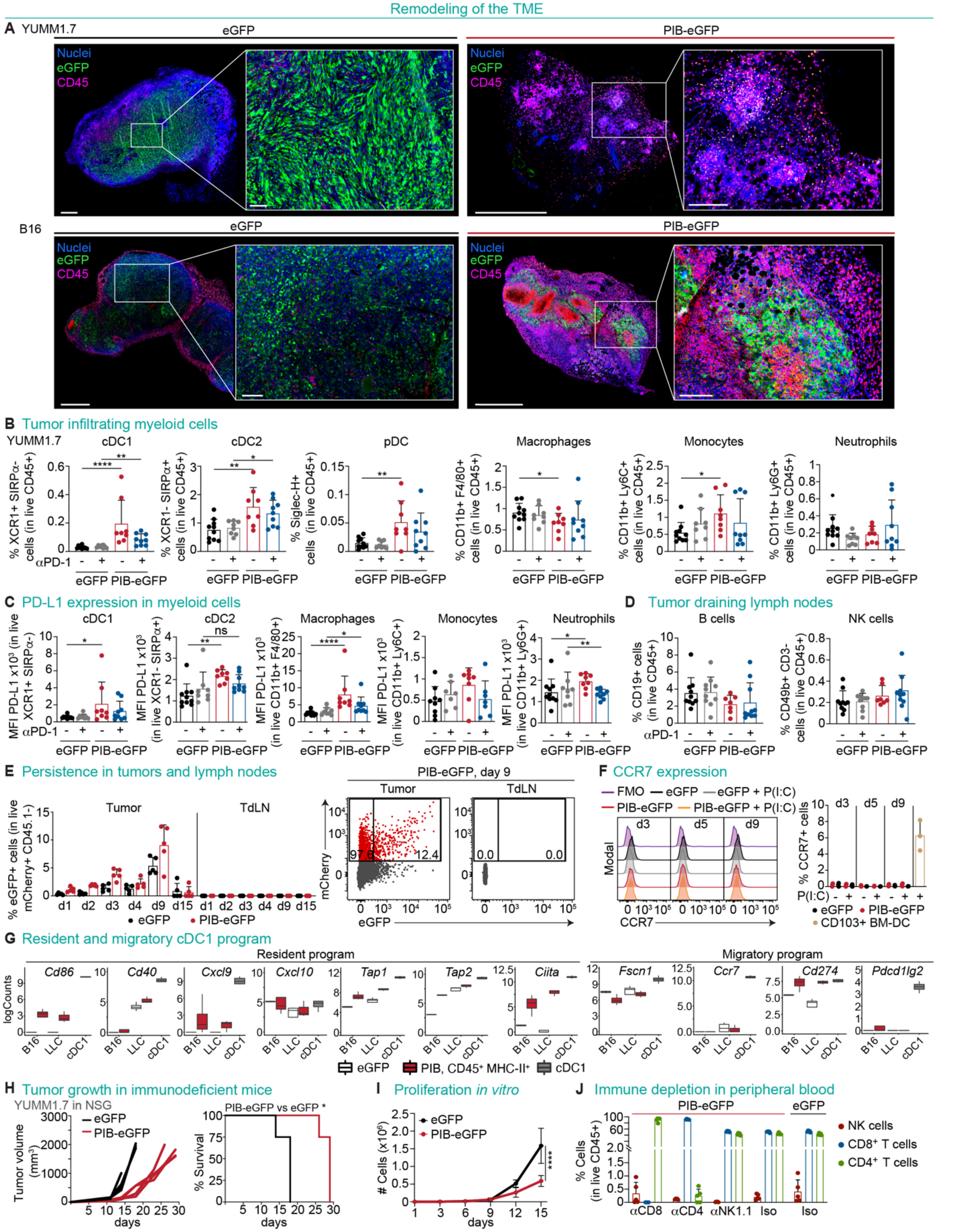


**Figure S4. *In vivo* reprogrammed cells persist in the tumor, acquire a resident cDC1 profile and elicit PD-L1 expression in myeloid cells.**

**(A)** Immunofluorescence analysis of paraffin-embedded YUMM1.7 (upper) and B16 (lower) tumors 9 days after subcutaneous implantation of PIB-eGFP- or control eGFP-transduced cells (both mixed at 1:1 ratio with parental cells). Tumor sections were stained for eGFP (transduced cells), CD45 (immune cells) and nuclei (Syto 13). Scale bars in images showing the whole tumor (left) are 1 mm and scale bars within the magnified regions are 200 µm. **(B-D)** Tumors and tumor-draining lymph nodes (tdLN) were isolated at day 21 after establishment with PIB-eGFP- or eGFP-transduced cells (1:1 ratio with parental cells) for flow cytometry immunophenotyping (n=8-10). (B) Percentages of myeloid cells including $XCR1^+$ cDC1, $SIRP\alpha^+$ cDC2, Siglec-$H^+$ pDC, $F4/80^+$ macrophages, $Ly6C^+$ monocytes, and $Ly6G^+$ neutrophils gated in live $CD45^+$ cells in the tumor. (C) PD-L1 expression in myeloid cells measured by mean fluorescence intensity (MFI). (D) Percentages of B cells and NK cells in tdLN within live $CD45^+$ cells. **(E)** Tumors and tdLN were isolated from CD45.1 mice at days 1, 2, 3, 4, 9 and 15 after establishment with PIB-eGFP- or eGFP-transduced YUMM1.7-$mCherry^+$ cells (1:1 ratio with parental cells) for flow cytometry quantification of transduced cells. The percentages of $eGFP^+$ cells were gated in live $mCherry^+CD45.1^-$ cells (left) (n=5). Representative flow cytometry plots showing $mCherry^+eGFP^-$ and $mCherry^+eGFP^+$ cells in tumors and tdLN gated in live $mCherry^+$ $CD45.1^-$ cells at day 9 (right). (**F**) Quantification (right) and representative histograms (left) for the percentages of $CCR7^+$ YUMM1.7 cells 3, 5 and 9 days of transduction with PIB-eGFP and eGFP as control. Mouse $CD103^+$ bone marrow-derived dendritic cells (BM-DC) are shown as reference. Cells were stimulated for 24 hours *in vitro* with polyinosinic-polycytidylic acid (P(I:C)) before flow cytometry profiling (n=3-9). (**G**) mRNA level of expression of genes associated with a resident (*Cd86, Cd40, Cxcl9, Cxcl10, Tap1, Tap2, Ciita*) or migratory (*Fscn1, Ccr7, Cd274, Pdcd1lg2*) cDC1 program in reprogrammed B16 melanoma or LLC lung tumor cells. Analysis was performed using bulk RNA-seq data (*23*) for eGFP-transduced (white), FACS-sorted $CD45^+$ MHC-$II^+$ B16 and LLC cells after 9 days of reprogramming (red) and splenic cDC1s (gray). **(H)** Immunodeficient NOD.Cg-$Prkdc^{SCID}$ $Il2rg^{tm1Wjl}$/SzJ (NSG) mice were injected subcutaneously with PIB-eGFP- or eGFP-transduced YUMM1.7 cells mixed at a 1:1 ratio with parental cells. Tumor growth (left) and survival (right) are shown (n=4). (**I**) *In vitro* proliferation of PIB-eGFP- or eGFP-transduced YUMM1.7 cells mixed 1:1 with parental cells. 10,000 cells were seeded at day 0 and live cell counts were performed using an automated cell counter at days 1, 3, 6, 9, 12 and 15 (n=9). **(J)** C57BL/6J mice were depleted from $CD8^+$ ($\alpha$CD8), $CD4^+$ T cells ($\alpha$CD4), and NK cells ($\alpha$NK1.1) by intraperitoneal injections of depleting or isotype control antibodies at days -2, 0 and 2 and every three days after tumor establishment with a mixture of transduced YUMM1.7 cells. Flow cytometry validation of $CD8^+$ T cells, $CD4^+$ T cells, and NK cell depletion in peripheral blood at the day of tumor establishment. Data in panel B-F and I-J are shown as mean ± SD. Comparisons in panel B-D and I were analyzed using the Mann-Whitney test. Survival analysis in panel H was performed by log-rank Mantel-Cox test. ns - non-significant *p<0.05, **p<0.01, ****p<0.0001.

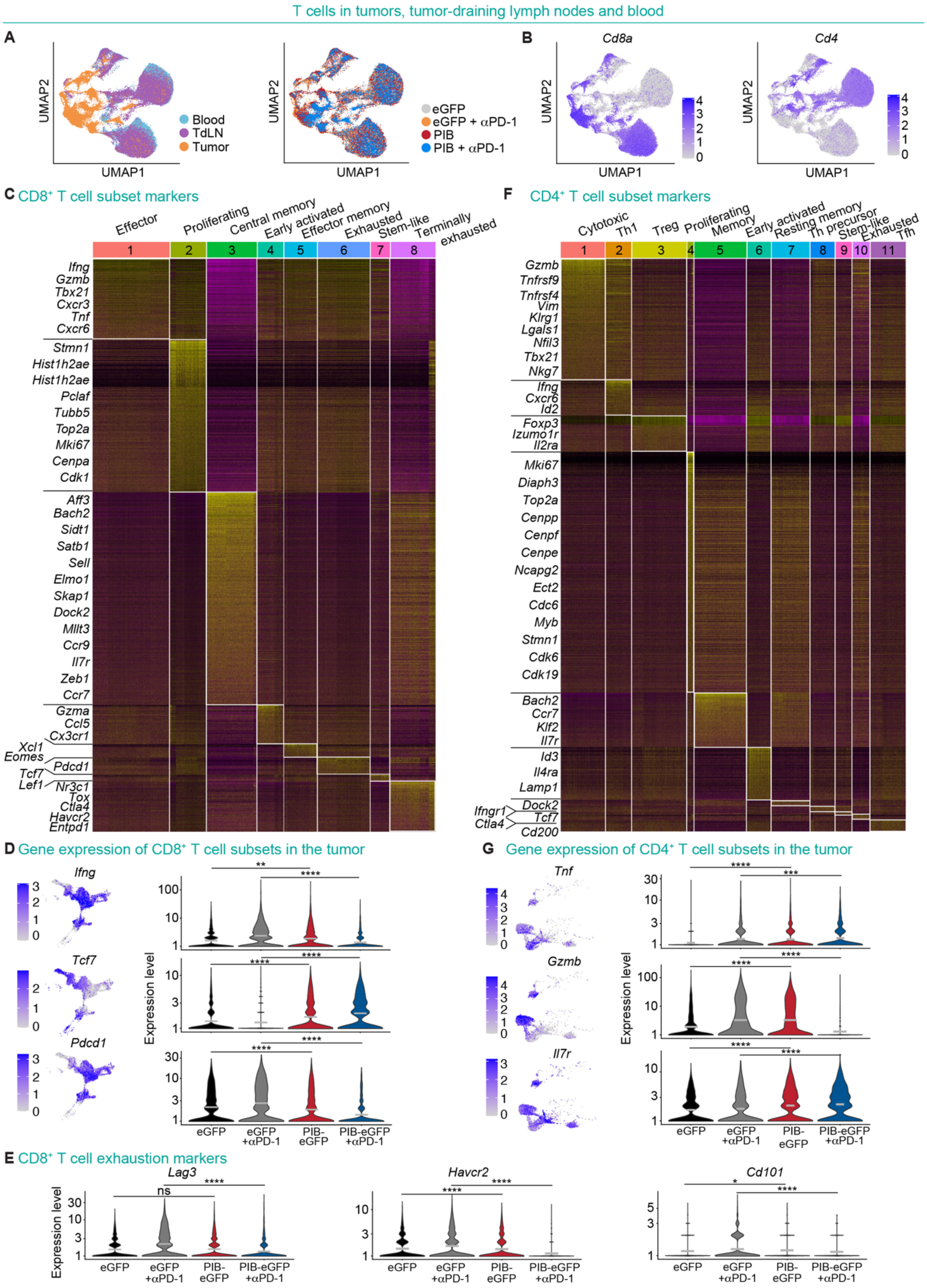


**Figure S5. Annotation of CD8⁺ and CD4⁺ T cell subsets with single cell RNA-sequencing data.**

**(A-G)** YUMM1.7 cells were transduced with PIB-eGFP or eGFP, mixed at a 1:1 ratio with parental cells and injected subcutaneously to establish tumors. Additional groups received anti-PD-1 treatment intraperitoneally at day 7, 10, and 13 after tumor establishment (n=5). Peripheral blood, tumor-draining lymph nodes (tdLN) and tumors were isolated and processed 21 days after tumor establishment to sort $CD45^+$ $CD3^+$ T cells by FACS for 5' scRNA-seq analysis with TCR enrichment. (A) Principal component analysis of $CD8^+$ and $CD4^+$ T cells visualized by Uniform manifold approximation and projection (UMAP) plots from tumors, tdLN and blood (left) across treatment conditions (right). (B) UMAP plots showing *CD8a* and *CD4* gene expression in the whole population of $CD45^+$ $CD3^+$ T cells. (C) Heatmap indicating differentially expressed genes and clusters used to annotate $CD8^+$ T cell subsets. (D) UMAP and violin plots showing the expression of *Ifng, Tcf7* and *Pdcd1* in $CD8^+$ T cells within tumors across treatment conditions. (E) Violin plots showing expression of the $CD8^+$ T cell exhaustion markers *Lag3, Havcr2* and *Cd101*. (F) Heatmap showing differentially expressed genes and clusters used to annotate $CD4^+$ T cell subsets. (G) UMAP and violin plots showing the expression of *Tnf, Gzmb* and *Il7r* in $CD4^+$ T cells in treated and control tumors. In panel D, E and G the mean is indicated by the gray bar. Comparisons in D, E and G were performed using the Mann-Whitney test for eGFP vs. PIB-eGFP or eGFP + anti-PD-1 vs. PIB-eGFP + anti-PD-1. ns - non-significant *$p<0.05$, **$p<0.01$, ***$p<0.001$, ****$p<0.0001$.

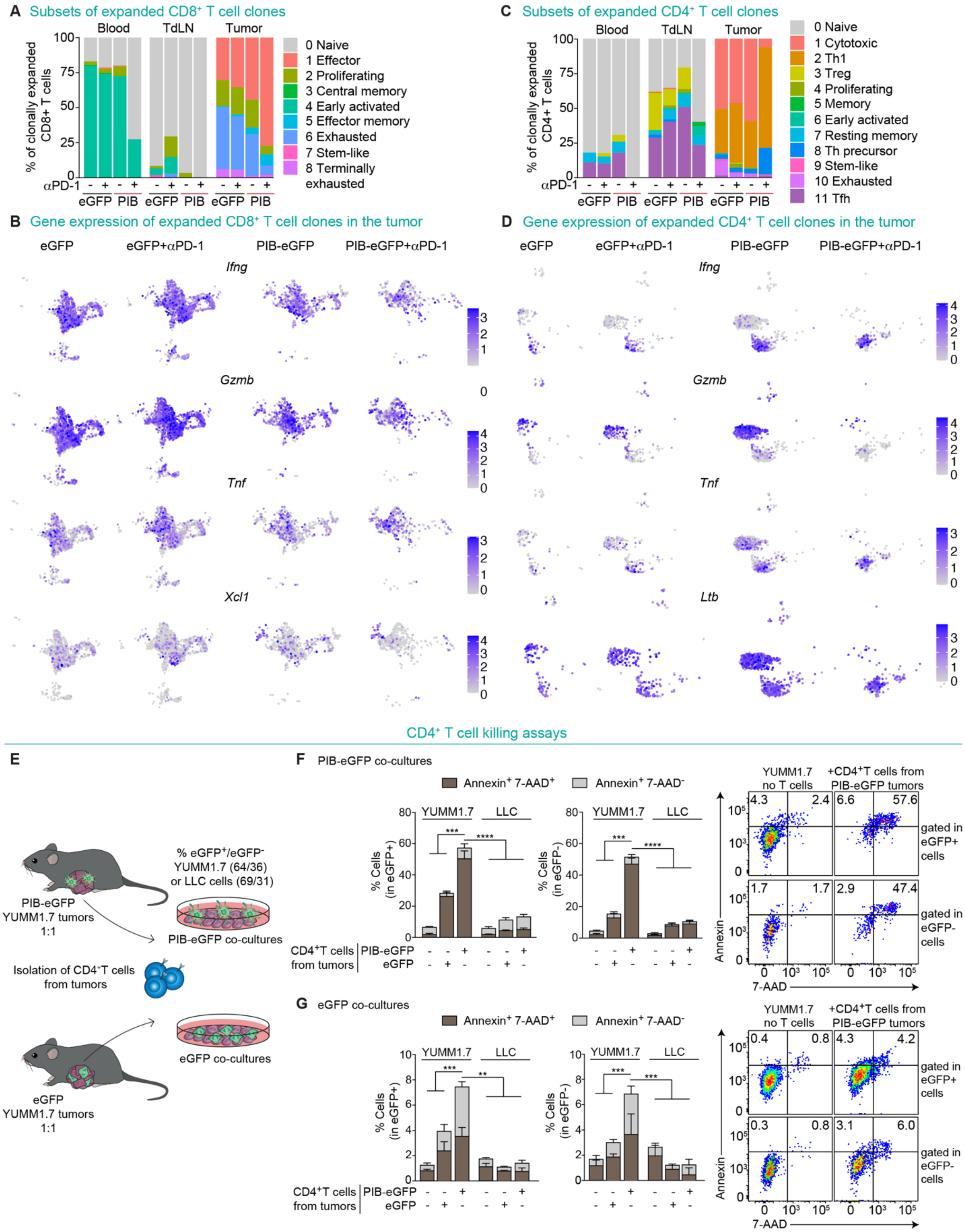


**Figure S6. CD4$^+$ T cells in tumors undergoing reprogramming directly kill melanoma cells. (A-D)** YUMM1.7 cells were transduced with PIB-eGFP or eGFP, mixed at a 1:1 ratio with parental cells and injected subcutaneously to establish tumors. Anti-PD-1 was administered intraperitoneally at day 7, 10, and 13 post tumor establishment (n=5). Peripheral blood, tdLN and tumors were isolated and processed 21 days after tumor establishment and T cells were FACS-purified to perform 5' scRNA-seq with TCR enrichment. (A) Bar plots show the frequencies of expanded clones (>1 cell per clonotype) color-coded by annotation to the CD8$^{+}$ T cell subsets. (B) UMAP plots visualizing gene expression of *Ifng, Gzmb, Tnf,* and *Xcl1* for clonally expanded CD8$^{+}$ T cells in tumors across the four treatment conditions. (C) Bar plots showing frequencies of expanded clones (>1 cell per clonotype) color-coded by annotation to the CD4$^{+}$ T cell subsets. (D) UMAP plots visualizing gene expression of *Ifng*, *Gzmb, Tnf,* and *Ltb* for clonally expanded CD4$^{+}$ T cells in tumors across the treatment conditions. **(E-G)** Experimental strategy for the cytotoxicity assays using CD4$^{+}$ T cells from tumors that were established with PIB-eGFP or eGFP-transduced cells (1:1 mix with parental cells). CD4$^{+}$ T cells were isolated at day 21 after tumor establishment and co-cultured *in vitro* with PIB-eGFP or eGFP-transduced cells mixed with parental cells for 24 hours in a 1:1 effector to target ratio. After 24 hours, killing was assessed by flow cytometry with Annexin V and 7-AAD staining. Co-cultures with PIB-eGFP- or eGFP-transduced non-target Lewis lung carcinoma (LLC) cells were included as controls for the specificity of killing. Percentages of eGFP$^{+}$ and eGFP$^{-}$ cells quantified by flow cytometry are indicated. (F) CD4$^{+}$ T cell mediated killing of PIB-eGFP-transduced cells (eGFP+, left) or untransduced cells (eGFP-, middle). Apoptotic cells (Annexin$^{+}$ 7-AAD$^{-}$) and dead cells (Annexin$^{+}$7-AAD$^{+}$) were quantified. Representative flow cytometry plots (right) from cancer cell cultures without CD4$^{+}$ T cells and co-cultures with CD4$^{+}$ T cells from PIB-treated tumors are shown. (G) Killing of eGFP-transduced cells (eGFP$^{+}$, left) or untransduced cells (eGFP-, middle) and representative flow cytometry plots (right) are shown. In panel F and G mean ± SD is represented (n=4). Comparisons in F and G were performed using the Kruskal-Wallis test by comparing the co-cultures with CD4$^{+}$ T cells isolated from PIB-eGFP mice with YUMM1.7 cells against the other experimental conditions. **p<0.01, ***p<0.001, ****p<0.0001.

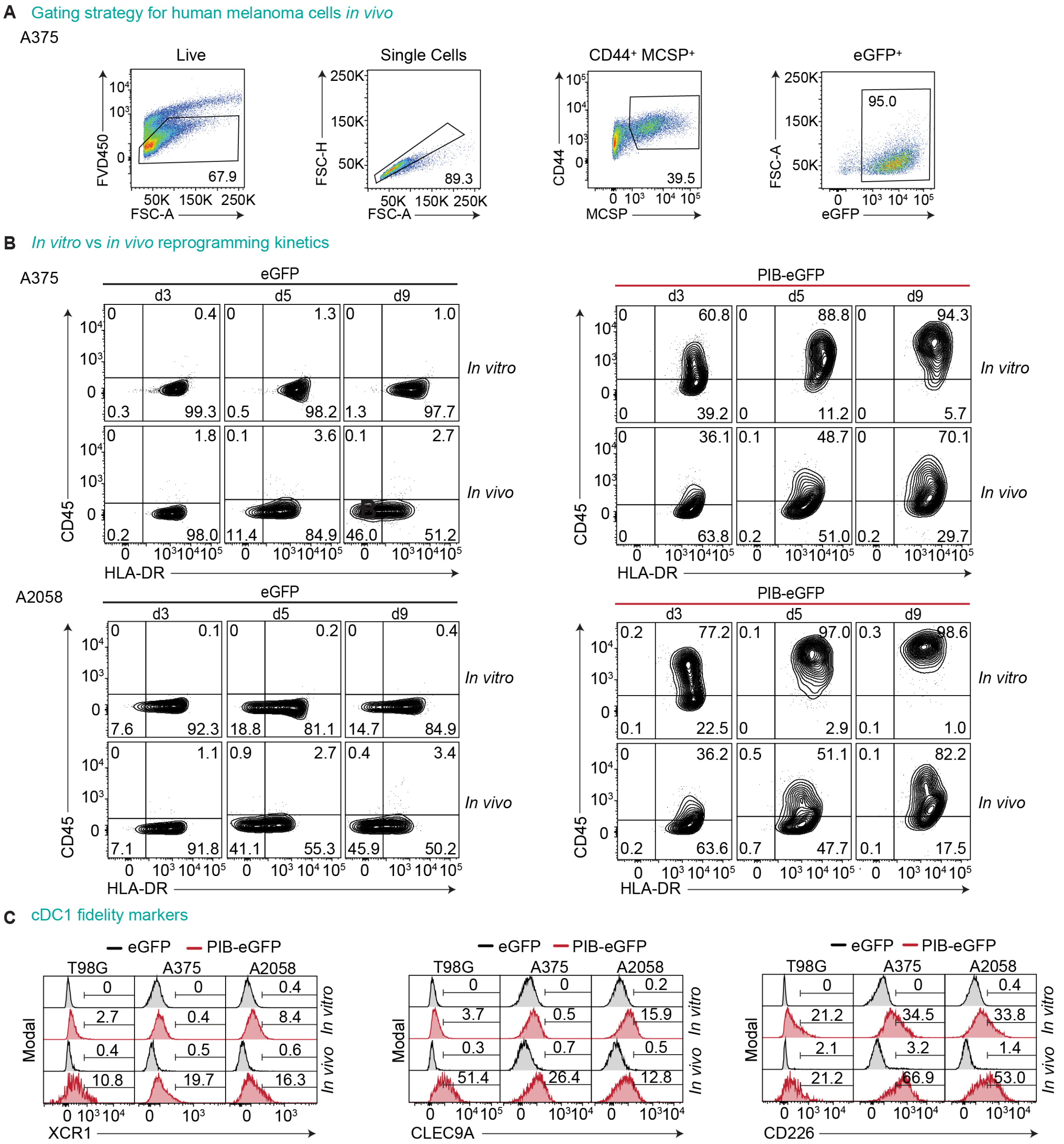


**Figure S7. PU.1, IRF8 and BATF3 induce dendritic cell reprogramming of human cancer cells *in vivo*.**

**(A-C)** Human glioblastoma T98G, melanoma A375 and A2058 cells were transduced with PIB-eGFP or eGFP *in vitro*, implanted subcutaneously into NSG mice and isolated at day 3, 5, and 9 for phenotypic profiling by flow cytometry. eGFP-transduced cells were used as controls and *in vitro* reprogrammed cells for comparison (n=3). (A) Gating strategy used for flow cytometry analysis of *in vivo* reprogramming in the human melanoma cell line A375. Reprogramming efficiency was evaluated by flow cytometry as the percentage of $CD45^{+}$ $HLA\text{-}DR^{+}$ cells (completely reprogrammed) and $CD45^{+}$ $HLA\text{-}DR^{-}$ or $CD45^{-}$ $HLA\text{-}DR^{+}$ cells (partially reprogrammed) gated in MCSP$^+$ CD44$^+$ eGFP$^+$ transduced A375 cells. (B) Representative flow cytometry plots showing *in vitro* and *in vivo* reprogramming kinetics of human A375 and A2058 cells at day 3, 5 and 9 within live CD44$^+$ MCSP$^+$ eGFP$^+$ cells. (C) Histograms of cDC1 fidelity markers XCR1 (left), CLEC9A (middle) and CD226 (right) expression in live CD44$^+$ MCSP$^+$ eGFP$^+$ A375 and A2058 melanoma cells or CD44$^+$ eGFP$^+$ T98G cells. The percentages of positive cells for each marker are highlighted.

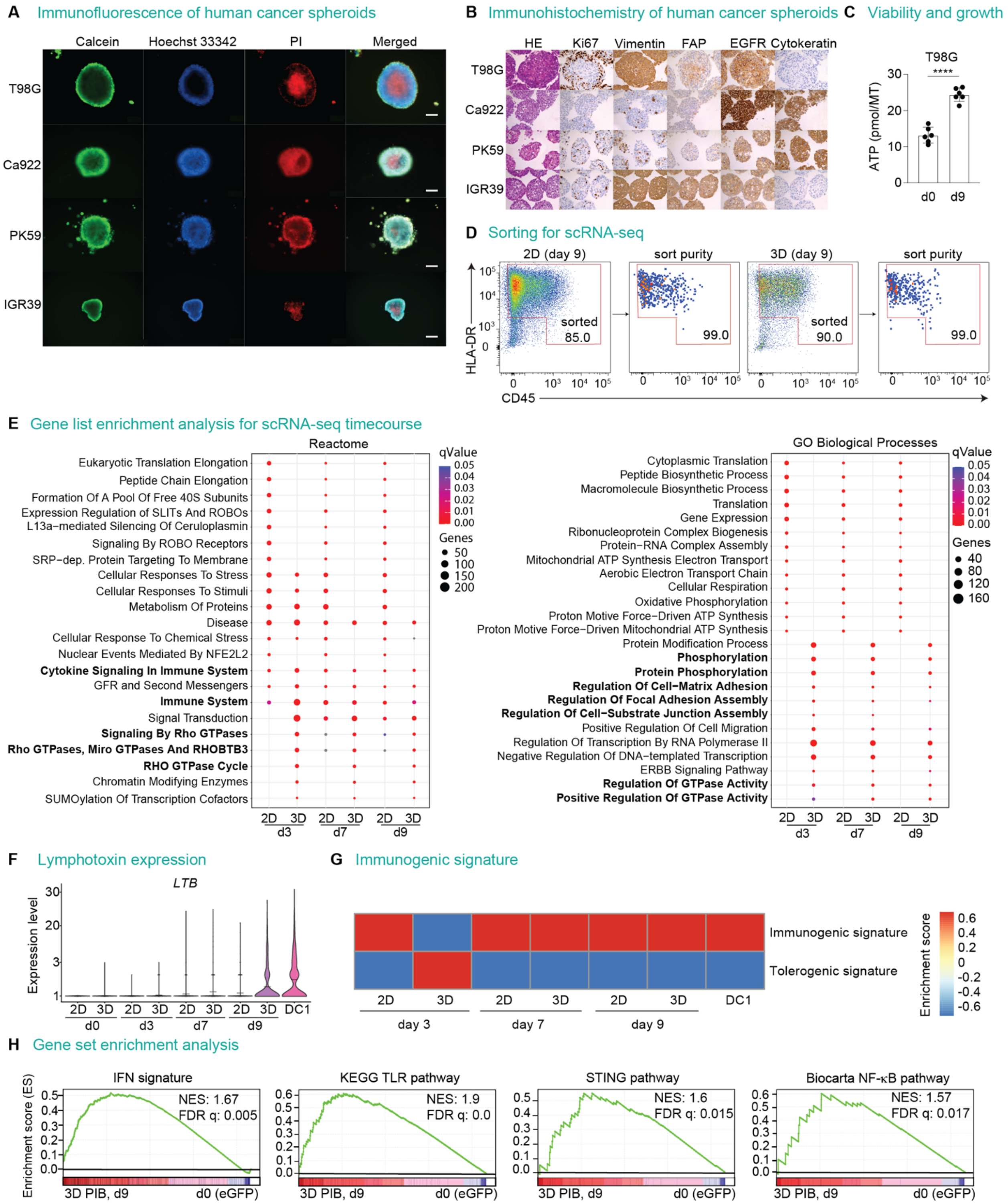


**Figure S8. cDC1 reprogramming progresses in human cancer spheroids.**
**(A-C)** Spheroids from human cancer cell lines were formed using the forced-floating method. (A) Representative fluorescent microscopy images show the spheroid architecture with live cells in the outer rim (Calcein AM$^+$ cells, green) and a necrotic core (PI$^+$ cells, red). Hoechst 33342 (blue) marks nuclear staining. Scale bar is 200 µm. (B) Immunohistochemistry profiling of spheroids

derived from human cancer cell lines stained for Hematoxylin and Eosin, Ki67, Vimentin, FAP, EGFR and Cytokeratin. Scale bar is 200 µm. (C) Spheroid growth over 9 days measured by the detection of ATP from lysed T98G-derived spheroids (n=6). **(D-H)** Before spheroid formation or 2D culture, T98G cells were transduced with lentiviral particles encoding PIB-eGFP or eGFP. $CD45^{+}HLA\text{-}DR^{+}$ cells (completely reprogrammed) and $CD45^{+}HLA\text{-}DR^{-}$ or $CD45^{-}HLA\text{-}DR^{+}$ cells (partially reprogrammed) were FACS-purified from single cell suspensions derived from 2D and 3D cultures at day 3, 7 and 9 and profiled by scRNA-seq. eGFP-transduced cells were included as controls and donor peripheral blood DC1 served as a reference. (D) Gating strategy for sorting of completely and partially reprogrammed cells (red) from 2D and 3D cultures by FACS. The purity of sorted populations is indicated. (E) Gene list enrichment analysis for Reactome pathways and GO biological processes for genes differentially expressed during reprogramming in 2D and 3D. Color gradient depicts adjusted p-values (qValue). The size of circles refers to the number of genes. (F) Violin plots showing mRNA expression of the Lymphotoxin Beta gene (*LTB*) along the reprogramming time course in 2D and 3D in cDC1-affiliated cells or all cells at day 0 and day 3 in 2D. (G) Gene set enrichment analysis for immunogenic signature (TLR-induced maturation) and tolerogenic signature (homeostatic maturation) (*32*) during reprogramming in 2D and 3D. Color gradient depicts enrichment score. (H) Gene set enrichment analysis for interferon (IFN) signature, KEGG Toll-like Receptor (TLR) signaling, STING and NF-κB signaling pathways in T98G cells at day 9 of reprogramming in 3D (3D PIB, d9) compared to eGFP-transduced cells (d0). Data in panel C are shown as mean ± SD and were compared using the t-test. ****$p<0.0001$.

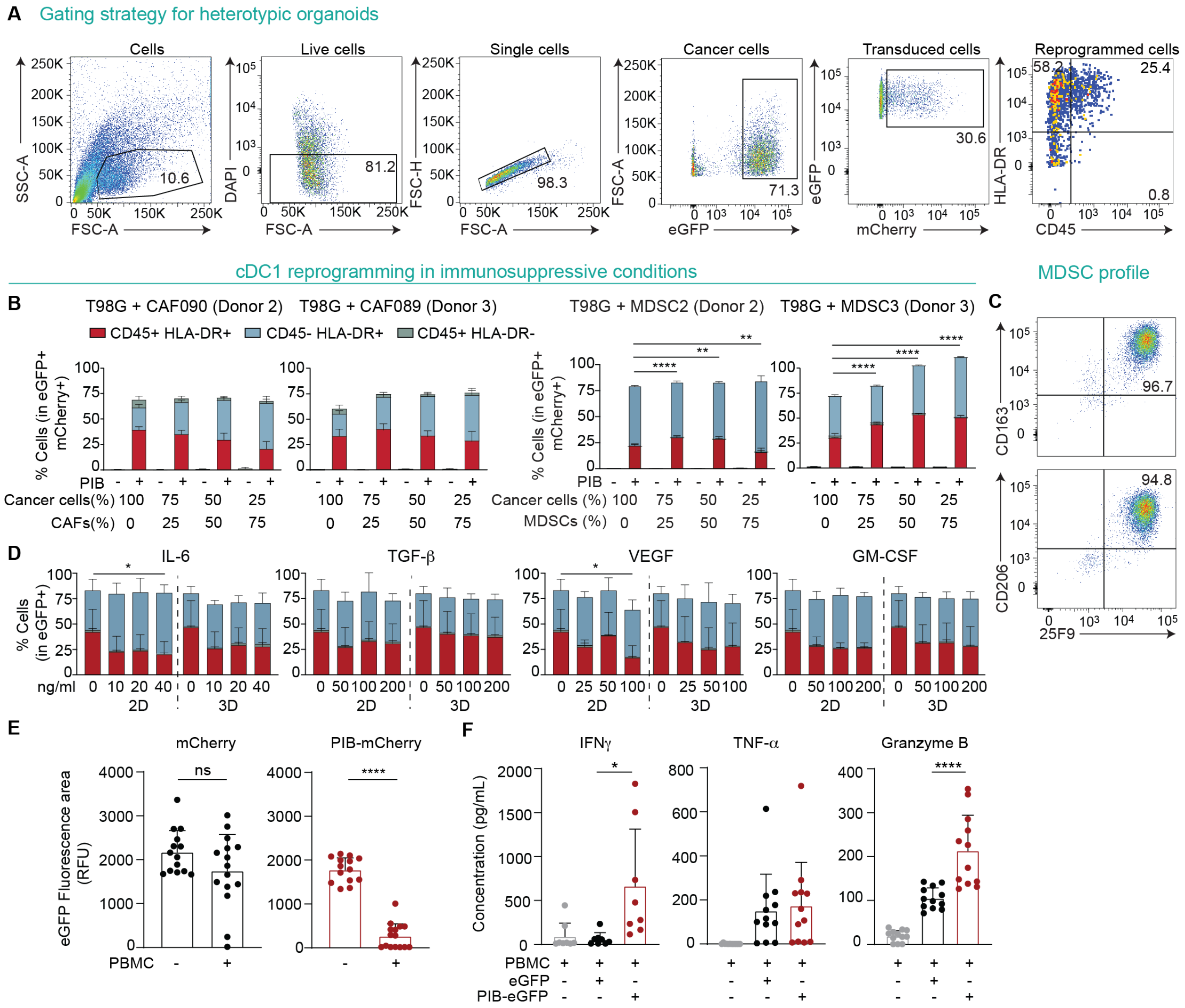


**Figure S9. Immunosuppressive cells do not impair cDC1 reprogramming.**

**(A-B)** Multi-component heterotypic spheroids were formed from eGFP$^{+}$ expressing T98G cells (T98G-eGFP$^{+}$) after transduction with PIB-mCherry or mCherry lentiviral vectors and combined with immunosuppressive cancer-associated fibroblasts (CAFs) or myeloid-derived suppressor cells (MDSC) at different ratios (indicated in x-axis). CAF090, CAF089, MDSC2, and MDSC3 refer to cells from different donors. (A) Gating strategy to evaluate reprogramming efficiency in T98G-eGFP$^{+}$ cells in heterotypic spheroids. Reprogramming efficiency was measured by CD45 and HLA-DR expression within transduced cancer cells (eGFP$^{+}$ mCherry$^{+}$) at day 9 by flow cytometry. (B) Reprogramming efficiency in heterotypic spheroids established with increasing proportions of CAFs (n=3-9) and MDSCs (n=3-4). **(C)** Representative plots showing flow cytometry analysis of the macrophage markers CD163, CD206, and 25F9 in MDSCs. **(D)** T98G cells were transduced with PIB-eGFP or eGFP and maintained either in 2D or used for spheroid formation. 2D and 3D cultures were reprogrammed for 9 days in the presence of increasing concentrations of immunosuppressive cytokines IL-6, TGF-β, VEGF, and immuno-regulatory GM-CSF (indicated in x-axis), and reprogramming efficiency was evaluated at day 9 (n=7-11).

Data in panel B and D are shown as mean ± SD. **(E)** Spheroid sizes as a measure of T cell cytoxicity against T98G-eGFP$^+$ containing CAFs after 7 days of co-culture with non-activated, unmatched (HLA-A2-negative) PBMCs. Relative fluorescence units (RFU) were quantified by the eGFP$^+$ fluorescence area by imaging (n=14-15). **(F)** Quantification of cytokine release 24 hours after co-culture of non-activated, unmatched (HLA-A2-negative) PBMCs from three donors with T98G-eGFP$^+$ spheroids containing CAFs (n=8-12). Data in panel B, D, E and F are shown as mean ± SD. Comparisons in panel B and D were performed using a two-way ANOVA and F using one-way ANOVA followed by Tukey's multiple comparison test. Data in panel E were analyzed using the Mann-Whitney test. *p<0.05, **p<0.01, ****p<0.0001.

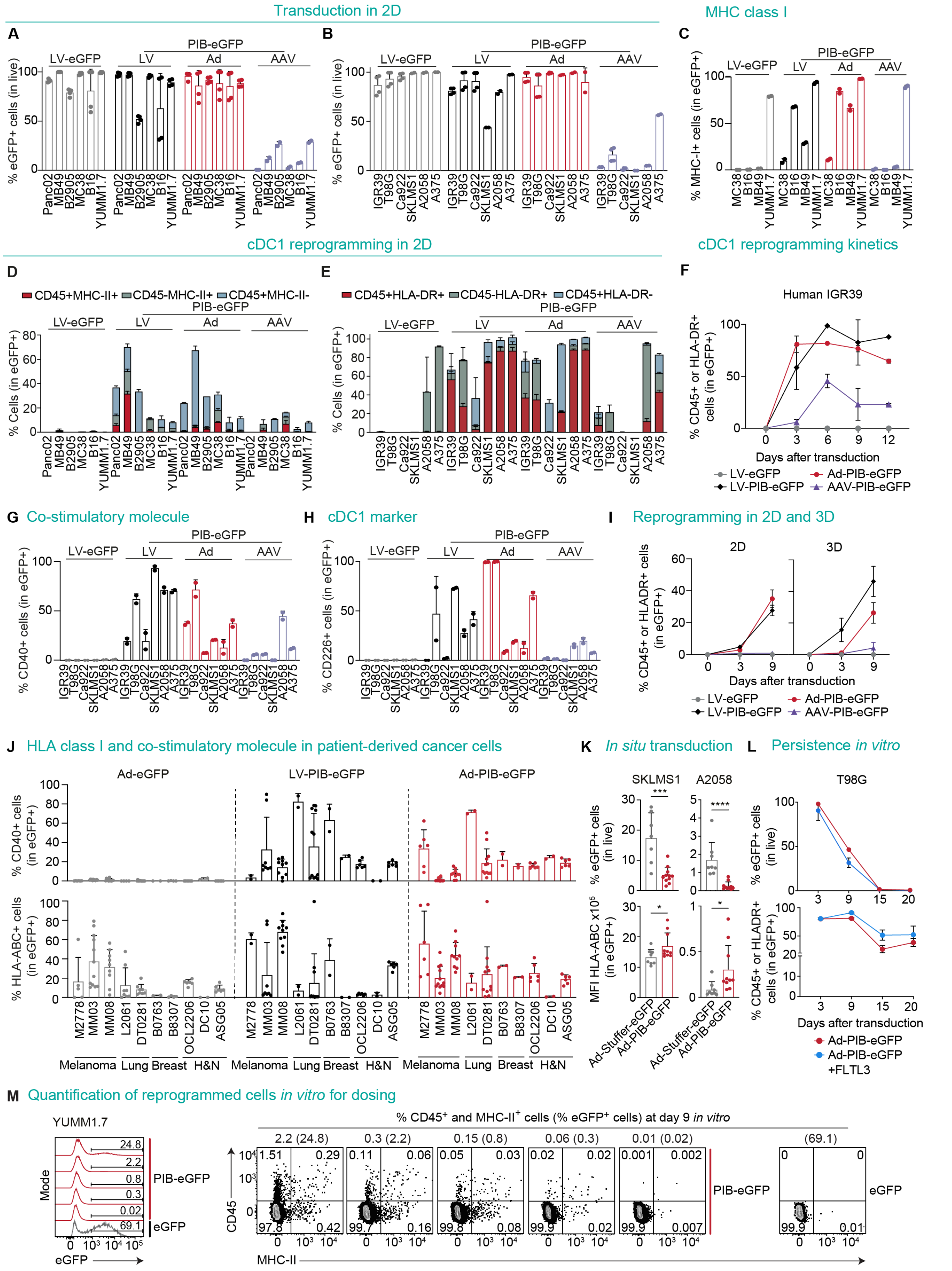


**Figure S10. Adenoviral vectors transduce and reprogram mouse and human cancer cells *in vitro* and *in vivo*.**

**(A-H)** Mouse and human cancer cell lines were transduced in 2D cultures with lentiviral (LV), adenoviral (Ad) or adeno-associated viral (AAV) vectors encoding PIB-eGFP. Transduction and reprogramming efficiencies were evaluated by flow cytometry. Lentiviral vectors encoding eGFP (LV-eGFP) were included as control. (A) Quantification of transduction efficiency by eGFP expression 3 days after transduction in murine cancer cells and (B) human cancer cell lines (n=2-4). (C) Percentages of MHC-I expression in mouse cancer cell lines 3 days after transduction (n=2). (D) Reprogramming efficiency in mouse cancer cell lines at day 3 and (E) human cancer cell lines at day 9 after transduction (n=4). (F) Reprogramming in human melanoma IGR39 measured by cumulative percentages of $CD45^+$ and HLA-$DR^+$ cells at days 0, 3, 6, 9, and 12 after transduction (n=2). (G) CD40 and (H) CD226 expression in human cancer cell lines at day 3 after transduction (n=2). **(I)** Reprogramming efficiency measured by the cumulative percentages of $CD45^+$ and HLA-$DR^+$ human T98G cancer cells maintained in 2D and 3D cultures 0, 3 and 9 days after transduction with LV, Ad, or AAV vectors (n=2). **(J)** CD40 and HLA-ABC expression in patient-derived cancer cells 9 days after transduction with LV, Ad, or AAV vectors in 2D (n=3). **(K)** *In situ* transduction efficiency measured by percentages of $eGFP^+$ cells (top) and mean fluorescence intensity (MFI) of MHC-I expression (bottom) in melanoma (A2058) and sarcoma (SKLSM1) cells after 4 intratumoral injections of Ad-PIB-eGFP or Ad-Stuffer-eGFP. Analysis was performed 9 days after the first intratumoral injection (n=8-10). (**L**) Persistence of adenoviral-mediated eGFP expression (upper) and reprogramming (lower) in T98G cells at days 3, 9, 15, and 20 after transduction with Ad-PIB-eGFP *in vitro.* Persistence of reprogrammed cells was measured as cumulative percentages of $CD45^+$ and HLA-$DR^+$ cells. When indicated cells were kept in culture with 100 ng/ml FLT3 ligand (FLT3L) (n=3). **(M)** YUMM1.7 cells were transduced with lentiviral vectors encoding PIB-eGFP or eGFP, serially diluted with untransduced cells to lower the amount of transduced and reprogrammed cells and injected subcutaneously into WT mice. Flow cytometry quantification of transduced cells by eGFP expression and reprogramming efficiency measured by $CD45^+$ and MHC-$II^+$ cells from *in vitro* cultures 9 days after transduction. Numbers indicate percentages of transduced $eGFP^+$ cells and cumulative percentages of reprogrammed $CD45^+$ and MHC-$II^+$ cells at day 9 gated within live cells. Data in panel A-L are shown as mean ± SD. Comparisons in panel K were analyzed using the Mann-Whitney test. *p<0.05, ***p<0.001, ****p<0.0001.

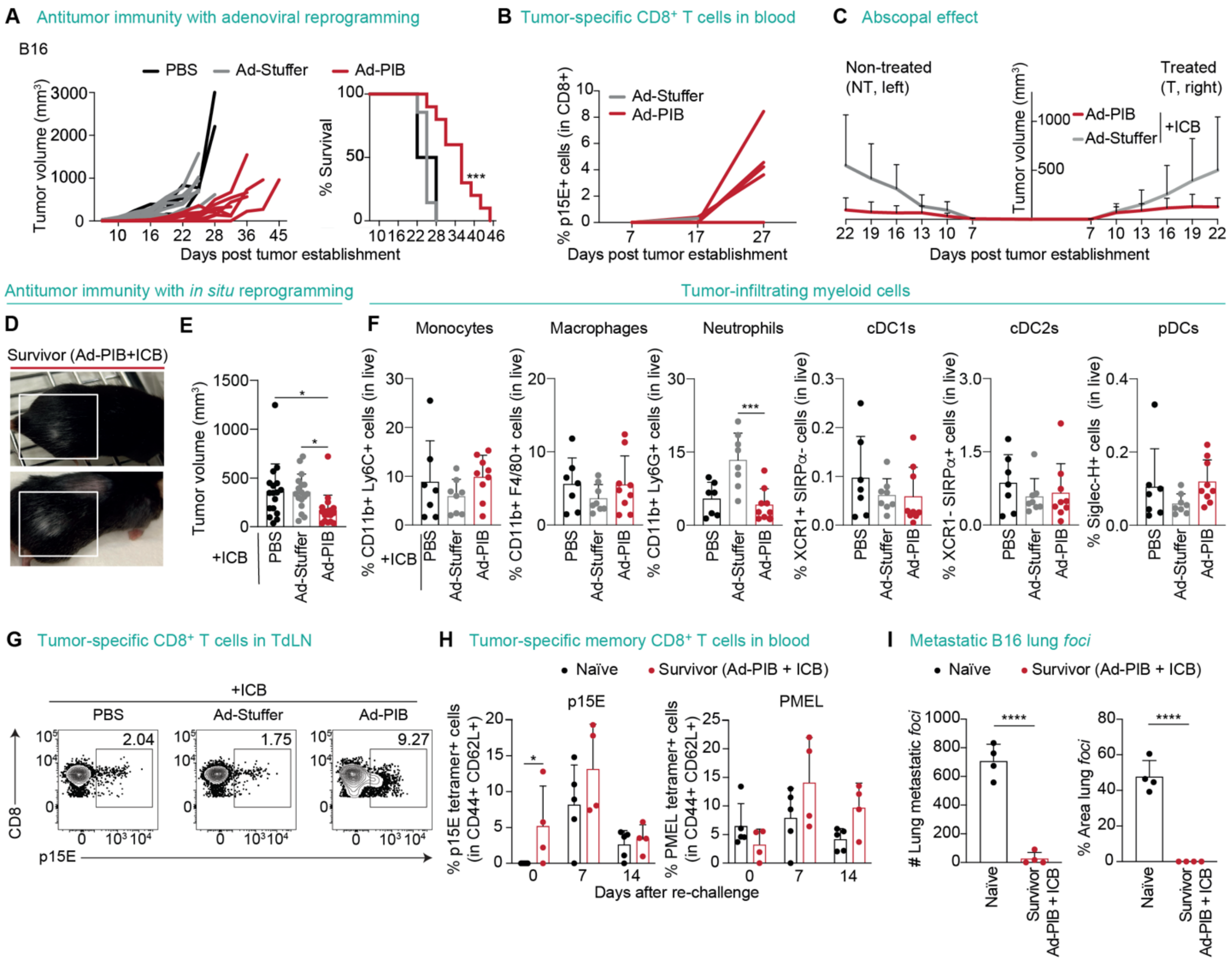


**Figure S11. Delivery of PIB intratumorally with adenoviral vectors elicits systemic and long-lasting antitumor memory.**

**(A, B)** B16 cells were transduced *in vitro* with Ad-PIB, mixed at 1:1 ratio with parental cells and injected subcutaneously to induce tumor establishment and *in vivo* reprogramming within the tumor. Untransduced B16 cells or B16 cells transduced with Ad-Stuffer (mixed with parental cells 1:1) were included as controls (n=3-7). (A) Tumor growth (left) and survival (right) of animals after tumor establishment. (B) Flow cytometry quantification of tumor antigen p15E-specific $CD8^+$ T cells in peripheral blood by tetramer staining at day 7, 17 and 27. **(C)** Tumor growth of bilateral B16 tumors established by subcutaneous injection of $2x10^5$ Ad-PIB- or Ad-Stuffer-transduced B16 cells mixed 1:1 with parental cells on the right flank and $1x10^5$ untransduced B16 cells on the left flank. ICB (anti-PD-1 and anti-CTLA-4) were administered intraperitoneally at day 7, 10, and 13 post tumor establishment (n=7-9). **(D-G)** B16 tumors were established subcutaneously and injected intratumorally at day 7, 9, 11, and 13 with Ad-PIB gene therapy. Intratumoral injections with Ad-Stuffer or PBS were included as controls. ICB (anti-PD-1 and anti-CTLA-4) were administered intraperitoneally at day 7, 10, and 13 (n=7-9). (D) Representative animals treated with Ad-PIB in combination with ICBs showing vitiligo. (E) Tumor volume of B16 tumors used for immunophenotyping 9 days after first intratumoral injection. (F) Flow cytometry quantification of tumor-infiltrating myeloid cells 9 days after first intratumoral injection. (G) Flow cytometry analysis of tumor antigen p15E-specific $CD8^+$ T cells in tumor-draining lymph nodes at day 16. **(H)** Ad-PIB-treated mice with complete tumor regression were re-challenged at day 100 via subcutaneous injection of $1x10^5$ parental B16 cells. Treatment-naïve age-matched mice were included as controls. Flow cytometry quantification of p15E (left) and PMEL (right) $tetramer^+$ memory $CD44^+CD62L^+CD8+$ T cells from peripheral blood at day 0, 7 and 14 after subcutaneous re-challenge (n=4). **(I)** Survivor Ad-PIB-treated mice were re-challenged intravenously with $1x10^5$ B16 cells 160 days after initial tumor establishment. Treatment-naïve age-matched mice were included as controls. Quantification of lung metastatic *foci* 21 days after intravenous challenge (n=4). Data in panel C, E, F, H, and I are shown as mean ± SD. Survival analysis in panel A was performed by log-rank Mantel-Cox test. Comparisons in panel E, F, H, and I were analyzed using the Mann-Whitney test. *p<0.05, ***p<0.001, ****p<0.0001.

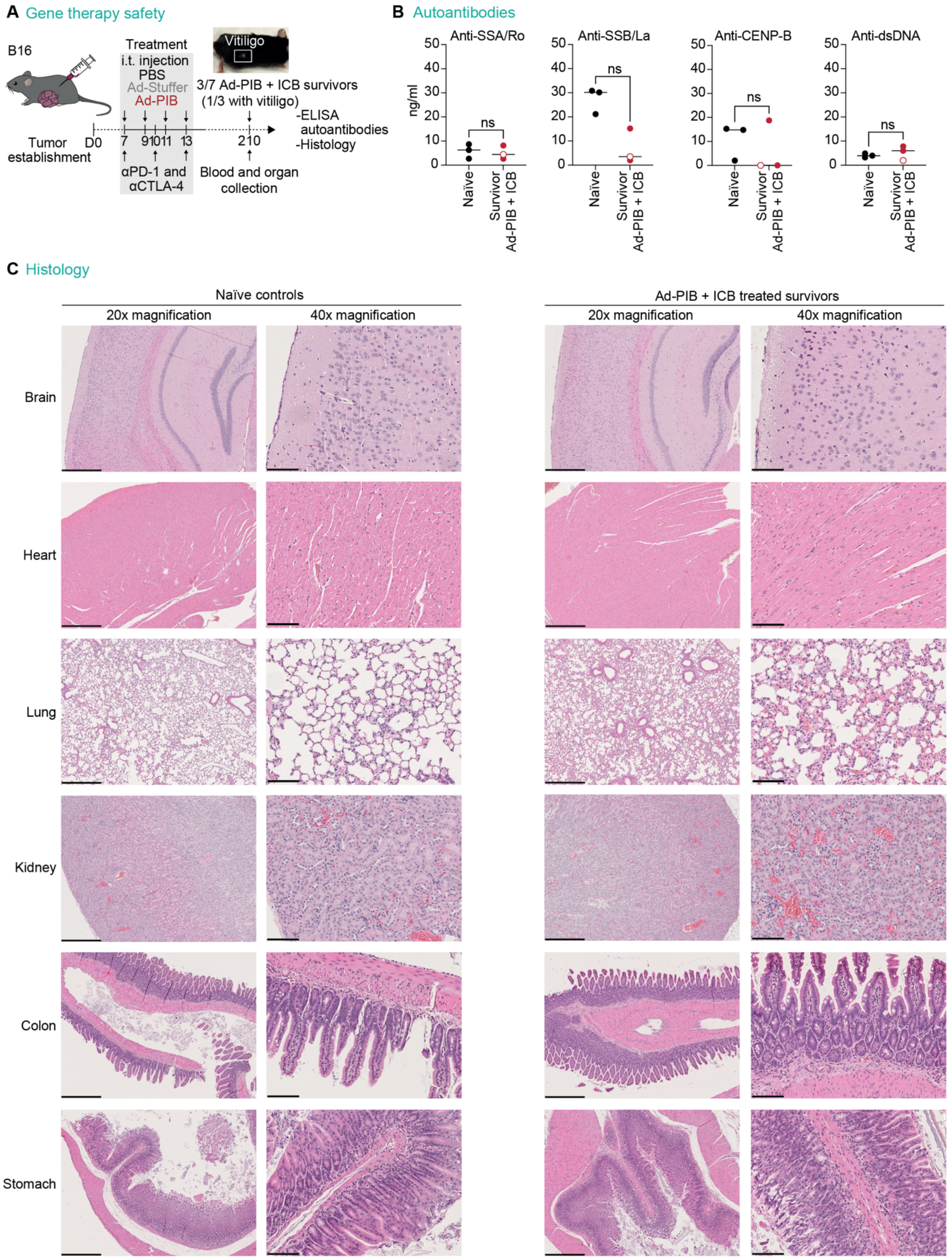


**Figure S12. Adenoviral delivery of PIB elicits long-term survival without off-target toxicities.** **(A)** Experimental design assessing long-term survival and safety with Ad-PIB gene therapy in

C57BL/6J mice with subcutaneous B16 melanoma tumors. Tumors were injected 4 times with Ad-PIB, control vector (Ad-Stuffer) or PBS at day 7, 9, 11 and 13 after tumor establishment. Anti-PD-1 and anti-CTLA-4 (ICB) were administered intraperitoneally at day 7, 10 and 13. At day 210, plasma and organs including brain, heart, lung, kidney, colon, and stomach were collected from 3 survivor mice including 1 mouse with vitiligo and 3 naïve control mice. Plasma was used to quantify autoantibodies and organs were collected to assess immune infiltration and off-target toxicities. (**B**) Quantification of autoantibodies against double-stranded DNA (dsDNA), SSA antigen (SSA/Ro), SSB antigen (SSB/La) or Centromere protein-B (CENP-B) in the plasma of survivor mice treated with Ad-PIB + ICB and naïve mice by ELISA (n=3). The open circle marks Ad-PIB survivor mouse with vitiligo. (**C**) Histology analysis score measuring off-target toxicities for brain, heart, kidney, colon, stomach (naive mice: 0/300; Ad-PIB + ICB: 0/300) and lung (naïve: 2/300; Ad-PIB + ICB: 12/300). Representative H&E-stained tissue sections are shown. Scale bars are 500 μm (left) and 100 μm (right). Data in panel B are shown as mean ± SD and comparisons were analyzed using the Mann-Whitney test. ns - non-significant.
